# Mitochondria-containing Extracellular Vesicles (EV) Reduce Mouse Brain Infarct Sizes and EV/HSP27 Protect Ischemic Brain Endothelial Cultures

**DOI:** 10.1101/2021.10.29.466491

**Authors:** Kandarp M. Dave, Donna B. Stolz, Venugopal R. Venna, Victoria A. Quaicoe, Michael E. Maniskas, Michael John Reynolds, Riyan Babidhan, Duncan X. Dobbins, Maura N. Farinelli, Abigail Sullivan, Tarun N. Bhatia, Hannah Yankello, Rohan Reddy, Younsoo Bae, Rehana K. Leak, Sruti S. Shiva, Louise D. McCullough, Devika S Manickam

**Affiliations:** Graduate School of Pharmaceutical Sciences, Duquesne University, Pittsburgh, PA; Center for Biologic Imaging, University of Pittsburgh Medical School, Pittsburgh, PA; Department of Neurology, McGovern Medical School, University of Texas Health Science Center, Houston, TX; Pittsburgh Heart Lung Blood Vascular Institute, University of Pittsburgh Medical School, Pittsburgh, PA; Department of Biochemistry and Molecular Biology, Gettysburg College, Gettysburg, PA; Psychological and Brain Sciences, Villanova University, Villanova, PA, USA; Departments of Chemical and Biomedical Engineering, Carnegie Mellon University, Pittsburgh, PA; Department of Pharmaceutical Sciences, College of Pharmacy, The University of Kentucky, Lexington, KY; Department of Pharmacology & Chemical Biology, University of Pittsburgh Medical School, Pittsburgh, PA

**Author notes:** Corresponding author: Devika S Manickam, Ph.D., 600 Forbes Avenue, 453 Mellon Hall, Pittsburgh, PA 15282., Twitter: @manickam_lab.

**Keywords:** extracellular vesicles, mitochondria, heat shock protein, paracellular permeability, BBB protection, ischemic stroke

## Abstract

Ischemic stroke causes brain endothelial cell (BEC) death and damages tight junction integrity of the blood-brain barrier (BBB). We harnessed the innate mitochondrial load of endothelial cell-derived extracellular vesicles (EVs) and utilized mixtures of EV/exogenous heat shock protein 27 (HSP27) as a one-two punch strategy to increase BEC survival (via EV mitochondria) and preserve their tight junction integrity (via HSP27 effects). We demonstrated that the medium-to-large (m/lEV) but not small EVs (sEV) transferred their mitochondrial load, which subsequently colocalized with the mitochondrial network of the recipient primary human BECs. BECs treated with m/lEVs increased relative ATP levels and displayed superior mitochondrial function. Importantly, m/lEVs isolated from oligomycin (mitochondrial complex V inhibitor) or rotenone (mitochondrial complex I inhibitor)-exposed BECs (RTN-m/lEVs or OGM-m/lEVs) did not increase BECs ATP levels compared to naïve m/lEVs. In contrast, RTN-sEV and OGM-sEV functionality in increasing cellular ATP levels was minimally impacted in comparison to naïve sEVs. Intravenously administered m/lEVs showed a reduction in brain infarct sizes compared to vehicle-injected mice in a mouse middle cerebral artery occlusion model of ischemic stroke. We formulated binary mixtures of human recombinant HSP27 protein with EVs: EV/HSP27 and ternary mixtures of HSP27 and EV with cationic polymer poly (ethylene glycol)-b-poly (diethyltriamine): (PEG-DET/HSP27)/EV. (PEG-DET/HSP27)/EV and EV/HSP27 mixtures decreased the paracellular permeability of small and large molecular mass fluorescent tracers in oxygen glucose-deprived primary human BECs. This one-two-punch approach to increase BEC metabolic function and tight junction integrity is a promising strategy for BBB protection and prevention of long-term neurological dysfunction post-ischemic stroke.

**Graphical Abstract:** 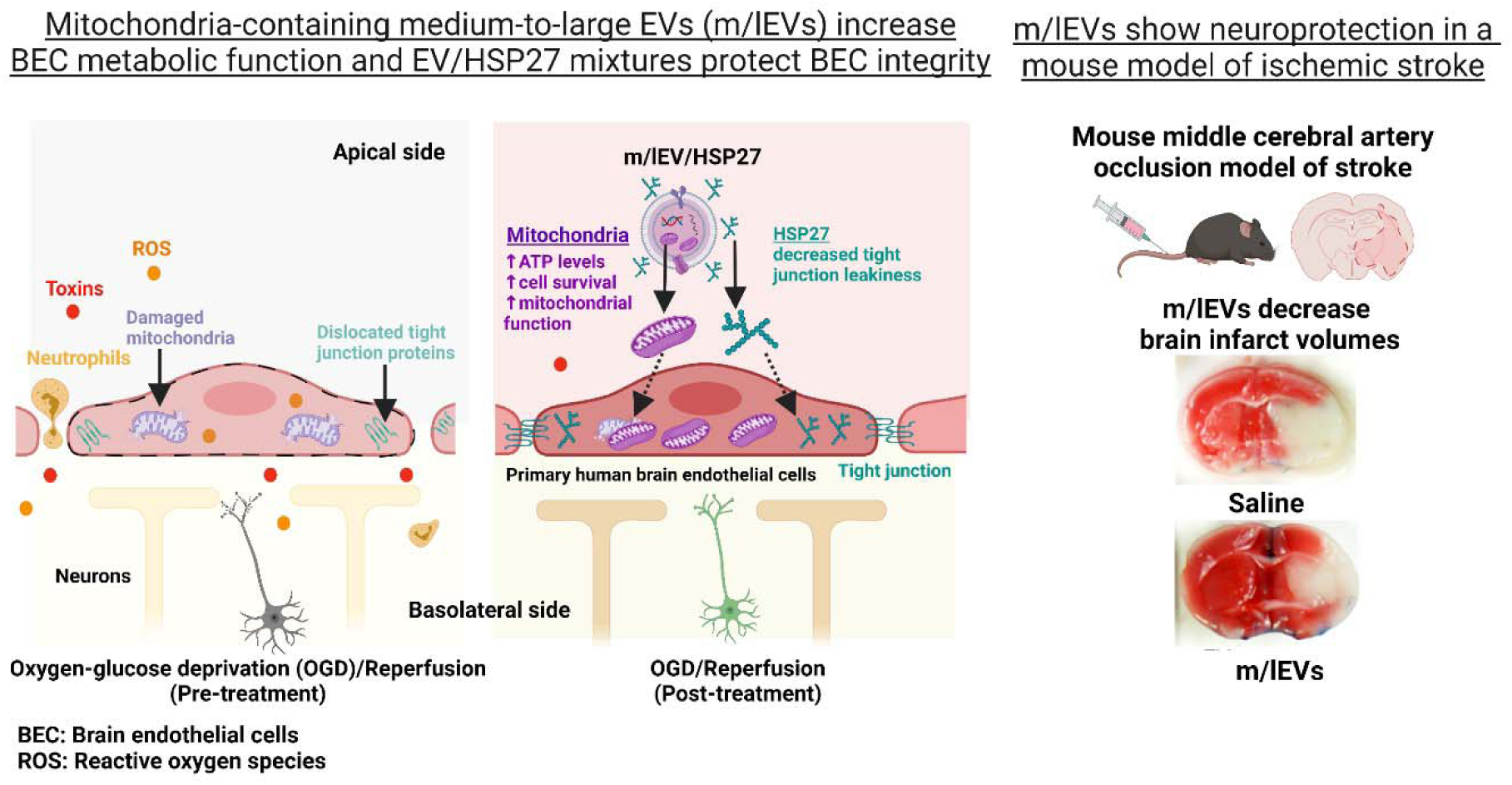

**Highlights:**

- Medium-to-large extracellular vesicles (m/lEVs), not small EVs contain mitochondria
- m/lEVs increased ATP and mitochondrial function in brain endothelial cells (BECs)
- m/lEVs from oligomycin-exposed BECs did not increase recipient BEC ATP levels
- Intravenously injected m/lEVs reduced brain infarct sizes in a mouse stroke model
- EV/HSP27 mixtures reduced small and large dextran molecule permeability across BECs

## 1. Introduction

EVs are an emerging class of natural carriers for drug delivery due to their known roles in intercellular communication. They retain membrane signatures reminiscent of the donor cells from which they are derived, and therefore possess inherent homing capabilities to recipient cells of the same type ^1, 2^. They are also likely to be less immunogenic compared to synthetic nanoparticles. The smaller EVs (sEVs) range from 30-200 nm in particle diameter and their biogenesis involves the inward budding of endosomal membranes that transforms into multivesicular bodies followed by their fusion with the plasma membrane and sEV release into extracellular spaces ^2^. The biogenesis of the medium-to-larger EVs (m/lEVs) involves their outward budding from the cell’s plasma membrane with particle diameters ranging from 100 – 1000 nm ^2^. The selective packaging of the functional mitochondria and mitochondrial proteins in the m/lEVs motivated us to harness the m/lEV mitochondrial load as a therapeutic modality. Mitochondria play a central role in cellular energy production and regulation of cell death including apoptosis and autophagy ^3^. Ischemia-induced mitochondrial dysfunction in the brain endothelial cells (BECs) lining the blood-brain barrier (BBB) initiates the generation of excessive reactive oxygen species, reduction in ATP levels, and consequently BEC death ^3^. Therefore, protection of mitochondrial function via exogenous mitochondria supplementation is a potent strategy to increase BEC survival post-ischemic stroke. Thus, we rationalize that mitochondria-containing m/lEVs derived from BECs can increase cellular bioenergetics and survival under hypoxic conditions.

Ischemic stroke-induced oxygen-glucose-deprivation (OGD) decreases ATP levels in the BECs leading to the accumulation of cellular cations and excitatory neurotransmitters ^4–6^. The cationic overload catalyzes enzymatic activities leading to generation of reactive oxygen species (ROS), impairment of mitochondrial ROS defense mechanisms, and the subsequent mitochondrial dysfunction. Therefore, restoring mitochondrial function is a viable strategy to reduce damage to the BECs. In addition, disruption of the BBB is a major hallmark of ischemic stroke that is associated with altered expression of tight junctions, adherens junction proteins, and BBB transporters ^4, 5, 7^. Early ischemia/reperfusion activates the polymerization of the actin cytoskeleton in endothelial cells which disassembles and internalizes the tight junction proteins and consequently lead to the loss of barrier properties of the BECs lining the BBB ^8, 9^. Uncontrolled actin polymerization-induced breakdown of BBB leads to the infiltration of proinflammatory mediators, blood cells, circulatory immune cells, and toxins into the brain parenchyma, and leads to the secondary injury cascade ^9, 10^. Hence, a combined strategy to decrease BEC death and their paracellular permeability, ultimately leading to protection of the BBB metabolic function and tight junction integrity is a potent approach for the treatment of ischemia/reperfusion injury.

Preclinical studies have demonstrated that endothelial, but not neuronal overexpression of heat shock protein 27 (HSP27), inhibited actin polymerization and elicited long-lasting protection against stroke-induced BBB disruption and neurological deficits ^9, 10^. HSP27 binds to actin monomer and inhibits tight junctional protein translocation in endothelial cells ^9, 11–14^. Intravenous administration of cell-penetrating transduction domain (TAT)-HSP27 rapidly enhanced HSP27 levels in brain microvessels, decreased infarct volumes, and attenuated ischemia/reperfusion-induced BBB disruption ^9^. Therefore, we selected human recombinant HSP27 protein as a model therapeutic protein to determine whether EV/HSP27 mixtures may decrease tight junction permeability post-ischemia/reperfusion in culture conditions.

Administration of EV/exogenous HSP27 mixtures is a promising strategy that can allow harnessing the inherent targeting capabilities of the EVs to the recipient BECs along with the added benefits of their mitochondrial load. Here, we *hypothesize* that innate mitochondria-containing BEC-derived EVs with exogenous HSP27 protein is a one-two-punch approach to increase BEC survival and protect its tight junction barrier via decreasing the paracellular permeability post-ischemia. This approach will protect and strengthen the BBB that in turn can ameliorate long-term neurological damage and dysfunction. We tested the effects of adding a cationic copolymer, poly (ethylene glycol)-*b*-poly (diethyltriamine) (PEG-DET) to the EV/HSP27 mixtures to determine if the degree of HSP27 interactions can be further improved. We have previously used PEG-DET polymer to form nanosized mixtures with superoxide dismutase protein and demonstrated a >50% reduction in brain infarct volume in a mouse model of acute ischemic stroke ^15^. PEG-DET is a cationic diblock copolymer known for its safety and gene transfer efficacy in comparison to commercial transfection agents such as lipofectamine, polyethyleneimine, and other cationic polymers ^16–18^.

In this study, we isolated sEVs and m/lEVs from hCMEC/D3: a human brain endothelial cell line using a sequential ultracentrifugation method ^19^ and characterized their particle diameter, zeta potential, and membrane integrity post-cold storage. According to recommendations from the 2018 Minimal Information for Studies of Extracellular Vesicles ^20^, EVs with an average particle diameter <200 nm should be referred to as small EVs (sEVs), and >200 nm as medium-to-large EVs (m/lEVs). In our studies, the average particle diameter of EVs that pelleted down at 120,000 ×*g* was about 122 nm, whereas the average diameter of EVs isolated at 20,000 ×*g* was about 185 nm. Therefore, we refer to EVs isolated at 120,000 *×g* as sEVs, and EVs isolated at 20,000 ×*g* as m/lEVs. A mixture of sEV and m/lEVs at a 1:1 weight: weight ratio is referred to as EVs. While we collectively refer to both large (m/lEV) and small (sEV) vesicle fractions as EVs, we have studied the singular effects of both m/lEVs and sEVs. We showed the presence of mitochondrial components in m/lEVs using transmission electron microscopy and western blotting. We evaluated the effects of EV dose and incubation times on their uptake to the recipient BECs and demonstrated the colocalization of m/lEV-delivered mitochondria with the mitochondrial network of the recipient BECs. We studied the effects of EV exposure on the resulting relative ATP levels, mitochondrial respiration, and glycolytic capacity of the recipient BECs. We conducted a pilot experiment in a mouse middle cerebral artery occlusion model of stroke to determine its potential therapeutic effects and to determine if m/lEV treatment is safe from any adverse effects when administered intravenously (i.v.) to the mice. Twenty-four hours post-injection, we analyzed the brain infarct sizes of mice treated with m/lEVs or saline to determine its therapeutic effects. The physicochemical characteristics of EV/HSP27 binary mixtures and (PEG-DET/HSP27)/EV ternary mixtures were studied using native polyacrylamide gel electrophoresis and dynamic light scattering. The effects of EV/HSP27 on the paracellular permeability of small and large molecule fluorescent tracers were evaluated under ischemic and ischemia/reperfusion conditions in primary human BECs.

## 2. Materials

Recombinant human HSP27 was purchased from Novus Biologicals (Centennial, CO). Cell Titer Glo 2.0 reagent (ATP assay) was procured from Promega (Madison, WI). Micro BCA and Pierce BCA protein assay kits were purchased from Thermo Scientific (Rockford, IL). PEG-DET polymer was synthesized by aminolysis of PEG-poly(β-benzyl L-aspartate) block copolymers with diethyltriamine as previously reported ^16, 18, 21, 22^. Bio-Safe Coomassie G-250 stain was purchased from Bio-Rad Laboratories Inc. (Hercules, CA). Collagen Type I was purchased from Corning (Discovery Labware Inc, Bedford, MA) and endothelial cell basal medium-2 (EBM-2) was procured from Lonza (Walkersville, MD). Hydrocortisone, human basic fibroblast growth factor, ascorbic acid, and rotenone and oligomycin A were purchased from Sigma-Aldrich (Saint Louis, MO). The penicillin-Streptomycin solution, Chemically Defined Lipid Concentrate, and calcein-AM were procured from Invitrogen (Carlsbad, CA). Polycarbonate centrifuge tubes were purchased from Beckman Coulter, Inc. (Brea, CA). The electrophoresis sample buffer was purchased from Bio-Rad (Hercules, CA). PET track-etched membrane Falcon Cell Culture inserts of 0.4 µm pore size were procured from Corning (Discovery Labware Inc, Bedford, MA). TRITC 65-85 kD and 4.4 kD dextran was procured from Sigma (St. Louis, MO). A low-volume disposable cuvette (Part no. ZEN0040, Malvern) was used for particle size measurements. CellLight Mitochondria-GFP Backman 2.0 reagent, MitoTracker Green FM, and MitoTracker Deep Red FM, Dynabeads Protein G (cat#10003D), and DynaMag-2 magnetic stand (cat#12321D) were procured from Invitrogen (Eugene, OR). Mouse monoclonal antibodies against ATP5A (cat#ab14748), GAPDH (cat#ab8245), HSP27 (cat#ab2790) were purchased from Abcam. Mouse monoclonal antibody against CD9 (cat#ab92726) was received from Life Technologies Corporation (Eugene, OR), whereas Alexa Fluor 790-conjugated donkey anti-mouse IgG was received from Jackson ImmunoResearch Lab Inc (West Grove, PA).

### 2.1. Cell models

A human cerebral microvascular endothelial cell line (hCMEC/D3, cat#102114.3C) was received from Cedarlane Laboratories (Burlington, Ontario) at passage number (P) 25, and cells between P25 and P35 were used in all experiments ^19, 23^. hCMEC/D3 cells were grown in tissue culture flasks, multiwell plates, or transwell inserts precoated using 0.15 mg/mL rat collagen I in a humidified 5% CO_2_ incubator at 37 ± 0.5°C (Isotemp, Thermo Fisher Scientific). The cells were cultured in complete growth medium composed of endothelial cell basal medium (EBM-2) supplemented with fetal bovine serum (5% FBS), penicillin (100 units/mL)-streptomycin (100 μg/mL) mixture, hydrocortisone (1.4 µM), ascorbic acid (5 µg/mL), Chemically Defined Lipid Concentrate (0.01%), 10 mM HEPES (pH 7.4), and bFGF (1 ng/mL). The complete growth medium was replenished every other day until the cells formed confluent monolayers. Prior to passage, the cells were washed using 1x phosphate buffer saline (PBS) and detached from the flask using 1x TrypLE Express Enzyme (Gibco, Denmark). We received primary human brain microvascular endothelial cells (HBMEC, catalog no. ACBRI 376) from Cell Systems (Manassas, VA) at P3, and cells below P11 were used in all experiments. HBMECs maintained in Cell Systems Classic Culture Medium containing 1% culture boost were cultured in tissue culture flasks, multiwell plates, or transwell inserts pre-treated with attachment factors. The complete growth medium was replenished every day until the cells formed confluent monolayers. For passage, HBMEC monolayers were washed with Passage Reagent Group 1 (PRG 1, dPBS/EDTA solution), detached with PRG 2 (dPBS/trypsin-EDTA solution) and PRG 3 (trypsin inhibitor solution).

### 2.2. Isolation of sEVs and m/lEVs from hCMEC/D3 cells

It should be noted that while we collectively refer to both large (m/lEVs) and small (sEVs) vesicle fractions as EVs, we study both m/lEVs and sEVs. Wherever noted, a 1:1 w/w mixture of sEVs and m/lEVs is collectively referred to as EVs. sEVs and m/lEVs were isolated from conditioned medium supernatants of hCMEC/D3 cells using the differential ultracentrifugation method ^19^. We used three tissue culture flasks with a surface area of 175 cm^2^ for culturing the hCMEC/D3 BECs for EV isolation. The average total number of cells in each flask was about 16-19×10^6^ cells. Tissue culture flasks with a 175 cm^2^ growth area (T175) containing confluent hCMEC/D3 were washed with pre-warmed 1x PBS pH 7.4 (0.0067 M, PO4) without calcium and magnesium (catalog#SH30256.01, HyClone Lab, S Logan, Utah), and incubated with serum-free medium for 48 h in a humidified 5% CO_2_ incubator at 37 ± 0.5°C. Post-incubation, the EV-conditioned medium was collected in polypropylene centrifuge tubes and centrifuged at 2000 ×*g* for 22 min at 4°C to pellet down apoptotic bodies and cell debris using a Sorvall ST 8R centrifuge (ThermoFisher Scientific, Osterode am Harz, Germany). The supernatant was transferred into polycarbonate tubes and centrifuged at 20,000 ×*g* for 45 min at 4°C to pellet down m/lEVs using an Optima XE-90 ultracentrifuge equipped with a 50.2 Ti rotor (Beckman Coulter, Indianapolis, IN). Next, the supernatant was filtered through a 0.22 µm PES membrane syringe filter, and the flow-through was centrifuged at 120,000 ×*g* for 70 min at 4°C to collect sEVs. Lastly, m/lEVs and sEVs pellets were washed once with 1x PBS and suspended in 1x PBS for particle diameter measurements and *in vitro* experiments or 10 mM HEPES buffer pH 7.4 for zeta potential measurements. sEVs and m/lEVs samples were stored at −20°C until further use. The total protein content in sEVs and m/lEVs was quantified using Pierce MicroBCA assay.

For isolation of mitochondria-depleted EVs, confluent hCMEC/D3 BECs were treated with 1 µM rotenone (mitochondrial complex I inhibitor) or oligomycin A (electron transport chain complex V inhibitor) in complete culture medium for 4 h. Post-treatment, the treatment mixture was replaced with serum-free culture medium and incubated in a humidified incubator at 37°C for 48 h. We isolated sEVs and m/lEVs from the conditioned medium of hCMEC/D3 BECs using a differential ultracentrifugation method. EVs isolated post rotenone (RTN) and oligomycin A (OGM)-treatment were referred to as RTN-sEVs, RTN-m/lEVs and OGM-sEVs, OGM-m/lEVs, respectively.

### 2.3. Dynamic light scattering (DLS)

The stability of naïve EVs under storage conditions was determined by their measuring particle diameter and zeta potential using dynamic light scattering. EV samples at 0.5 mg protein/mL in 1x PBS were frozen at −20°C for 24 h and thawed at room temperature for 30 min. Post-thawing, the particle diameters, and dispersity indices were measured using Malvern Zetasizer Pro (Worcestershire, UK). The freeze-thaw cycle was repeated three times and all samples were run in triplicate. Average particle diameter, dispersity index, and zeta potential values were reported as mean ± standard deviation.

### 2.4. Membrane integrity of EVs after isolation and upon storage conditions

The membrane integrity of sEV and m/lEV post-isolation and upon revival after their storage conditions (described earlier in the DLS section) was determined using a calcein AM flow cytometry assay. First, polystyrene sub-micron-sized beads ranging from 0.2 to 2 µm particle diameters were used to calibrate the Attune NxT flow cytometer. The calibration beads, sEV, and m/lEV samples were tested and events captured in forward scatter (FSC) and side scatter (SSC) plots were analyzed using a small particle side scatter 488/10-nm filter (BL1) channel. sEVs and m/lEVs were diluted to 20 µg protein/mL in PBS and incubated/stained with 10 µM calcein AM for 20 min at room temperature in the dark. Unstained sEVs and m/lEVs were used to gate the background signals, whereas samples treated with 2% v/v Triton X-100 followed by staining with calcein AM were used as controls to determine if calcein-positive signals were specifically associated with intact EVs. For each sample analysis, an aliquot of 100 µL was run through Attune NxT Acoustic cytometer (Invitrogen, Singapore) and 50,000 events were recorded during the run. The calcein-associated fluorescence intensity was detected at 488/10 nm and percentage signal intensities were presented in histogram plots generated using Attune software. Calcein AM-associated background signals were gated using the controls such as PBS containing 10 µM calcein AM and PBS/2% Triton X-100/calcein AM mixture.

### 2.5. Transmission Electron Microscopy (TEM) analysis of EVs

*Negative-stain images of EVs:* Suspensions (150 µL) of EVs were pelleted at 100,000 ×g in a Beckman Airfuge for 45 min, the supernatant was carefully removed, and the pellets gently resuspended in 30 µL of PBS. Formvar coated, 100 mesh grids were floated onto 10 µL of this suspension and allowed to incubate for 5 min. The solution was wicked away with Whatman filter paper and stained with 0.45 mm filtered, 1% uranyl acetate, immediately wicked away with the filter paper. EV images were then imaged on a JEM-1400 Flash transmission electron microscope (JEOL, Peabody, 268 MA, USA) fitted with a bottom mount AMT digital camera (Danvers, MA, USA) ^24^.

*TEM of cross-sectioned EVs:* Suspensions of EVs were pelleted at 100,000 ×*g* in a Beckman airfuge for 20 min and the pellets were fixed in 2.5% glutaraldehyde in PBS overnight. The supernatant was removed and the cell pellets were washed 3x in PBS and post-fixed in 1% OsO_4_, 1% K_3_Fe(CN)_6_ for 1 hour. Following 3 additional PBS washes, the pellet was dehydrated through a graded series of 30-100% ethanol. After several changes of 100% resin over 24 hours, the pellet was embedded in a final change of resin, cured at 37 °C overnight, followed by additional hardening at 65°C for two more days. Ultrathin (70 nm) sections were collected on 200 mesh copper grids, stained with 2% uranyl acetate in 50% methanol for 10 minutes, followed by 1% lead citrate for 7 min. Sections were imaged using a JEOL JEM 1400 Plus transmission electron microscope (Peabody, MA) at 80 kV fitted with a side mount AMT 2k digital camera (Advanced Microscopy Techniques, Danvers, MA).

### 2.6. Western blot analysis of EV protein markers

The characteristic EV protein markers were evaluated using sodium dodecyl sulfate-polyacrylamide gel electrophoresis (SDS-PAGE) western blot analysis ^19^. Briefly, fifty µg EV protein lysates and hCMEC/D3 cell lysate were mixed with laemmli sample buffer and incubated at 95 °C for 5 min. The samples and premixed molecular weight markers (ladder) were separated on a 4-10% SDS-polyacrylamide gel at 120 V for two hours using a PowerPac Basic setup (BioRad Laboratories Inc.). The proteins were transferred onto a 0.45 µm nitrocellulose membrane at 75 V for 90 min using a transfer assembly (BioRad Laboratories Inc.). The membrane was washed with 0.1% tween-20-containing tris buffer saline (T-TBS) and blocked with Intercept blocking solution (Intercept blocking buffer:T-TBS:: 1:1) for an hour at room temperature. The membrane was incubated with rabbit anti-calnexin (1 µg/mL), rabbit anti-CD63 (1/1000 dilution), mouse anti-CD9 (0.2 µg/mL), mouse anti-ATP5A (1 µg/mL), rabbit anti-TOMM20 (1 µg/mL), and mouse anti-GAPDH (1 µg/mL) primary antibodies in blocking solution for overnight at 4 °C. The membrane was washed with T-TBS and incubated with anti-mouse AF790 (0.05 µg/mL), and anti-rabbit AF790 (0.05 µg/mL) secondary antibodies in a blocking solution for an hour at room temperature. The membrane was washed and scanned under 800 and 700-nm near-infrared channels using an Odyssey imager (LI-COR Inc., Lincoln, NE) at intensity setting 5.

### 2.7. Uptake of Mitotracker-labeled EVs into HBMECs using flow cytometry

#### 2.7.1. Isolation of mitochondria-labeled EVs

hCMEC/D3 cells (P32) were cultured in a T175 flask to confluency. The complete growth medium was removed, cells were washed with 1x PBS, and cells were incubated with 100 nM Mitotracker deep red (MitoT-red) diluted in a conditioned medium for 30 min in a humidified incubator. Next, the medium was replaced with serum-free medium after washing the cells with 1x PBS, and cells were kept in a humidified incubator for 24 h. Post-incubation, the conditioned medium was collected into centrifuge tubes. sEV (MitoT-red-sEV) and m/lEV (MitoT-red-m/lEV) from Mitotracker Red-labeled cells were isolated from the conditioned medium using the differential centrifugation method described in section 2.2. The EV protein content in MitoT-red-sEV and MitoT-red-m/lEV was determined using MicroBCA assay and the samples were stored at −20 °C until further use.

#### 2.7.2. Quantification of MitoT-EV uptake into recipient hCMEC/D3 cells using flow cytometry

hCMEC/D3 cells were cultured in 48-well plates at 50,000 cells/well in a complete growth medium. Unstained and untreated cells were used as a control, whereas cells stained with 100 nM of Mitotracker deep red (MitoT-red) for 45 min in a complete growth medium were used as a positive control. The cells were treated with MitoT-red-sEV and MitoT-red-m/lEV at 30, 75, and 150 µg EV protein/well in a complete growth medium for 72 h in a humidified incubator. The cells were also treated with unlabeled sEV and m/lEV at 150 µg EV protein/well in a complete growth medium for 72 h. Post-treatment, the cells were washed with 1x PBS, dissociated using TrypLE Express, diluted with PBS, and collected into centrifuge tubes. For each sample, an aliquot of a 100 µL cell suspension was analyzed through Attune NxT Flow cytometer and 10,000 events were recorded in FSC vs. SSC plots. The Mitotracker deep red fluorescence intensity was detected at 670/14 nm and percentage signal intensities were presented in histogram plots generated using Attune software version 3.2.3. Mitotracker deep red background signals were gated using the controls, including PBS and untreated cells.

### 2.8. Uptake of MitoT-EVs into primary human brain endothelial cells using fluorescence microscopy

#### 2.8.1. Uptake of MitoT-EVs into primary HBMEC monolayers

MitoT-red sEV and MitoT-red-m/lEV were isolated from the conditioned medium of hCMEC/D3 cells as described in section 2.6.1. Primary HBMEC (P6) were cultured in 96-well plates at 16,500 cells/well in a complete growth medium. Post-confluency, the cells were treated with MitoT-red-sEV and MitoT-red-m/lEV at 10, 25, and 50 µg EV protein/well in a complete growth medium for 24, 48, and 72 h in a humidified incubator. At each time-point, the cells were observed under an Olympus IX 73 epifluorescent inverted microscope (Olympus, Pittsburgh, PA) using the Cyanine-5 (Cy5, excitation 651 nm, and emission 670 nm) and bright-field channels at 20x magnification. Images were acquired using CellSens Dimension software (Olympus, USA).

#### 2.8.2. Colocalization of MitoT-EVs with the mitochondrial network in the recipient hCMEC/D3 and HBMEC monolayers

##### 2.8.2.1. Mitotracker green staining of recipient cell mitochondria

HBMEC and hCMEC/D3 cells were seeded in a 96 well-plate at 16,500 cells/well and incubated in a humidified incubator at 37°C. Post-confluency, the complete growth medium was removed, cells were washed with 1x PBS, and cells were incubated with 100 µM of Mitotracker green in a complete medium for 30 min. Post-treatment, the medium was replaced, washed with PBS, and incubated with MitoT-red-sEV or MitoT-red-m/lEV at 50 µg EV protein/well in a complete growth medium for 72 h in a humidified incubator. hCMEC/D3 cells stained with only Mitotracker green were used as a control. Post-incubation, the medium was removed, and cells were washed and incubated with phenol-red free and serum-containing DMEM-high glucose medium. The cells were then observed under an Olympus IX 73 epifluorescent inverted microscope (Olympus, Pittsburgh, PA) using Cyanine-5 channel (Cy5, excitation 651 nm, and emission 670 nm) to detect MitoT-red-EV uptake and GFP channel to detect Mitotracker Green signals at 20x magnification and images were acquired using CellSens Dimension software (Olympus, USA). Pearson’s correlation coefficient was obtained from the overlay images of Cy5 and GFP channels at constant signal intensities for both channels using the Cell Insight CX7 HCS microscope (ThermoFisher Scientific).

##### 2.8.2.2. Staining of a mitochondrial matrix protein in the recipient cells using CellLight Mitochondria-GFP BackMam reagent

We used CellLight Mitochondria-GFP BacMam to label a structural mitochondrial matrix protein ^25^. HBMECs were seeded in a 96 well-plate at 16,500 cells/well and incubated in a humidified incubator at 37°C. Post-confluency, the complete growth medium was removed, cells were washed with 1x PBS, and cells were incubated with CellLight Mitochondria-GFP reagent (at a dilution of 2 µL/10,000 cells as recommended by the manufacturer) for 16 h. Post-transduction, the medium was removed and cells were washed with 1x PBS. Next, the cells were incubated with MitoT-red-sEV and MitoT-red-m/lEV at 50 µg EV protein/well in a complete growth medium for 72 h in a humidified incubator. HBMECs transduced with only CellLight Mitochondria-GFP were used as a control. Post-incubation at 24 and 72 h, the medium was removed, and cells were washed and incubated with phenol-red free and serum-containing DMEM-high glucose medium. The cells were then observed under Olympus IX 73 epifluorescent inverted microscope (Olympus, Pittsburgh, PA) using Cyanine-5 channel (Cy5, excitation 651 nm, and emission 670 nm) to detect MitoT-EV uptake and GFP channel for CellLight Mitochondria-GFP signals at 20x magnification and images were acquired using CellSens Dimension software (Olympus, USA).

### 2.9. Effects of naïve EVs on the relative ATP levels in primary HBMECs under normoxic and ischemic/oxygen-glucose deprivation (OGD) conditions

To simulate ischemic conditions *in vitro*, HBMECs were exposed to different OGD settings, and the resulting cell viability was evaluated using an ATP assay ^19, 26, 27^. HBMECs were cultured in 96-well plates at 16,500 cells/well. Confluent HBMECs were incubated with OGD medium defined as follows: 120 mM NaCl, 5.4 mM KCl, 1.8 mM CaCl_2_, 0.8 mM MgCl_2_, 25 mM Tris-HCl, pH 7.4 for 0.5 to 24 h in either normoxic or hypoxic conditions ^28^. For normoxic conditions, the culture plates were incubated in a humidified incubator with 95% air and 5% carbon dioxide whereas for hypoxic conditions, the culture plates were incubated in an OGD chamber (Billups Rothenberg, Del Mar, CA) pre-flushed with 5% carbon dioxide, 5% hydrogen and 90% nitrogen at 37±0.5°C. For <u>normoxic conditions</u>, the medium was removed and the recipient HBMEC monolayers were incubated with hCMEC/D3-derived sEV, m/lEV, and sEV+m/lEV at doses of 10, 25, and 50 µg EV protein/well in a humidified incubator for 24, 48, and 72 h. For <u>hypoxic conditions</u>, the medium was removed and the recipient HBMEC monolayers were incubated with hCMEC/D3-derived sEV, m/lEV, and sEV+m/lEV at doses of 10, 25, and 50 µg EV protein/well in the OGD medium and a humidified incubator for 24 h. To evaluate the effect of EVs isolated from mitochondria complex I-inhibited cells on HBMEC ATP levels, confluent HBMECs were incubated with naïve sEVs, m/lEVs, RTN-sEVs, RTN-m/lEVs, OGM-sEVs or OGM-m/lEVs at 25 µg EV protein/well under OGD conditions. Normoxic HBMECs treated with complete culture medium and cells treated with OGD medium (OGD untreated cells) were used as controls.

Post-treatment, the treatment mixtures were replaced with pre-warmed complete growth medium, and an equal quantity of Cell Titer Glo 2.0 reagent was added. The plates were incubated for 15 min at RT on an orbital shaker in the dark and relative luminescence units were measured at 1 s integration time using a Synergy HTX multimode plate reader (BioTek Instruments Inc., Winooski, VT).

### 2.10. Measurement of mitochondrial function using Seahorse Analysis

The oxidative phosphorylation and glycolytic functions of hCMEC/D3 cells treated with EVs during normoxic conditions were evaluated using the Seahorse analysis by measuring oxygen consumption rate (OCR) and Extracellular Acidification rate (ECAR) ^23, 29^. hCMEC/D3 cells seeded at 20,000 cells/well were cultured in a Seahorse XF96 plate for four days. The cells were incubated with hCMEC/D3-derived sEV, m/lEV, and sEV+m/lEV for 24, 48, and 72h at 3.4 µg EV protein/well equivalent to 30 µg EV protein/cm^2^ in complete growth medium. Post-incubation, the medium was replaced with pre-warmed DMEM and subjected to Seahorse analysis ^23^. After measurement of baseline OCR, 2.5 µmol/L oligomycin A and 0.7 µmol/L carbonyl cyanide-p-trifluoromethoxyphenyl-hydrazone were consecutively added to measure the proton leak and maximal OCR, respectively ^29^. The total protein of the cells in each well was measured using Pierce BCA assay.

### 2.11. *In vivo* studies

#### Mice

Young male C57BL/6 mice (8-12 weeks) were purchased from Jackson laboratories. Mice were acclimated in our animal facilities for one week before use. Mice were housed 4-5 per cage, with a 12-h light/dark schedule in a temperature and humidity-controlled vivarium, with ad-libitum access to a pelleted rodent diet (irradiated LabDiet 5053, PicoLab® rodent diet 20) and filtered water. All experiments were approved by the Institutional Animal Care and Use Committee at the University of Texas Health Science Center in Houston. This study was performed in accordance with the guidelines provided by the National Institute of Health (NIH) and followed RIGOR guidelines. Only male animals were used in this study to allow comparability to earlier studies and to exclude potential sex and estrogen-related effects. Animals were randomly assigned to treatment groups.

#### Mouse middle cerebral artery occlusion stroke model

Focal transient cerebral ischemia was induced in mice by right middle cerebral artery occlusion (MCAO) followed by reperfusion as described previously ^30, 31^. Briefly, mice were initially anesthetized by being placed in a chamber with 4% isoflurane, and adequate sedation was confirmed by tail pinch for all surgical procedures. Surgery was performed under 1-2% continuous isoflurane. A 90-minute middle cerebral artery occlusion was achieved by blocking blood flow to MCA using a silicone-coated filament, via the external carotid artery. At the end of ischemia (90 minutes MCAO), mice were briefly re-anesthetized, and reperfusion was initiated by filament withdrawal. Body temperature was maintained by placing the mouse on a heating pad at ∼37°C during a surgical procedure. Two-hours after the onset of the stroke, mice were randomly assigned to receive 200 µL of m/lEVs/vehicle treatment by intravenous injection. Mice were euthanized 24 h after the stroke and brains were analyzed for infarct size using 2,3,5-triphenyl tetrazolium chloride (TTC) stained sections. Infarct analysis was performed by an investigator blinded to treatment groups.

### 2.12. Formulation of HSP27 mixtures with EVs, PEG-DET, and EV-PEG-DET

PEG-DET/HSP27 mixtures were prepared using a rapid mixing method. A PEG-DET polymer solution prepared in 10 mM HEPES buffer, pH 7.4 was mixed with 2 µg HSP27 at PEG-DET/HSP27 w/w ratio 0.05, 0.2, 1, 5, 10, and 20:1 for 30 s. The mixture was incubated at room temperature (RT) for 30 min. For the preparation of sEV/HSP27 and m/lEV/HSP27 mixtures, hCMEC/D3-derived EVs were suspended in 1x PBS were incubated with 2 µg HSP27 at EV protein/HSP27 w/w ratios 5, 10, and 15:1 in centrifuge tubes. The mixture was vortexed for 30 s and incubated at room temperature for 30 min. To prepare (PEG-DET/HSP27)/EV ternary mixtures, PEG-DET/HSP27 mixtures were prepared at 20:1 and 30:1 w/w ratios followed by incubation with 10 µg EV protein for 30 min at RT. The different mixtures were characterized by electrophoretic mobility, particle diameter, and zeta potential.

### 2.13. Native polyacrylamide gel electrophoresis (PAGE)

The mixtures of HSP27 with PEG-DET, sEV, and m/lEV were confirmed using native PAGE. A polyacrylamide gel consisting of 4% and 10% of acrylamide in the stacking and resolving sections, respectively, was prepared using a gel casting assembly (Bio-Rad Laboratories Inc., Hercules, CA) ^19, 26^. Native HSP27, PEG-DET/HSP27, and EV/HSP27 mixtures containing 2 µg of HSP27 at the indicated w/w ratios were mixed with native sample buffer (cat#1610738, Bio-Rad Laboratories Inc., Hercules, CA) and loaded into the gel lanes. Free PEG-DET polymer, naïve sEV, and m/lEV equivalent to indicated w/w ratios were used as controls. In an independent experiment, PEG-DET/HSP27 at w/w 20:1 and 30:1 was incubated with sEV, m/lEV, and EVs (sEV: m/lEV 1:1) at PEG-DET to EV protein 2:1 and 3:1 w/w ratios. The gel was run in 1x Tris-Glycine buffer, pH 8.3 at 100 V for 2 h at 2-8 °C using PowerPac Basic setup (Bio-Rad Laboratories Inc., Hercules, CA). Post-electrophoresis, the gel was washed with deionized water for 30 min and stained with 50 mL of Biosafe Coomassie blue G-250 for 1 h on an orbital shaker at room temperature. The gel was washed with deionized water for 30 min and scanned under an Odyssey imager (LI-COR Inc., Lincoln, NE) at an 800 nm channel and intensity setting 5. The band densities were quantified using ImageStudio 5.2 software.

### 2.14. Dynamic light scattering analysis of HSP27 mixtures

The average particle diameters, dispersity indices, and zeta potentials of the HSP27 mixtures were analyzed using a Malvern Zetasizer Pro (Worcestershire, UK). hCMEC/D3 cell line-derived naïve EVs at 0.5 mg EV protein/mL were diluted in either 1x PBS for size and dispersity index or 10 mM HEPES buffer pH 7.4 for zeta potential measurements. In addition, particle sizes and dispersity indices of native HSP27 protein at 20 µg/mL, PEG-DET/HSP27 mixtures at 10, 20, and 30:1 w/w ratios, sEV/HSP27 and m/lEV/HSP27 mixtures at 10:1 w/w ratio were measured in a low-volume disposable cuvette. Free PEG-DET equivalent to 10, 20, and 30:1 w/w ratios and naïve EVs equivalent to 10:1 w/w ratio were also analyzed. For the zeta potential of the above mixtures, a 50 µL sample of mixtures was further diluted in a zeta cuvette containing 10 mM HEPES buffer at pH 7.4. The samples were run in triplicate. Data are presented as average particle diameter ± standard deviation. The reported data are representative of 3 independent experiments.

### 2.15. Cytocompatibility of HSP27 mixtures with primary HBMEC and hCMEC/D3 monolayers

The cell viability of hCMEC/D3 and primary HBMEC monolayers treated with HSP27 mixtures was measured using Cell Titer Glo (ATP) assay. hCMEC/D3 and HBMEC were seeded at 16,500 cells/well in a 96 well-plate and cultured in a humidified incubator at 37±0.5°C. The growth medium was replaced with treatment mixtures containing either native HSP27, sEV/HSP27 (10:1), m/lEV/HSP27 (10:1), sEV+m/lEV at 1:1/HSP27 (10:1), or PEG-DET/HSP27 (20:1) at a dose of 2 µg HSP27 protein per well. Naïve sEV, m/lEV, equivalent amounts of sEV+m/lEV, and free PEG-DET equivalent to 20:1 in complete medium were used as controls. Polyethyleneimine (PEI) at 50 and 100 µg/mL in a complete growth medium was used as a positive control. The mixtures and controls were treated for 72 h in a humidified incubator at 37±0.5°C. Post-incubation, the ATP assay was performed using Cell Titer Glo 2.0 reagent as described earlier in section 2.8. The cell viability of HSP27 mixtures treated HBMECs was calculated using Equation 1.

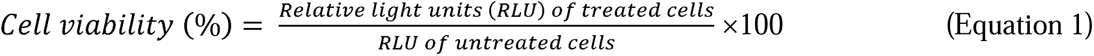

### 2.16. Paracellular permeability of TRITC-labeled 4.4 and 65-85 kD Dextran in HBMEC monolayers pretreated with EV/HSP27, PEG-DET/HSP27 binary mixtures and PEG-DET/EV/HSP27 ternary mixtures

#### 2.16.1 Hypoxic conditions (OGD only)

Primary HBMEC (P4-P9) in a complete growth medium were seeded at 50,000 cells/cm^2^ in a 24-well format cell culture insert (insert area: 0.33 cm^2^) for four days to form a complete endothelial monolayer. The medium was replaced every 48 h during this culturing period. The abluminal wells were filled with complete growth medium throughout the culturing period. The complete growth medium was replaced with 300 µL of growth medium containing 2 µg HSP27/well (formulated as described in 2.10) for 72 h. Post-treatment, the complete growth medium was replaced with 300 µL of OGD medium containing 1 µM TRITC-Dextran. The OGD medium containing 1µM TRITC-Dextran alone was used as a control. The abluminal chamber was filled with 0.5 mL of fresh complete growth medium. The untreated group was incubated in a humidified incubator whereas OGD treatment groups were incubated in an OGD chamber (as described earlier in section 2.8.). The concentration of TRITC-Dextran in the abluminal medium was measured at 4, and 24 h post-treatment. A 50 µL aliquot was collected at indicated time points from the abluminal side. An equal volume of fresh medium was replaced to maintain the sink conditions. The concentration of TRITC-Dextran was measured using Synergy HTX multimode plate reader at excitation 485/20 and emission 580/50 nm. The relative diffusion was calculated using **equation 2** shown below in **2.16.2**.

#### 2.16.2. OGD/reperfusion

Post-24 h of OGD exposure, HBMECs in the culture inserts were washed with 1x PBS. HBMECs were incubated with a complete growth medium containing 1 µM 65-85 or 4.4 kD TRITC-Dextran in a 24-well format cell culture insert for 1h – 24h (reperfusion) in a humidified incubator. The abluminal chamber was filled with 0.5 mL of fresh complete growth medium. The concentration of TRITC-Dextran in the abluminal medium was measured at 1, 2, 4, and 24 h during reperfusion. A 50 µL of samples from the abluminal side were collected at indicated time points. An equal volume of fresh medium was replaced to maintain the sink conditions. The concentration of TRITC-Dextran was measured using Synergy HTX multimode plate reader at excitation 485/20 and emission 580/50 nm. The relative diffusion of TRITC-Dextran at each time point was calculated as the ratio of TRITC-Dextran present in the basolateral compartment between the treated groups and untreated control.

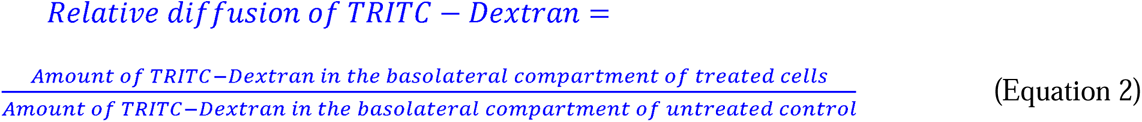

### 2.17. Statistical analysis

Statistically significant differences between the means of controls and treatment groups or within treatment groups were determined using one-way analysis of variance (ANOVA) or two-way ANOVA at 95% confidence intervals using GraphPad Prism 9 (GraphPad Software, LLC). The notations for the different levels of significance are indicated as follows: *p<0.05, **p<0.01, ***p<0.001, ****p<0.0001.

## 3. Results

### 3.1. EVs retained their physicochemical characteristics and membrane integrity upon revival from storage conditions

We collectively refer to sEVs and m/lEVs as EVs wherever applicable. We used dynamic light scattering to measure particle size and dispersity index of freshly-isolated EVs using a Malvern Zetasizer Pro (**Fig. 1a**). Average particle diameters of freshly isolated sEV were 109.9±1.1 nm with a dispersity index (DI) of 0.39±0.02 and m/lEV was 228.8±15.35 nm with DI of 0.35±0.03 (**Fig. 1a**). The representative particle size distribution (PSD) of sEV and m/lEV were depicted in **Fig. S1a-b**. It should be noted that intensity-weighted PSD of sEV showed about 74% of particles were < 200 nm in diameter ranging from 20 nm to 197 nm (**Fig. S1a**). PSD of m/lEV showed about 54% of particles were >200 nm particle diameter ranging from 200 nm to 500 nm (**Fig. S1b**). It is known that EVs are heterogenous in size and the larger EVs are known to show diameters ranging from 100 – 1000 nm ^32^. Next, We studied the effect of storage conditions on the retention of physicochemical characteristics of hCMEC/D3 BEC-derived sEVs and m/lEVs using dynamic light scattering (**Fig. S1c-h**). During freeze-thaw (FT) cycles, sEV particle diameters significantly (p<0.0001) increased after the first FT cycle from 110 nm to 140 nm. sEV diameter gradually increased to 150 nm at FT2 and FT3 (**Fig. S1c**). It should be noted that the sEV size remained <200 nm during the storage conditions (**Fig. S1c**). There were no significant differences in the particle diameters of m/lEVs between a freshly-isolated sample and when measured after three freeze-thaw cycles (**Fig. S1d**). The average particle diameter of m/lEV remained above 200 nm during storage conditions (**Fig. S1d**). Additionally, sEV and m/lEVs retained a consistent dispersity index after three freeze-thaw cycles (**Fig. S1e,f**). sEVs and m/lEVs showed an initial negative zeta potential of about −22 mV that ranged between −15 and −30 mV after three consecutive freeze-thaw cycles (**Fig. 1a, S1g-h**). The particle concentration of freshly isolated sEVs and m/lEVs in PBS was determined using nanoparticle tracking analysis and was found to be 4.6 and 5.1 ×10^8^ particles/mL, respectively (**Fig. 1a**).

**Fig. 1.**
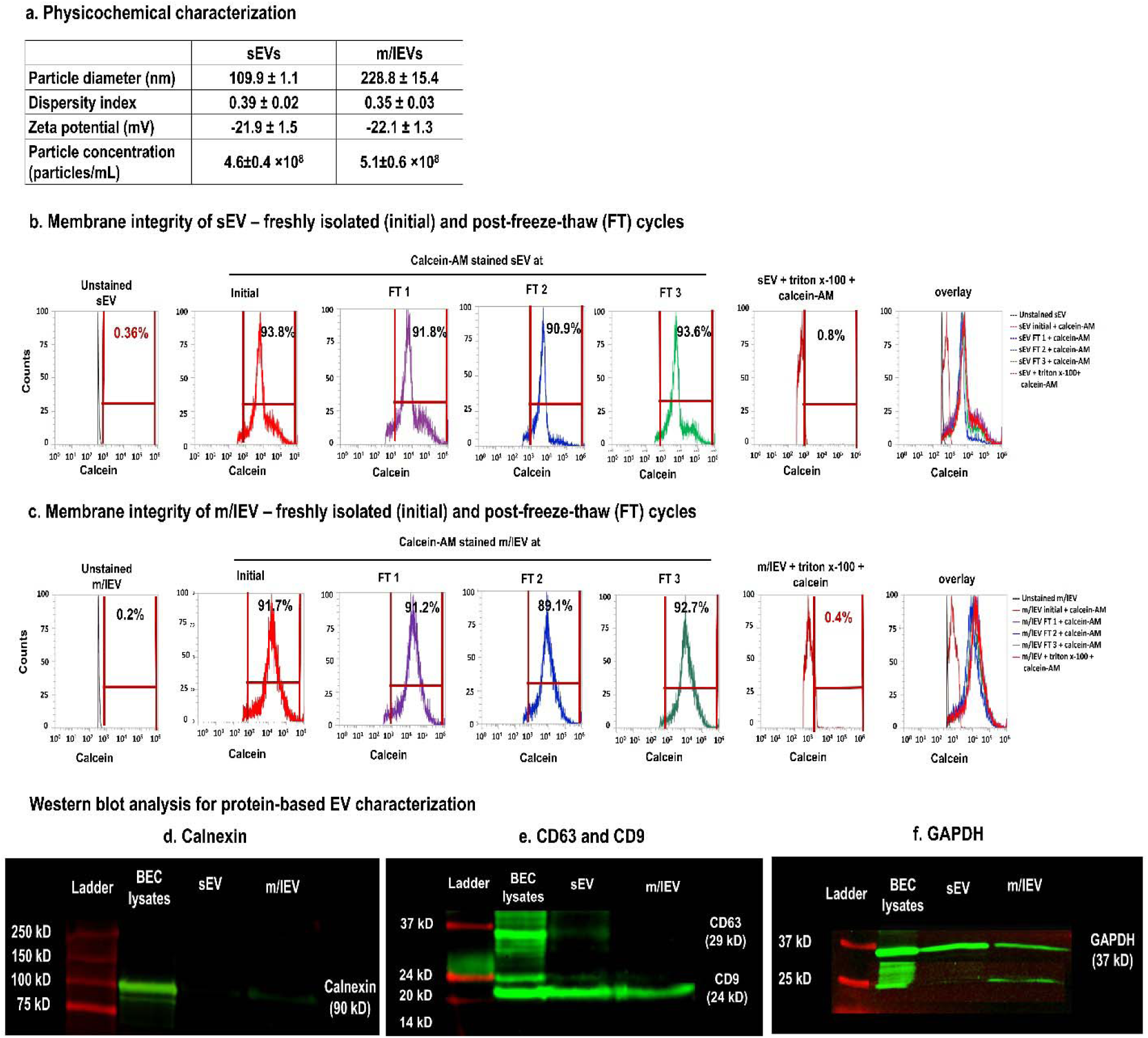
Physicochemical characteristics, membrane integrity, and protein content-based EV characterization. (**a**) Average particle diameters, dispersity indices, and zeta potentials of fresh hCMEC/D3 BEC-derived EVs were determined using dynamic light scattering on a Malvern Zetasizer Pro-Red. Samples were diluted to 0.1 mg protein/mL in 1x PBS for particle diameter and 10 mM HEPES buffer pH 7.4 for zeta potential measurements. Data are presented as mean±SD of n=3 measurements. EV particle concentrations were measured using nanoparticle tracking analysis (NTA). For NTA, stock samples of sEV and m/lEV were diluted 100 times in PBS and analyzed on a multiple-laser ZetaView f-NTA Nanoparticle Tracking Analyzer (Particle Metrix Inc., Mebane, NC). Three 60 s videos were acquired at 520 nm laser wavelengths for particle diameter and concentration measurements. Average particle concentrations were reported as mean ± standard deviation of n=3 measurements. (**b-c**) The histograms of calcein-positive events of intact sEVs (**b**) and m/lEVs (**c**) at initial and post-three FT cycles were detected using a small particle side scatter 488/10-nm filter in an Attune flow cytometer. Unstained EVs were used to gate the histograms for estimating percentage calcein-positive counts. (**d-f**) Detection of membrane protein markers of sEVs and m/lEVs-derived from hCMEC/D3 cells using western blot analysis. Western blot shows the relative expression of calnexin (90 kD), CD63 (29 kD), CD9 (24 kD), and GAPDH (37 kD) protein in sEVs and m/lEVs. The total protein content in the cell lysates, sEV, and m/lEV was measured using a micro BCA assay. Fifty μg protein/sample was incubated with Laemmli sample buffer at 95°C for 5 min. The sample was run on sodium dodecyl sulfate-polyacrylamide gel electrophoresis, proteins were transferred to a nitrocellulose membrane, and the membrane was incubated with a blocking buffer. The blot was incubated with primary antibodies overnight at 4°C, and secondary antibodies at room temperature. The blots were imaged on the 800 nm channel using an Odyssey imager (LI-COR Inc. Lincoln, NE) at intensity setting 5 and processed using ImageStudio 5.2 software. Uncropped western blots of triplicate runs are shown in **Fig. S3**.

The membrane integrity of EVs upon revival from cold storage was determined using a previously reported calcein-based flow cytometry assay ^19^. Prior to analyzing the EVs, we first calibrated the flow cytometer using polystyrene beads of particle diameters ranging from 0.1 to 2 µm using a small particle side scatter filter (488/10 nm, BL1) (**Fig. S1i**). The relative position of EV clusters in the FSC/SSC overlay plots was proportional to the particle diameter since an increase in diameter showed a right-upward shift of the clusters. Notably, sEVs and m/lEVs clusters overlapped to a large extent in the area corresponding to 0.1-0.2 µm bead diameters suggesting that this protocol allowed us to detect EV-sized particles (**Fig. S1j**). The particle counts for PBS/calcein AM, unlabeled-sEVs and m/lEVs, PBS/Triton X-100/calcein AM mixture (sample processing controls) were acquired on SSC/BL1 density plots and were used for gating calcein AM positive signals. About 90% of freshly-isolated sEVs and m/lEVs were calcein-positive suggesting that EVs retained intact membranes after the ultracentrifugation and resuspension processes (**Fig. 1b-c**). Importantly, >85-90% sEVs and m/lEVs maintained their membrane integrity after three consecutive freeze-thaw cycles confirming the lack of significant membrane damage during and upon revival from storage conditions (**Fig. 1b-c**). In addition, EVs lysed with Triton X-100 showed less than 10% calcein-positive particle counts demonstrating the specificity of calcein signal intensities associated with the intact EVs (**Fig. 1b-c**).

We additionally performed an ATP assay to determine if EVs retained their functionality upon revival from frozen storage conditions. HBMECs exposed to OGD conditions for 24 h showed about a 60% reduction in relative ATP levels compared to normoxic HBMECs (**Fig. S2a,b**). Importantly, HBMECs treated with freshly isolated sEVs at 50 µg EV protein in OGD conditions showed about a three-fold (p<0.0001) increase in relative ATP levels compared to untreated cells (**Fig. S2a**). OGD HBMECs treated with 50 µg sEV protein (sEVs were revived after three subsequent freeze/thaw cycles) showed a similar increase in relative ATP levels compared to untreated cells (**Fig. S2a**). Importantly, there were no statistical (p>0.05) differences between relative ATP levels of HBMECs treated with either freshly isolated sEVs or sEVs post-freeze/thaw. Like sEV, OGD HBMECs treated with freshly isolated m/lEV showed about a four-fold and statistically significant (p<0.0001) increase in relative ATP levels compared to untreated cells (**Fig. S2b**). Importantly, there was no statistical (p>0.05) difference between relative ATP levels of HBMECs treated with either freshly isolated m/lEV or freeze/thaw-subjected m/lEV. These observations suggest that EVs retain their functionality after frozen storage conditions.

Per the MISEV 2018 guidelines for protein-based characterization of EVs, we used the following categories of protein markers (**Fig. 1a**). **a.** *Transmembrane proteins associated with the plasma membrane and/or endosomes:* tetraspanins CD63 and CD9. **b.** *Cytosolic proteins recovered in EVs:* GAPDH. **c.** *Transmembrane and soluble proteins associated with other intracellular compartments than plasma membrane/endosomes:* ATP5A and TOMM20. **d.** Lastly, we used the lack of calnexin (an endoplasmic reticulum marker) as a purity marker to confirm the lack of cellular contaminants in EVs. We performed western blot analysis for protein content-based characterization of freshly isolated sEVs and m/lEVs (**Fig. 1d-f**). We evaluated the expression of calnexin (endoplasmic reticulum marker, **Fig. 1d**), CD63 (exosome or sEV marker, **Fig. 1e**), CD9 (a common marker for both sEV and m/lEV, **Fig. 1e** (14)), and GAPDH (**Fig. 1f**) in our EV samples (**Fig. 1d**).

Calnexin was used as a control to evaluate whether sEVs and m/lEVs contained endoplasmic reticulum (ER) contaminants. BEC cell lysates showed a strong band of calnexin at 75 kD as expected (**Fig. 1d**). sEV did not show any calnexin expression, whereas m/lEV showed a faint band suggesting that sEVs are free of ER contaminants and m/lEV contained minimal ER contaminants (**Fig. 1d**). Next, only sEV showed CD63 expression whereas m/lEV did not show CD63 band at its characteristic molecular weight of 28 kD suggesting the purity of sEV isolation (**Fig. 1e**). Lastly, CD9 tetraspanin, a common 25 kD EV membrane protein marker, was seen in both sEVs and m/lEVs (**Fig. 1e**). Both EVs expressed GAPDH (**Fig. 1f**). The presence of GAPDH in sEV and m/lEV in our western blot is likely due to the incorporation of cytosolic components in EVs during their biogenesis.

### 3.2. m/lEVs contain mitochondria and transfer their mitochondrial load to recipient primary HBMECs and hCMEC/D3 cells

We studied the morphology of naïve sEVs and m/lEVs using transmission electron microscopy (TEM, **Fig. 2a-b**). Negative stain TEM analysis showed sEV of <200 nm—scale bar (**Fig. 2a**). Negatively stained m/lEV showed structures >200 nm (**Fig. 2b**). It should be noted that our TEM images are comparable to the published literature (16). The raw images of the selected EVs are shown in **Fig. S4a,b**.

**Fig. 2:**
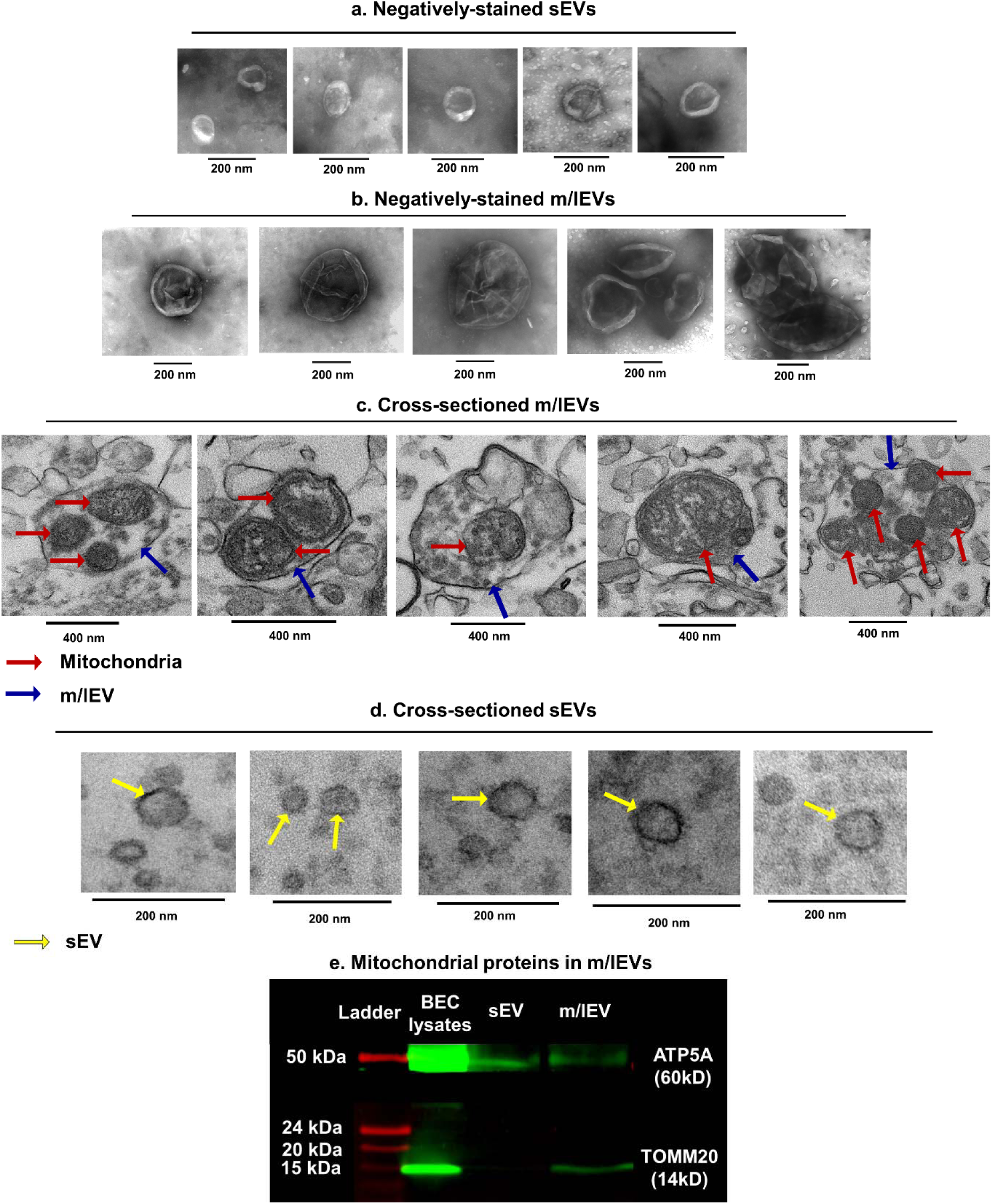
Transmission electron microscopy and western blot analysis of sEVs and m/lEVs. Negative stain TEM images of hCMEC/D3-derived sEV (**a**) and m/lEV (**b**). Representative TEM images of sectioned m/lEVs **(c)** and sEVs **(d)**. m/lEVs (blue arrow) contained one or more mitochondria (electron-dense structures, maroon arrows). sEV cross sections (yellow arrow) lacked electron-dense structures in the lumen. Scale bars of 400 nm and 200 nm. (**e**) Detection of mitochondrial proteins in sEVs and m/lEVs using western blot analysis. The uncropped western blots of triplicate runs are shown in **Fig. S7a,b.**

We acquired multiple TEM images of cross-sectioned m/lEVs (**Fig. 2c and Fig. S5**). The blue arrow points to the m/lEV membrane, whereas the maroon arrow points to mitochondria in m/lEVs. Mitochondria were detected as electron dense structures in the lumen of m/lEVs (**Fig. 2c**). Notably, m/lEVs contain one or multiple mitochondria with varying sizes and morphologies (**Fig. 2c**). The uncropped raw images of the data shown in **Fig. 2c** are presented in **Fig S4a,b**. In contrast, images of sEV cross-sections looked empty and did not contain electron dense structures in the lumen (yellow arrow, **Fig. 2b**), suggesting the absence of mitochondria in sEVs. The structure and shape of the mitochondria in m/lEVs (**Fig. 2c**) were comparable with the mitochondria observed in the hCMEC/D3 cell buds/protrusions (**Fig. S4**). The high-speed centrifugation and extensive sample processing during the sample pelleting step prior to cutting ultrathin sections for TEM imaging likely resulted in the observed mitochondrial morphology. We noted mitochondria structures with a characteristic double membrane that were highly similar to published reports ^33–36^. It should also be noted that these previous reports have demonstrated functional activity of the m/lEV/extracellular mitochondria. Our data showed that m/lEV- and not sEV sections contained mitochondria.

We performed a western blot analysis of sEV and m/lEV lysates for determining the presence of mitochondrial proteins, including ATP5A (subunit of the mitochondrial ATP synthase) and TOMM20, an outer mitochondrial membrane protein (**Fig. 2e**). Our data showed that while both sEVs and m/lEVs contained ATP5A protein, TOMM20 was selectively present in m/lEVs (**Fig. 2e**). This suggests that while sEVs contain mitochondrial proteins, only m/lEVs contain mitochondria. The western blot aligned with the TEM data confirming the presence of mitochondria in m/lEVs. Collectively, our data demonstrate the presence of mitochondria in the m/lEVs but not sEVs.

Next, we wanted to determine if EVs can transfer their mitochondrial load into the recipient primary human brain microvascular endothelial cells (HBMECs) (**Fig 3a-c**). We isolated sEVs and m/lEVs from hCMEC/D3 cells pre-stained with Mitotracker Red (MitoT-red). First, we performed cytocompatibility of MitoT-red stained EVs at the same doses i.e., 30, 75, and 150 µg EV protein/cm^2^ for 72 h in normoxic conditions (**Fig. S8**). Untreated cells were used as control, whereas polyethyleneimine, PEI, at 50 µg/mL was used as a positive control for the ATP assay. PEI-treated BECs showed a significant (p<0.0001) decrease in cell viability (**Fig. S8**) suggesting the sensitivity and functionality of ATP assay measuring the cell viability of treated BECs. HBMECs treated with MitoT-red-sEV and MitoT-red-m/lEV at 10, 30, and as high as 150 µg EV protein/cm^2^ for 72 h did not show any significant reduction in BECs viability (**Fig. S8**), suggesting that MitoT-red-EVs were cytocompatible with recipient BECs up to 150 µg EV protein/cm^2^ for 72 h exposure.

**Fig. 3.**
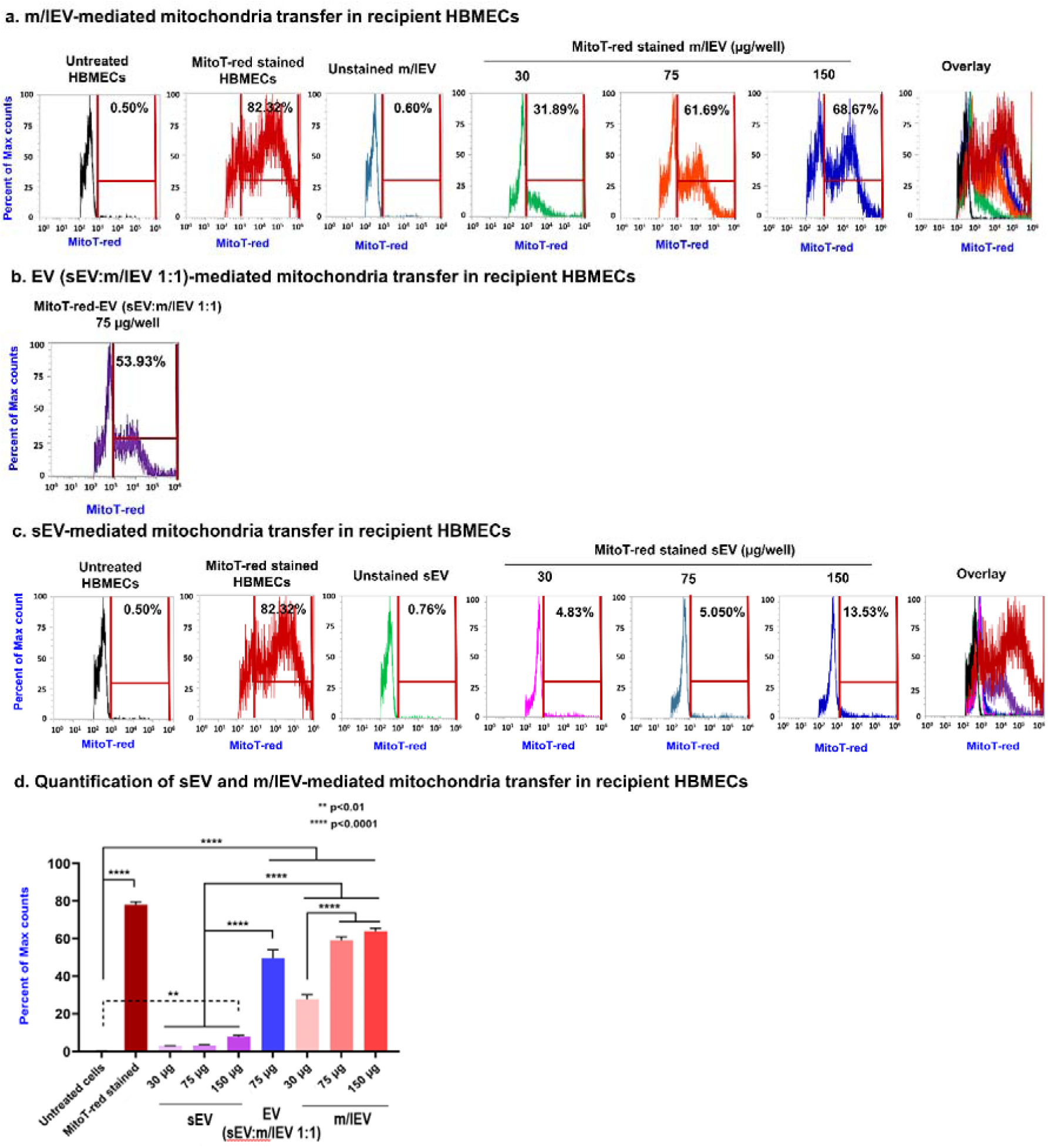
Transfer of EV mitochondria into recipient HBMECs cells at varying doses. HBMECs cell were cultured in 48-well plates for 48 h in a humidified incubator. Cells were then incubated with th indicated amounts of MitoTracker red-labeled samples: (MitoT-red)-sEV, MitoT-red-EV (at a 1:1 sEV:m/lEV ratio, collectively referred to as EVs) and MitoT-red-m/lEV diluted in complete growth medium for 72 h. Post-incubation, the cells were washed, collected, and run through the Attune NxT flow cytometer. The histograms of hCMEC/D3 cells treated at indicated doses of MitoT-red-m/lEVs (**a**), sEV:m/lEV 1:1 (**b**), and sEV (**c**) were collected using a 674/10-nm side scatter filter in the Attune flow cytometer. Untreated HBMECs and unstained EVs were used as controls to gate the background signals in histograms. MitoT-red-stained HBMECs were used as a positive control to gate the histograms for MitoT-red-positive counts. Subsequently, this gate was applied to quantify the percentage of MitoT-red HBMECs treated with MitoT-red-EVs. (**d**) Quantification of sEV and m/lEV-mediated mitochondria transfer in recipient HBMECs. Data represent mean±SD of n=3.

HBMECs were treated with MitoT-red-sEV and MitoT-red-m/lEV at 30 to 150 µg EV protein/well for 72 h and MitoT-red signals in the recipient HBMECs were measured using flow cytometry. The intensity of MitoT-red-EVs in the recipient HBMECs (**Fig. 3a-c**) was analyzed using histogram plots. Untreated HBMECs cells were used as control and were gated for data analysis. Cells pre-stained with MitoT-red were used as a positive control and showed about 82% MitoT signals suggesting the presence of polarized mitochondria (**Fig. 3a**). Cells treated with unstained m/lEV did not show any MitoT-red signals in HBMECs suggesting the absence of non-specific MitoT signals (**Fig. 3a**). Cells treated with MitoT-red-m/lEV at a low dose of 30 µg dose showed about 32% MitoT-positive signals suggesting mitochondrial transfer into the recipient HBMECs (**Fig. 3a, d)**. The fraction of cells showing MitoT+ve signals increased from 61% at 75 µg to 68% at 150 µg m/lEV dose (**Fig. 3a, d**). Cells treated with EV (sEV: m/lEV 1:1) at 75 µg EV protein showed about 54% MioT-red signal intensity suggesting that inclusion of m/lEVs in the EV mixture treatment increased mitochondrial transfer to HBMECs (**Fig. 3b, d**). sEVs at lower doses of 30 and 75 µg protein per well showed <5% of mitochondrial transfer that increased to only about 13% at the 150 µg dose (**Fig. 3c, d**). As expected, m/lEVs showed a greater transfer of mitochondria into recipient HBMECs compared to sEVs consistent with the presence of mitochondria in m/lEVs (**Fig. 2c-e**).

We also studied EV-meditated mitochondrial transfer into recipient hCMEC/D3 cells (**Fig. S9**). Consistent with our observations from the HBMECs, hCMEC/D3 cells treated with MitoT-red-m/lEV showed a dose-dependent increase in mitochondrial transfer from 48% at 30 µg to 68% at 150 µg dose (**Fig. S9a,c**). We noted a dose-dependent increase in the mitochondrial transfer that increased from 13% at 30 µg sEV to about 33% at the 150 µg dose (**Fig. S9b,c**). Interestingly, sEV-mediated a greater extent of mitochondrial transfer into hCMEC/D3 cells compared to the levels noted in primary HBMECs (33 vs. 13%) (**Fig. 3c and Fig. S9b**). We performed additional flow cytometry studies at lower MitoT-red EV doses (i.e., 10, 25, and 50 µg EV protein/well) in 48-well plates and have added the data in **Fig. S10**.

### 3.3. EVs transferred polarized mitochondria to recipient primary HBMECs

We tested if EV mitochondria can be transferred into the recipient BECs by isolating EVs from donor BECs pre-stained with MitoT-red. We subsequently incubated recipient BECs with MitoT-red-EVs and observed MitoT-red signals for 24 to 72 h using fluorescence microscopy. We first determined the uptake of EV-associated mitochondria in recipient primary HBMECs and hCMEC/D3 cells. The recipient HBMECs or hCMEC/D3 cells were treated with Mitotracker red-stained EVs (MitoT-red-EV) for 72 h prior to observation with epifluorescence microscopy. No MitoT-red-associated signals were observed in unstained/untreated primary HBMECs, and HBMECs treated with only MitoT-red (100 µM for 30 min in complete culture medium) showed purple puncta in Cy5 channel, suggesting that only MitoT-red specific signals were detected under the Cy5 channel (**Fig. 4a**). MitoT-red-sEV at 10 and 25 µg doses did not show positive signals in HBMECs for 48 h; however, MitoT-red-sEV at 50 µg doses showed faint intracellular Cy5 signals at 48 and 72 h, suggesting low levels of uptake after 48 h (**Fig 4a**). In contrast, strong intracellular signals in HBMECs treated with MitoT-sEV+m/lEV at the 25 µg (**Fig. 4a**) dose suggested that the inclusion of m/lEVs in the EV mixture led to an efficient uptake of polarized mitochondria into HBMECs. Moreover, the increase in the sEV+m/lEV-mediated mitochondrial transfer was statistically significant (p<0.01) at 48 h compared to sEVs alone (**Fig. 4b**).

**Fig. 4.**
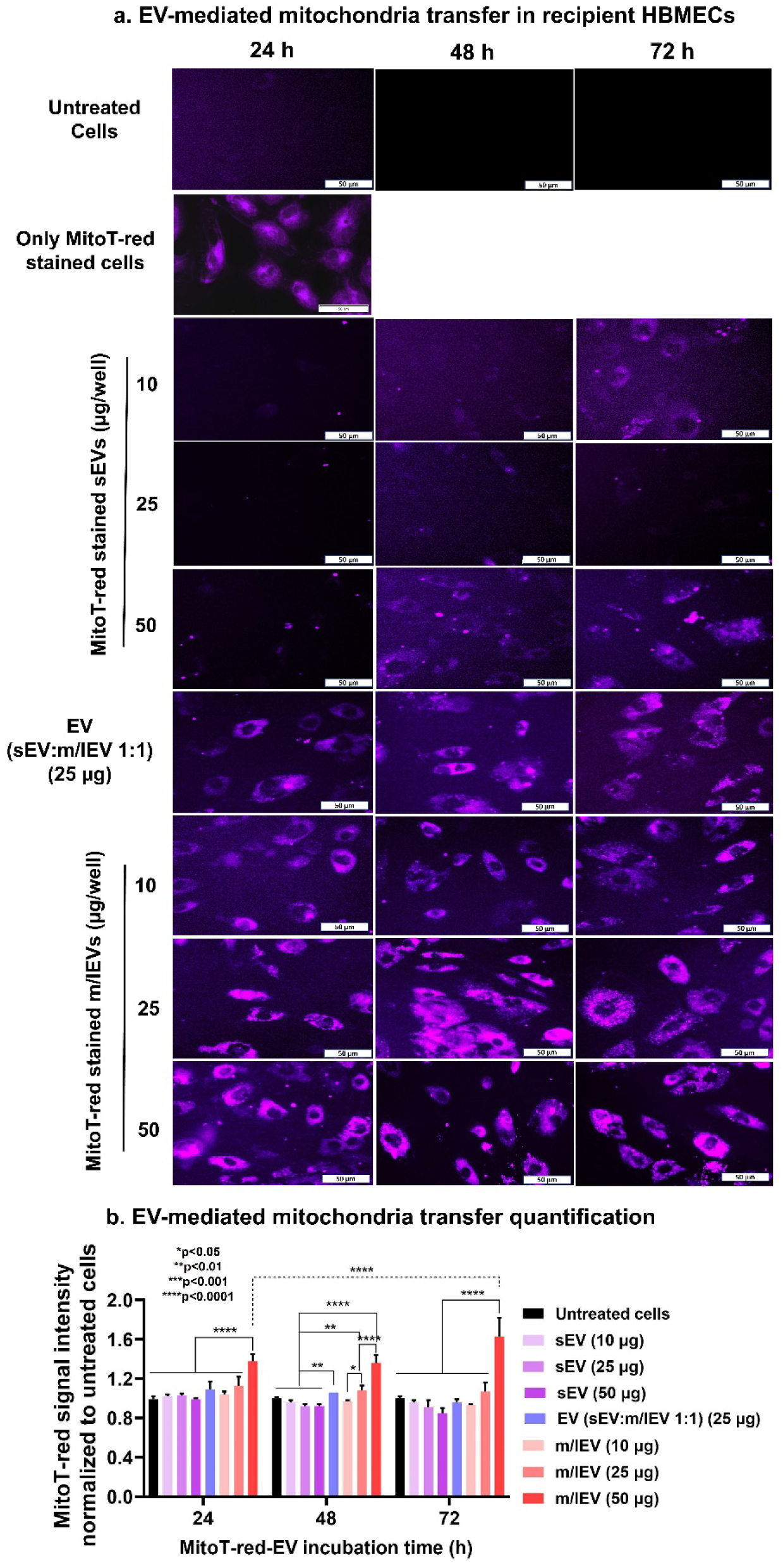
Transfer of EV mitochondria into the recipient HBMEC at varying doses and incubation times. **(a)** HBMECs were cultured in 96-well plates until 80% confluency in a humidified incubator. Cells were then incubated with the indicated amounts of MitoTracker red-labeled samples: (MitoT-red)- sEVs, MitoT-red-EV (at a 1:1 sEV: m/lEV ratio), MitoT-red-m/lEVs diluted in complete growth medium for 24, 48, and 72 h. Post-incubation, the cells were washed and incubated with phenol-red-free growth medium. Intracellular MitoT-red-sEV/sEV:m/lEV/m/lEV signals were observed under an Olympus IX 73 epifluorescent inverted microscope using Cy5 channel (purple puncta) at 20x magnification. Scale bar: 50 µm. **(b)** EV mitochondria transfer quantification. HBMECs were treated with the indicated samples and doses for 24, 48, and 72 h. At each time point, from each control and treatment group, at least three images were acquired and the total sum of grayscale signal intensities in the Cy5 channel was estimated using Olympus CellSens software. The measured intensities were normalized with those of the untreated cells.

Interestingly, MitoT-red-m/lEV at a dose as low as 10 µg showed efficient uptake in HBMECs 72 h post-exposure. The levels of uptake increased significantly (p<0.0001) as the dose of MitoT-red-m/lEV increased from 10 to 50 µg at 48 h (**Fig. 4a,b**). The superior transfer of m/lEV-mediated mitochondria into HBMECs is likely due to a greater enrichment of functional mitochondria in the m/lEVs compared to sEVs. Therefore, we expected that m/lEVs may increase the cellular bioenergetics of the recipient BECs at lower doses compared to sEVs. We further confirmed the effect of dose and type of EV subtype (m/lEVs vs. sEVs) on the transfer of mitochondria into hCMEC/D3 cells (**Fig. S11a**). Similar to the observations noted in the primary HBMEC cultures at 72 h post-exposure, sEVs at 50 µg protein/well showed faint MitoT-red+ signals in the recipient hCMEC/D3 cells, whereas cells treated with m/lEVs showed a dose-dependent increase (p<0.01) in mitochondrial transfer compared to sEV-treated cells (**Fig. S11a,b**).

We additionally studied the uptake of EV mitochondria into the recipient HBMECs at different time points, including 4, 8, 16, 24, 48, and 72 h (**Fig. S12**). We incubated primary HBMECs with MitoT-red-stained m/lEVs and sEVs at a 30 µg EV protein dose. EV mitochondria (purple puncta) in recipient HBMECs were again observed under Cy5. HBMECs treated with MitoT-red-stained m/lEVs for 4 h did not show MitoT-red signals in the recipient cells; however, faint purple signals were observed at 8 and 16 h incubation times (**Fig. S12a**). These data indicate that m/lEV mitochondrial transfer in recipient HBMECs takes about 8 to 16 h of exposure. m/lEV incubation for 24 h resulted in intense MitoT-red associated purple puncta in HBMECs, suggesting efficient m/lEV mitochondria transfer into HBMECs (**Fig. S12a**). At 48 h exposure time, m/lEV MitoT-red signals increased compared to 24 h and remained consistent for 72 h. It can be inferred that m/lEVs show time-dependent mitochondria transfer, wherein mitochondria transfer from m/lEV to HBMECs was likely initiated at about 8 h, peaked at 48 h, and persisted for at least 72 h in recipient HBMECs (**Fig. S12a**).

In contrast, incubation of MitoT-red-stained sEVs with HBMECs for 4 to 48 h did not show any MitoT-red signals in HBMECs, suggesting a considerably lower mitochondrial load in the sEVs compared to m/lEVs (**Fig. S12b**). HBMECs treated with MitoT-red-stained sEVs for 72 h showed only faint MitoT-red signals (**Fig. S12b**). The representative images of EV mitochondria load transfer into the recipient HBMEC at varying EV doses for 24 h are shown at higher magnification in **Fig. S13**.

### 3.4. EV-transferred mitochondria colocalized with the mitochondrial network in the recipient BECs

We wanted to determine if EV-mitochondria colocalized with the mitochondrial network of the recipient BECs. We isolated MitoT-red-stained polarized mitochondria from the donor BECs and the recipient BEC mitochondria were prestained using Mitotracker green (MitoT-green). Post-treatment of the recipient BECs, the overlap of these fluorescent signals was observed under an epifluorescent microscope. HBMECs and hCMEC/D3 cells prestained with Mitotracker green were subsequently incubated with MitoT-red-sEVs and MitoT-red-m/lEVs at 10, 25, and 50 µg doses for 72 h. Cytosolic, diffuse MitoT-green signals were observed under the GFP channel whereas punctate MitoT-red EV signals were captured under the Cy5 channel (purple puncta). The prestaining of HBMECs with Mitotracker green resulted in robust fluorescent signals for 72 h and incubation with Mitrotracker red-stained EVs did not affect the Mitotracker green signals.

The absence of GFP and Cy5 signals in the untreated cells suggested the absence of non-specific signals at the respective channel settings (**Fig. 5a**). The cells prestained with MitoT-green alone showed green cytosolic signals associated with the recipient mitochondria (**Fig. 5a**). MitoT-red-m/lEVs showed a greater intensity of Cy5 signals at all tested doses compared to MitoT-red-sEVs, once again demonstrating that m/lEV contain a greater mitochondrial load (**Fig. 5a**). The overlay images of recipient HBMEC mitochondria and EV-associated polarized mitochondria in the m/lEV-treated cells showed considerably higher colocalization compared to sEV-treated cells (**Fig. 5a**). The Pearson’s correlation coefficient (PCC) of GFP and Cy5 channel intensities demonstrated that m/lEV exposure resulted in a statistically significant (p<0.0001), greater degree of mitochondria colocalization compared to sEVs (**Fig. 5b**). Similar to the primary HBMECs, hCMEC/D3 cells prestained with Mitotracker green and treated with MitoT-red-m/lEVs showed a greater degree of colocalization indicated by the overlap of MitoT-green and MitoT-red signals compared to sEV-treated cells (**Fig. S14**).

**Fig. 5.**
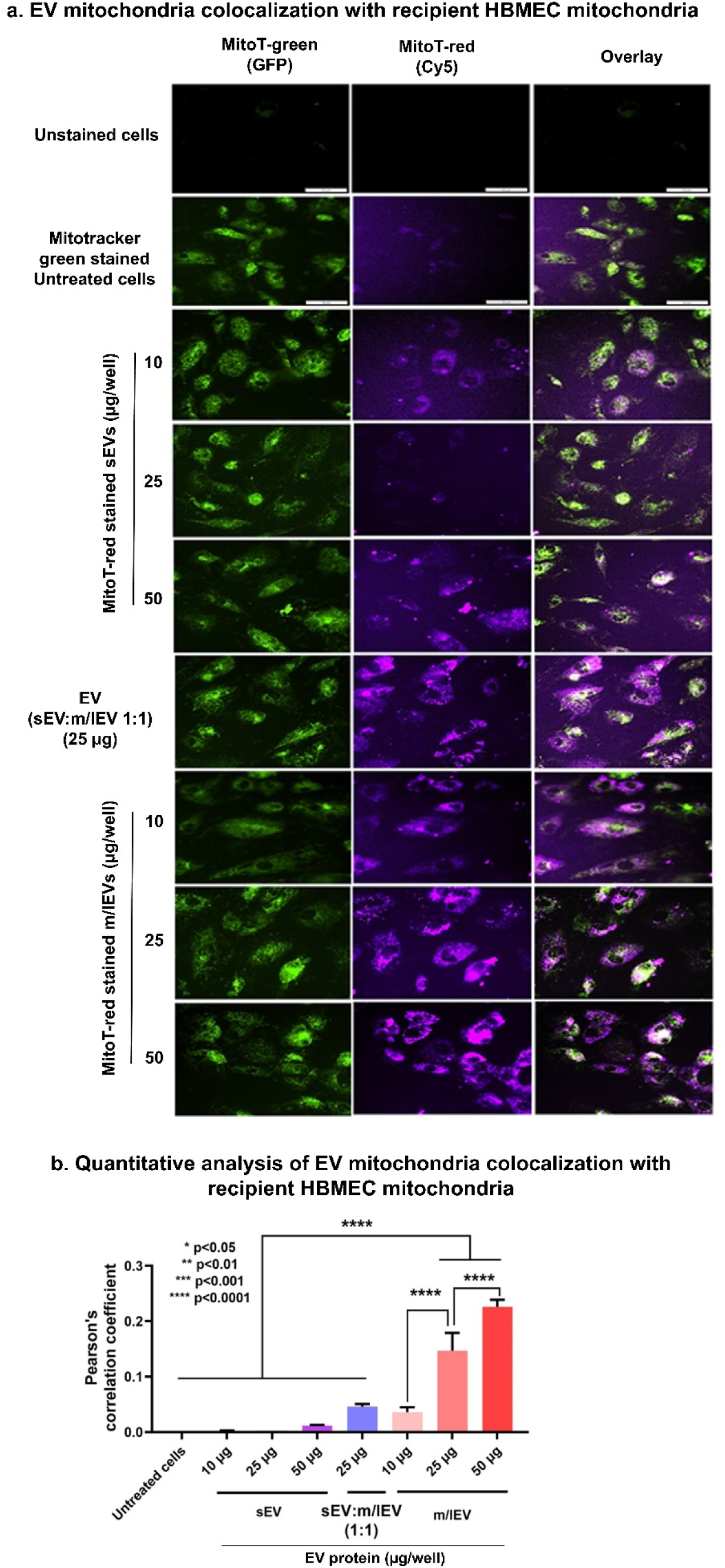
Colocalization of EV mitochondria with the recipient HBMEC mitochondria. **(a)** HBMECs were cultured in 96-well plates until 80% confluency in a humidified incubator. HBMECs were stained with Mitotracker Green for 30 min. Post-staining, the cells were washed and treated with the indicated doses of MitoT-red-sEV, MitoT-red-EVs (at a 1:1 sEV: m/lEV ratio, collectively referred to as EVs), and MitoT-red-m/lEV for 72 h. Untreated cells and cells stained with MitoTracker Green only were used as controls. Post-incubation, the treatment mixture was replaced with phenol-red-free growth medium. The Mitotracker green staining in recipient HBMEC was acquired using the GFP channel, whereas the purple fluorescence associated with EV mitochondria was captured using Cy5 channel in an Olympus IX 73 epifluorescent inverted microscope. Colocalization of the mitochondria signals was confirmed by the presence of yellow signals in the overlay images. Scale bar: 50 µm. (**b**) Pearson’s correlation coefficient was obtained from the overlay images of Cy5 and GFP channels at constant signal intensities for both channels using a Cell Insight CX7 HCS microscope. Data are presented as mean±SD (n=3 images per treatment group). *p<0.05, **p<0.01, ***p<0.001, ****p<0.0001

We performed additional microscopic studies (**Fig. S15 and S16**) to confirm that the observed PCC in **Fig. 5b** for colocalization of MitoT-red m/lEVs with MitoT-green-stained recipient cell mitochondria is specific to MitoT-red signals. The details are explained in **Fig. S15 and S16** of the supplementary file.

We further confirmed the colocalization of EV-mitochondria with mitochondria of the recipient BECs using an orthogonal technique that allowed us to confirm that the EV-mitochondria colocalized with the mitochondrial matrix in the recipient BECs. HBMECs were first transduced with CellLight Mitochondria-GFP (CellLight-MitoGFP) before MitoT-red-EV treatment. Despite the low absolute frequency of transduction ^37^, the cells showed strong GFP fluorescence suggesting that CellLight-MitoGFP transduction effectively tagged the alpha pyruvate mitochondrial matrix protein in the recipient HBMECs (**Fig. S17**). Similar to **Fig. 5**, MitoT-red-m/lEV at a dose of 50 µg showed efficient transfer of mitochondria (purple puncta) into the recipient HBMECs. The EV MitoT-red puncta signals colocalized with recipient mitochondria (CellLight-MitoGFP) at 72 h post-exposure. Notably, cells exposed to MitoT-red m/lEVs showed stronger Cy5 signals compared to MitoT-red-sEV-treated cells confirming again that m/lEVs contain more mitochondrial load compared to sEVs. The Cy5 signal intensity in m/lEV-treated cells was considerably increased at 72 h post-exposure compared to the 24 h time point suggesting that 72 h is an optimal exposure period for m/lEVs internalization into the recipient HBMECs (**Fig. S17**). Importantly, the overlap of the structural protein-tagged recipient BEC mitochondria with the MitoT-red-stained m/lEVs indicates that the m/lEV-delivered mitochondria colocalized with the recipient cell mitochondria.

### 3.5. Naïve EVs increased HBMEC ATP levels under normoxic and hypoxic conditions

Once we confirmed EV mitochondria transfer to the recipient BECs, we determined the relative ATP levels of EV-treated BECs under normoxic and hypoxic conditions. One of the main functions of mitochondria is to synthesize ATP from ADP during mitochondrial aerobic respiration, and therefore, we measured relative ATP levels in the recipient BECs treated with sEVs or m/lEVs using a luciferase-based ATP assay. The effect of BEC-derived EVs on the ATP levels in recipient primary HBMECs was first evaluated under normoxic conditions using an ATP assay. Primary HBMECs were treated with sEVs and m/lEVs at 10, 25, and 50 µg EV protein per well for 24, 48, and 72 h. **Fig. 6a** shows that the increase in HBMEC ATP levels upon sEV and m/lEV treatment for 24 h was not statistically significant (p>0.05). Importantly, HBMECs treated at a dose of 25 µg sEVs for 48 h showed a significant (p<0.01) increase in ATP levels compared to untreated cells. Interestingly, cells treated with m/lEVs at 25 and 50 µg doses showed a dose-dependent and significant (p<0.0001) increase in relative ATP levels compared to untreated cells. In addition, the m/lEV-mediated increase in ATP levels was significantly (p<0.05) higher compared to sEVs after 48 h exposure (**Fig. 6b**). At 72 h post-exposure, m/lEVs at 25 and 50 µg doses showed a dose-dependent and significant (p<0.01) increase in relative ATP levels compared to untreated cells under normoxic conditions (**Fig. 6c**). In contrast, sEVs did not show any significant increase in relative HBMEC ATP levels compared to the control. In addition, m/lEV-treated HBMECs showed significantly higher ATP levels compared to sEVs at 72 h at all tested doses. Thus, the m/lEV-mediated significant increase in relative ATP levels at 48 h (**Fig. 6b**) and 72 h (**Fig. 6c**) may likely be due to their innate mitochondrial load—including mitochondria, mitochondrial DNA, and mitochondrial proteins.

**Fig. 6.**
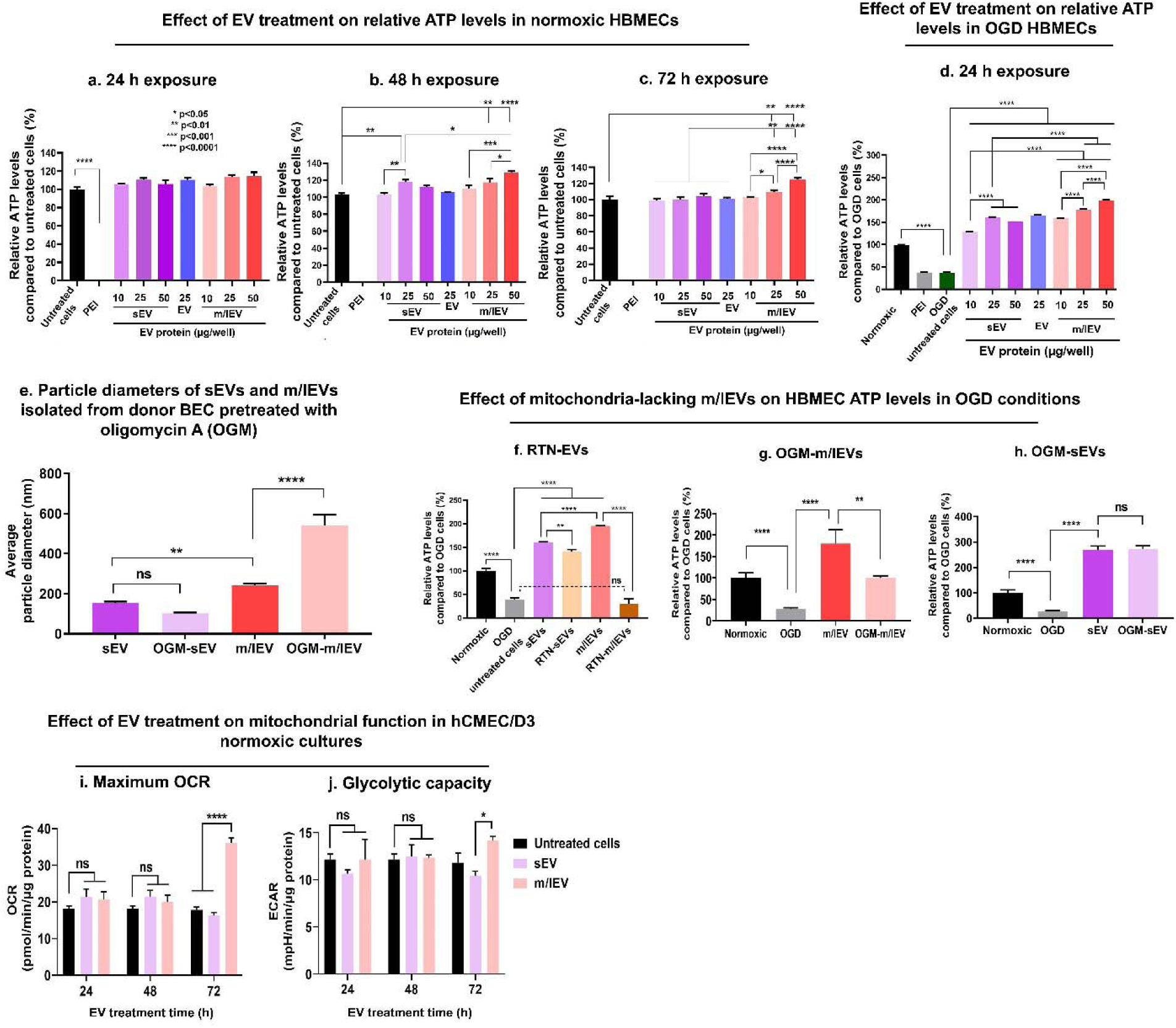
EV-mediated increase in HBMEC ATP levels and mitochondrial respiration during normoxic and hypoxic conditions. HBMEC cells were cultured in the 96-well plates until 80% confluency in a humidified incubator. (**a-c**) Confluent monolayers were treated with sEV, m/lEV, and EVs (sEV: m/lEV 1:1) at the indicated amounts for 24 (**a**), 48 (**b**), and 72h (**c**). Polyethyleneimine, PEI, at 50 µg/mL was used as a positive control for the ATP assay. Post-treatment, cells were incubated with a 1:1 mixture of fresh growth medium and Cell titer Glo reagent. The relative luminescence units (RLU) of the samples were measured using a SYNERGY HTX multimode plate reader at 1s integration time. Relative ATP levels were calculated by normalizing the RLU of treatment groups to the RLU of control, untreated cells. (**d**) Confluent HBMECs were treated with the indicated doses of sEV and m/lEV in OGD medium and cell viability was measured 24 h post-treatment while untreated cells were used as a control. Data represent mean ±SD (n=3). (**e**) Particle diameters of naïve and OGM-EVs. EVs were isolated from either untreated hCMEC/D3 cells (naïve sEVs and m/lEVs) or cells pretreated 1 µM of oligomycin A (OGM-sEV and OGM-m/lEVs). Naïve and OGM-EVs were suspended in 1x PBS at 0.1 mg EV protein/mL concentration and particle diameters were measured using Malvern Zetasizer Pro. (**f-h**) Effect of RTN-EV and OGM-EVs treatment on HBMEC ATP levels in OGD conditions. Recipient HBMECs were incubated with naïve, RTN-EVs, and OGM-sEVs. EVs were isolated from the conditioned medium of hCMEC/D3 BECs pretreated with 1 µM OGM for 4 h. Confluent HBMECs were treated with the indicated samples at 25 µg EV protein/well in OGD condition for 24 h. Normoxic cells and cells treated with OGD medium (untreated cells) were used as controls. Post-treatment, relative ATP levels were measured using the Cell Titer Glo-based ATP assay. Data represent mean ±SD (n=3). (**i-j**) EV-mediated increase in recipient cell mitochondrial respiration and glycolysis capacity: Cells were cultured in a Seahorse XF96 plate for 4 days at 20,000 cells/well. sEV, and m/lEV were diluted in complete growth medium at 3.4 µg EV protein/well and cells were incubated in a humidified incubator for 24, 48, and 72 h. Post-treatment at each time point, the medium was replaced with DMEM and maximum oxygen consumption rate (OCR) (**i**), and glycolytic capacity (**j**) by measuring extracellular acidification rate (ECAR) were analyzed using the Seahorse XFe96 analyzer. Data represents mean±SEM (n=3). * p<0.05, ** p<0.01, *** p<0.001, **** p<0.0001.

We then exposed primary HBMECs under oxygen-glucose deprived (OGD) conditions for 24 h to simulate ischemic conditions (**Fig. S18**). We studied the effects of naïve sEVs and m/lEVs on the cell survival of OGD-exposed primary HBMECs. HBMECs in OGD medium were incubated with sEVs and m/lEVs at 10, 25, and 50 µg EV protein/well for 24 h. The relative ATP levels of EV-treated HBMECs were compared with untreated HBMECs maintained in an OGD medium (**Fig. 6d**). sEV and m/lEV at all treated doses showed a significant (p<0.0001) increase in relative ATP levels compared to control, untreated HBMECs (**Fig. 6d**). In addition, an increase in sEV and m/lEV dose from 10 to 25 µg EV protein/well showed a significant (p<0.0001) increase in ATP levels. HBMECs treated with m/lEVs at 50 µg/well showed a maximum, ca. five-fold increase in ATP levels compared to untreated cells. Importantly, m/lEVs showed significantly (p<0.0001) higher HBMEC ATP levels compared to sEVs at the same dose (25 and 50 µg), suggesting that m/lEVs outperformed sEVs in increasing HBMEC cellular energetics under ischemic conditions. We also confirmed the EV-mediated increase in endothelial ATP levels under OGD conditions in the recipient hCMEC/D3 cells (**Fig. S19**). Consistent with the primary HBMECs, hCMEC/D3 cells treated with sEVs and m/lEVs at 10, 25, and 50 µg EV protein/well showed about a three to four-fold increase in endothelial ATP levels compared to untreated cells. Moreover, the EV-mediated increases in ischemic hCMEC/D3 ATP levels were dose-dependent. Lastly, m/lEV-treated ischemic hCMEC/D3 cells showed a greater increase in ATP levels compared to sEV-treated cells (**Fig. S19**).

We conducted the following studies to study the effects of EV mitochondria. We isolated sEVs and m/lEVs from donor hCMEC/D3 cells pre-treated with oligomycin A (OGM, mitochondria electron transport complex V inhibitor) to determine if the m/lEV-meditated increase in recipient HBMEC ATP levels was associated with m/lEV mitochondria. Donor hCMEC/D3 cells were treated with OGM at 1 µM concentrations for 4 h in a complete culture medium. Then cells were washed with 1x PBS and incubated with serum-free medium for 24 in a humidified incubator at 37°C. Post-incubation, OGM-sEVs, and OGM-m/lEVs were isolated from the EV-conditioned medium using the sequential ultracentrifugation method. First, we measured particle diameters of naïve and OGM-EVs using dynamic light scattering (**Fig. 6e**). Naïve sEV showed a particle diameter of about 150 nm whereas OGM-sEV showed particle diameters of about 110 nm (**Fig. 6e**). There was no statistical difference between particle diameters of naïve sEV and OGM-sEVs suggesting that OGM-mediated inhibition of mitochondria function in donor cells did not affect the physicochemical properties of sEVs. On the other hand, naïve m/lEV showed an average particle diameter of about 220 nm that significantly (p<0.0001) increased to 550 nm for OGN-m/lEV (**Fig. 6e**). OGM-mediated inhibition of mitochondria function in the donor BECs selectively increased particle diameter of OGM-m/lEVs compared to naïve m/lEVs likely due to OGM-mediated depolarization of donor mitochondria. This depolarization likely increased mitochondria fission and subsequent mitophagy that may have led to the incorporation of a greater number of depolarized mitochondria in m/lEVs ^38^. The increased mitochondrial load in m/lEVs likely increased OGM-m/lEV size compared to naïve m/lEVs. Since sEV did not contain mitochondria, OGM pretreatment to the donor cells did not affect the particle diameters of OGM-sEVs.

We performed an additional ATP assay to determine if the m/lEV-meditated increase in HBMECs ATP levels was associated with m/lEV mitochondria (**Fig. 6f**). OGD treatment showed a significant (p<0.0001) reduction in HBMEC ATP levels compared to normoxic cells (**Fig. 6f**). Consistent with the above studies (**Fig. 6d**), naïve sEVs and m/lEVs showed a significant (p<0.0001) increase in HBMEC ATP levels compared to the OGD control (**Fig. 6f**). m/lEV-mediated increase in ATP levels were significantly (p<0.0001) higher than sEVs suggesting that m/lEVs outperformed sEVs in increasing HBMEC cellular energetics under ischemic conditions. Importantly, RTN-m/lEVs treated HBMECs did not show a significant (p>0.05) increase in ATP levels compared to OGD control (**Fig. 6f**). Notably, at the same dose, m/lEV-mediated increase in ATP levels were significantly (p<0.0001) higher than RTN-m/lEVs treatment suggesting that the m/lEV-mediated increase in ATP levels is likely a function of their innate mitochondrial load—including mitochondria and mitochondrial proteins. In contrast, RTN-sEV treatment showed a significant (p<0.0001) increase in ATP levels compared to OGD control (**Fig. 6f**). RTN-sEV-mediated increase in ATP levels was significantly (p<0.01) lower than sEVs. Notably, the suppression of ATP is much more profound in the m/lEVs than in the sEVs, suggesting that the increase in ATP mediated by m/lEVs is even more dependent on mitochondrial complex I function.

In contrast, OGM-m/lEVs showed a significant (p<0.0001) decrease in HBMEC ATP levels compared to naïve m/lEVs (**Fig. 6g**) suggesting that the m/lEV-mediated increase in ATP levels is likely a function of their innate mitochondrial load— including mitochondria and mitochondrial proteins. In contrast, OGM-sEV treatment showed a significant (p<0.0001) increase in ATP levels compared to OGD control (**Fig. 6h**). OGM-sEV-mediated increase in ATP levels was significantly (p<0.01) higher than sEVs (**Fig. 6h**). Notably, the suppression of ATP is much more profound in the m/lEVs than in the sEVs, suggesting that the increase in ATP mediated by m/lEVs is even more dependent on mitochondrial complex I and V function.

### 3.6. EVs increased the oxidative phosphorylation and glycolytic functions of recipient BECs under normoxic conditions

The mitochondrial function of hCMEC/D3 cells treated with EVs under normoxic conditions was evaluated using Seahorse analysis by measuring their oxygen consumption rate (OCR). hCMEC/D3 cells were treated with sEVs and m/lEVs at 3.4 µg protein/well in a complete growth medium for 24, 48, and 72 h. sEV or m/lEV-treated cells did not show changes in maximal OCR compared to untreated cells at 24 and 48 h post-exposure (**Fig. 6i**). In contrast, BECs exposed to sEVs and m/lEVs for 72 h showed a significant (p<0.0001) increase in maximum OCR compared to control, untreated cells (**Fig. 6i**). Importantly, the m/lEV-mediated increase in OCR was significantly (p<0.05) higher compared to sEV-treated cells suggesting that m/lEVs outperformed sEVs in increasing the recipient BECs’ mitochondrial function (**Fig. 6i**). The m/lEV-mediated increase in mitochondrial function was consistent with the m/lEV-mediated increase in cell viability (**Fig. 6c**), intracellular uptake of m/lEV mitochondria (**Fig. 3-4**), and the co-localization of m/lEV-associated mitochondria with the recipient BEC’s mitochondrial network (**Fig. 5)**.

We further evaluated the effects of EVs on non-mitochondrial energy generation pathways such as glycolytic capacity in the recipient BECs. Extracellular acidification rate (ECAR) is a key indicator of cellular glycolysis and can be determined in real-time by measuring free protons in a Seahorse plate transient microchamber ^39^. ECAR (basal glycolysis rate and glycolytic capacity) was measured in hCMEC/D3 cells treated with sEVs and m/lEVs (**Fig. 6j**). We did not note any changes in the glycolytic capacity of hCMEC/D3 cells pretreated with sEVs and m/lEVs for 24 and 48 h compared to untreated cells. However, treatment with m/lEVs for 72 h showed a significantly (p<0.05) greater glycolytic capacity compared to untreated cells and sEV-treated cells (**Fig. 6j**). To summarize the first part of this study, our data demonstrated that (1) m/lEVs but not sEVs contain mitochondria, (2) m/lEVs outperformed sEVs in transferring mitochondrial components to recipient BECs, (3) m/lEVs resulted in a greater magnitude of relative ATP levels and mitochondrial functions (OCR and ECAR) compared to sEV-treated BECs, and (4) m/lEVs isolated from rotenone and oligomycin A exposed BECs did not increase recipient BEC ATP levels.

The second goal of the present work was to evaluate the effects of an exogenous HSP27 protein formulated with EVs in an OGD-exposed BEC model of ischemia/reperfusion injury. We formulated HSP27 protein with EVs and a synthetic cationic polymer, PEG-DET. We studied the physicochemical characteristics of the formed mixtures and evaluated their effects on the paracellular permeability of small and large molecular mass fluorescent tracers across primary HBMECs exposed to OGD.

### 3.7. m/lEVs showed neuroprotection in a mouse model of stroke

In this pilot experiment, we determined the feasibility of whether m/lEV treatment is safe from any adverse effects when administered intravenously (*i.v.*) to the mice (**Fig. 7**). We treated mice 2 h after the onset of stroke with 200 µL of m/lEVs or vehicle. Mice were euthanized 24 h after stroke and brains were analyzed for infarct size using 2,3,5-triphenyl tetrazolium chloride (TTC)-stained sections (**Fig. 7a**). Despite the small cohort size, we observed a trend toward neuroprotection (**Fig. 7b**) in m/lEV-treated mice compared to vehicle-treated mice. Importantly, we did not observe any detrimental effect as a result of m/lEV treatment—this is a significant finding given that this is the first report where mitochondria-containing m/lEVs were injected into a live animal. To date, all EV stroke studies have only injected sEVs/exosomes ^40–42^.

**Fig. 7.**
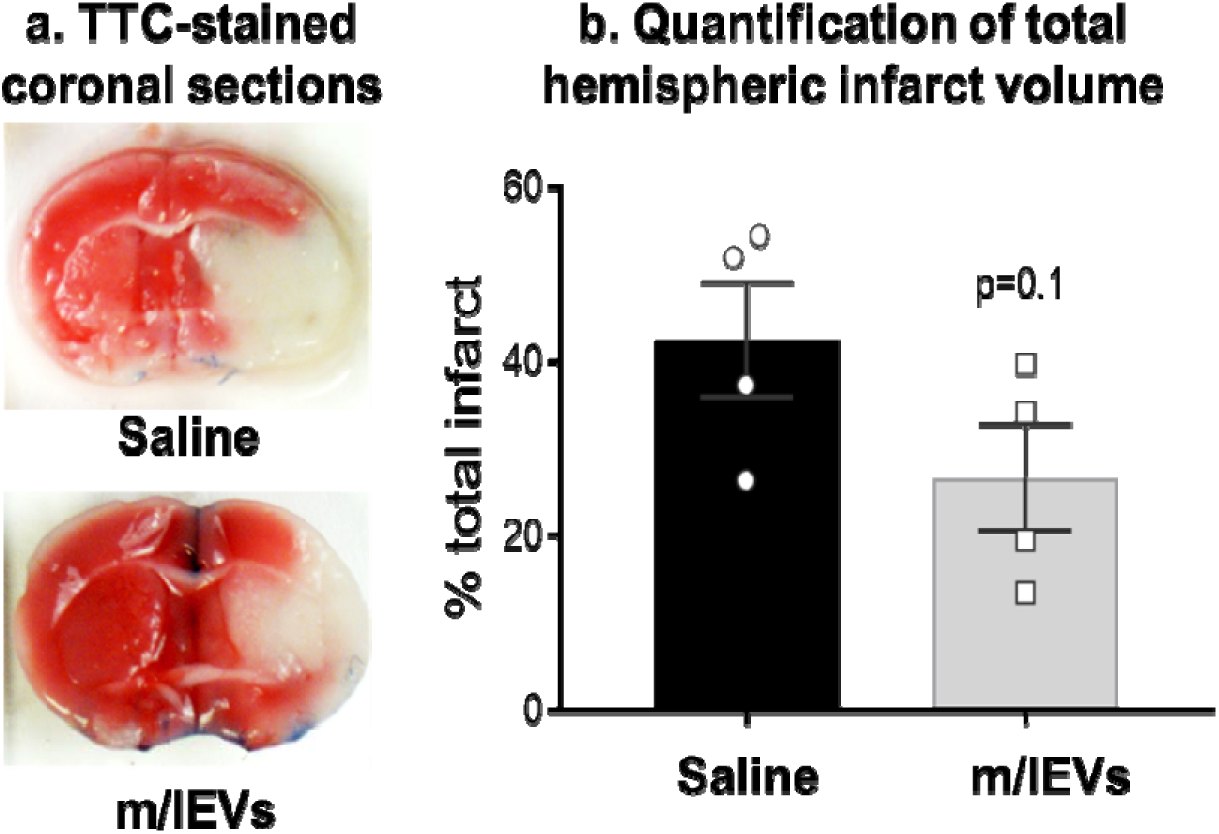
Pilot study demonstrating potential neuroprotective effects of m/lEVs in a mouse middle cerebral artery occlusion model of ischemia/reperfusion injury (stroke). **(a)** Representative 2,3,5-triphenyl tetrazolium chloride (TTC)-stained coronal sections of the vehicle and m/lEV-treated stroke brains from young male mice. **(b)** Quantification of total hemispheric infarct volume at 24 h post-stroke. Data are mean ± SEM (n=4) and were analyzed using an unpaired t-test.

### 3.8. Exogenous HSP27 protein mixtures with PEG-DET, EVs, and PEG-DET-EV mixtures

The mixtures of HSP27 protein with PEG-DET and EVs were confirmed by studying the electrophoretic mobility of HSP27 in a native polyacrylamide gel electrophoresis (PAGE) setup. Native recombinant human HSP27 at the running buffer of pH 8.3 carries a net negative charge (estimated charge: −4.2 mV) ^43^, and therefore, it migrated from the loading spot towards the anode during electrophoresis (**Fig. 8a**). First, the interactions of PEG-DET with HSP27 was studied by comparing the relative changes in HSP27 band densities at polymer: protein weight/weight (w/w) ratios ranging from 0.05:1 to 20:1 (**Fig. 8a,b**). Compared to native HSP27 (100%, **Fig. 8b**), the relative band density of PEG-DET/HSP27 mixtures at w/w 0.2:1 was considerably reduced to about 43%. As the w/w ratios increased, there was a gradual and significant decrease in HSP27 band densities (**Fig. 8a,b**). At PEG-DET/HSP27 w/w 10:1 and 20:1, the mean HSP27 band density decreased to nearly 25% suggesting that PEG-DET may form electrostatic interactions with HSP27 at physiological pH. The free polymer did not show any non-specific staining at w/w 1 and 20:1.

**Fig. 8.**
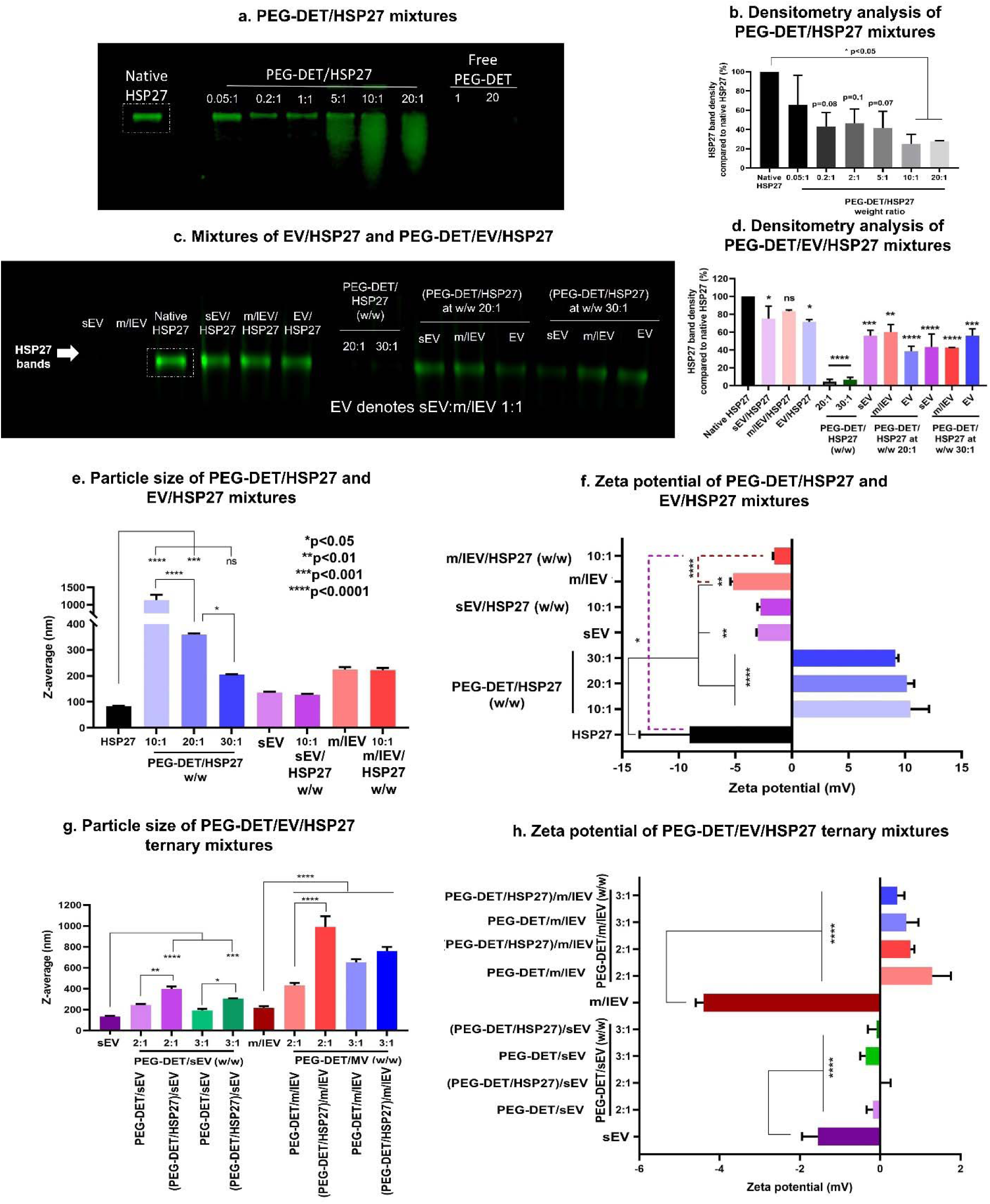
Formation of EV- and PEG-DET/HSP27 binary/ternary mixtures. (**a**) Native polyacrylamide gel electrophoresis (PAGE) for PEG-DET/HSP27 mixtures. Native HSP27, PEG-DET/HSP27 at indicated weight ratios, and free PEG-DET polymers were mixed with 1x native sample buffer and loaded in an SDS-free 4-10% polyacrylamide gel at 1 µg HSP27 per lane. (**c)** Native PAGE for hCMEC/D3-derived EV/HSP27 mixtures: Native HSP27 and mixtures of sEV/HSP27, and m/lEV/HSP27 at 10:1 weight/weight (w/w) ratios were loaded in a SDS-free 4-10% polyacrylamide gel at 1 µg HSP27 per lane. Free sEV and m/lEV equivalent to the amounts in 10:1 w/w mixtures were used as controls. Native PAGE for (PEG-DET/HSP27)/EV ternary mixtures. PEG-DET/HSP27 mixtures were prepared at 20:1 and 30:1 w/w ratios followed by incubation with 10 µg of EVs. The indicated samples were loaded in the gel at 1 µg HSP27/lane. Each gel was run at 100 V for 2 h and stained using Biosafe Coomassie G250. The gel was then scanned at 800 nm using an Odyssey imager at intensity setting 5. (**b,d**) Densitometry analysis was performed by measuring band densities of HSP27 in the different experimental groups in comparison to the band density of native HSP27 in the respective gel using Image Studio 5.0 software *p<0.05 (**e-h**) Physicochemical characterization of HSP27 mixtures with PEG-DET and EVs. Average particle diameters (**e**), and zeta potentials (**f**) of PEG-DET/HSP27 mixtures and EV/HSP27 mixtures at the indicated w/w ratios. Average particle diameters (**g**), and zeta potentials (**h**) of PEG-DET/EV and (PEG-DET/HSP27)/EV ternary mixtures at the indicated weight ratios. The samples containing 1 µg HSP27 protein were diluted to 50 µL in 10 mM HEPES buffer pH 7.4 for particle diameter measurements. The diluted samples were further diluted to 800 µL in 10 mM HEPES buffer pH 7.4 for zeta potential measurements. Data represent mean±SD (n=3). * p<0.05, **p<0.01, ***p<0.001, ****p<0.0001

hCMEC/D3 BEC-derived sEVs and m/lEVs were mixed with HSP27 at EV protein/HSP27 protein w/w ratios of 10:1 and the electrophoretic mobility of HSP27 was studied using native PAGE (**Fig. 8c-d)**. Mixtures of HSP27 with EVs did not affect the migration of HSP27 at the tested w/w ratios (**Fig. 8c**). The signal intensity of free/native HSP27 was set as 100%, and the relative band density of HSP27 in the different EV/HSP27 mixtures was compared with the band density of native HSP27. Although the bands look similar, densitometry analysis showed that sEV/HSP27 and m/lEV/HSP27 mixtures decreased HSP27 band densities to about 77.5±20% and 84.1±2.4% compared to native HSP27 (100%) (**Fig. 8c-d, and Fig. S20**).

We then formulated ternary mixtures of PEG-DET/HSP27 with EVs, and the resulting changes in HSP27 band intensity were studied using native PAGE followed by densitometry analysis (**Fig. 8c**). sEVs, m/lEVs, and EV/HSP27 at w/w 10:1 showed about 20-40% reduction in band density whereas PEG-DET/HSP27 at w/w 20:1 and 30:1 showed >90% HSP27 reduction compared to native HSP27 (**Fig. 8d**). The HSP27 band density was decreased by 40-50% when PEG-DET/HSP27 at w/w 20:1 were incubated with 10 µg of sEV, m/lEV, and sEV+m/lEV 1:1 (EV). The % extent of reduction in HSP27 band density of (PEG-DET/HSP27)/EV ternary mixtures ranged among the values noted in the case of PEG-DET/HSP27 and EV/HSP27 mixtures suggesting the competitive binding of negatively-charged EVs and HSP27 with the positively charged PEG-DET.

### 3.9. Physicochemical characterization of the HSP27 mixtures

Particle diameters and surface charge of the formed mixtures were measured using dynamic light scattering (**Fig. 8e-h**). The average diameter of native HSP27 protein was about 85 nm (**Fig. 8e**). The average zeta potential of native HSP27 (1 µg/mL in 10 mM HEPES buffer, pH 7.4) was about −9 mV suggesting that HSP27 exerts a net negative surface charge under physiological conditions (**Fig. 8f**). The average particle diameter of PEG-DET/HSP27 mixtures at 10:1 w/w ratio was over 1000 nm with a broad dispersity index. As the w/w ratio increased from 20:1 to 30:1, the particle diameter significantly (p<0.05) decreased from about 359 nm to 205 nm (**Fig. 8e**). PEG-DET/HSP27 mixtures showed a unimodal size distribution in the intensity plots (**Fig. S21a**). PEG-DET/HSP27 mixtures at 10:1 w/w ratio shifted the zeta potential of the native HSP27 protein from −9 mV to +10 mV confirming the electrostatic interactions of PEG-DET with HSP27 (**Fig. 8f**). The zeta potentials, however, did not change for w/w ratios of 20:1 and 30:1 (**Fig. 8f**) suggesting that the excess polymer do not bind to HSP27 protein.

sEV and m/lEV showed diameters of about 136 and 225 nm, respectively (**Fig. 8e**). sEV showed a bimodal particle size distribution compared to m/lEVs (**Fig. S21c,e**). sEV/HSP27 or m/lEV/HSP27 mixtures at a 10:1 w/w ratio did not change the particle diameter compared to naïve EVs (**Fig. 8e**). The zeta potential of m/lEV/HSP27 mixtures shifted towards near-neutral values compared to naïve m/lEVs and native HSP27 protein indicating (**Fig. 8f)**.

Next, PEG-DET was mixed with sEVs and m/lEVs at PEG-DET/EV w/w ratios 2:1 and 3:1, and the resulting changes in particle sizes and zeta potentials were compared with naïve sEVs and m/lEVs (**Fig. 8g,h**). The incubation of PEG-DET to sEV showed a considerable increase in particle size from 134 nm to 245 nm (**Fig. 8g**). The shift in mean zeta potential from −1.55 mV to −0.18 mV suggested the electrostatic interactions of PEG-DET and sEV (**Fig. 8h**). Furthermore, the z-average particle diameter of (PEG-DET/HSP27)/sEV significantly (p<0.01) increased to about 400 nm with a near-neutral zeta potential suggesting the interactions of PEG-DET/HSP27 and sEVs (**Fig. 8g,h**). A similar trend was observed for m/lEVs where the particle diameter gradually and significantly (p<0.0001) increased from naïve m/lEVs (216 nm) to PEG-DET/m/lEV (431 nm) to (PEG-DET/HSP27)/m/lEV (991 nm) (**Fig. 8g**). A shift in zeta potential was observed from −4 mV for naïve m/lEV to 1.29 mV for PEG-DET/m/lEV, and 0.75 mV for (PEG-DET/HSP27)/m/lEV confirming the electrostatic interactions of PEG-DET and m/lEV (**Fig. 8h**). In addition, an increase in PEG-DET to EV w/w ratio from 2:1 to 3:1 showed a reduction in z-average diameter (**Fig 8g**). A slight decrease in zeta potential at PEG-DET/EV at w/w 3:1 suggested that increasing PEG-DET amounts may increase the extent of interactions with EVs (**Fig. 8h**). The representative distribution plots of PEG-DET/EVs and (PEG-DET/HSP27)/EVs were shown in **Fig. S22**.

To summarize, PEG-DET/HSP27 mixtures showed a w/w ratio-dependent decrease in particle diameter and an overall positive surface charge. EV/HSP27 mixtures showed physicochemical characteristics similar to naïve EVs. The observed changes in particle diameter and zeta potential of (PEG-DET/HSP27)/EVs confirmed the interactions of EVs with PEG-DET/HSP27 mixtures.

### 3.10. PEG-DET/HSP27 and EV/HSP27 mixtures were cytocompatible with primary HBMECs

We performed an ATP assay to determine the cell viability of primary HBMECs treated with native HSP27, m/lEV, or sEV/HSP27 and PEG-DET/HSP27 mixtures. The cell viability of treatment groups was calculated using *Equation 1*. The average viability of cells treated with native HSP27 was 108.7% and there were no significant (p>0.05) differences between control, untreated cells, and native HSP27-treated groups suggesting that native HSP27 at 2 µg/well was well tolerated by HBMECs under normoxic conditions for 72 h (**Fig. S23**). The cell viability of HBMECs increased significantly (p<0.01) when treated with sEV/HSP27 (124.7%), m/lEV/HSP27 (123.1%), and sEV+m/lEV/HSP27 (116.8%) mixtures at 10:1 w/w ratio compared to untreated cells suggesting that EV/HSP27 mixtures were well tolerated by HBMECs for 72 h (**Fig. S23**). The EV/HSP27 mixture-mediated increase in cell viability can be correlated with a significant increase in HBMEC ATP levels that was observed when cells were treated with naïve sEV (121%), m/lEV (126.4%), or sEV+m/lEV (120.1%) at amounts equivalent those present in the EV/HSP27 mixtures. HBMECs treated with PEG-DET/HSP27 mixtures at w/w 20:1 showed an average cell viability of 100.4% suggesting that PEG-DET/HSP27 mixtures were cytocompatible. Free PEG-DET polymer was also well tolerated by HBMECs for 72 h. Polyethyleneimine, a positive control, at 50 and 100 µg/mL concentrations showed a significant (p<0.0001) reduction in HBMEC viability indicating that the assay was responsive to the toxicities.

### 3.11. Exogenous HSP27 attenuated the hypoxia-induced increase in tight junction permeability in primary HBMECs

#### 3.11.1. Paracellular permeability of <u>4.4 kD TRITC-Dextran (a small molecule tracer)</u> in pretreated HBMEC culture inserts

We measured the paracellular permeability of 4.4 kD TRITC-Dextran to evaluate the effect of (PEG-DET/HSP27)/EV and EV/HSP27 pre-treatment on the diffusion of a small molecule-mimic across primary HBMECs during normoxic, OGD, and OGD/reperfusion conditions. HBMECs were treated at a dose of 2 µg/well HSP27 mixed with PEG-DET at a 20:1 w/w ratio and with sEV, m/lEV, EVs (sEV: m/lEV=1:1) at a 10:1 w/w ratio. The total amount of EV in EV/HSP27 mixtures was 20 µg EV protein per insert for all the permeability assays. Native HSP27, free PEG-DET, and naïve sEV, m/lEV, and EVs were used as controls. The difference in the relative diffusion of 4.4 kD dextran between untreated and HSP27 mixtures-treated cells during normoxic conditions and pre-OGD phase was not statistically significant (p>0.05) (**Fig. S24a**). During the OGD phase, native HSP27, PEG-DET/HSP27, and (PEG-DET/HSP27)/EV-treated HBMECs showed a significant (p<0.05) reduction in the rates of 4.4 kD dextran diffusion compared to OGD, and free PEG-DET-treated HBMECs for 4 h (**Fig. 9a**). Besides, naïve sEV, m/lEV, EV, and their HSP27 mixture-treated HBMECs showed a significant (at least p<0.001) reduction in paracellular permeability for 4 h (**Fig. 9a**) of OGD exposure. sEV, m/lEV, and EV/HSP27 mixtures showed a consistent and significant (p<0.001) reduction in the relative diffusion of 4.4 kD dextran for 24 h of OGD exposure (**Fig. S24b**).

**Fig. 9.**
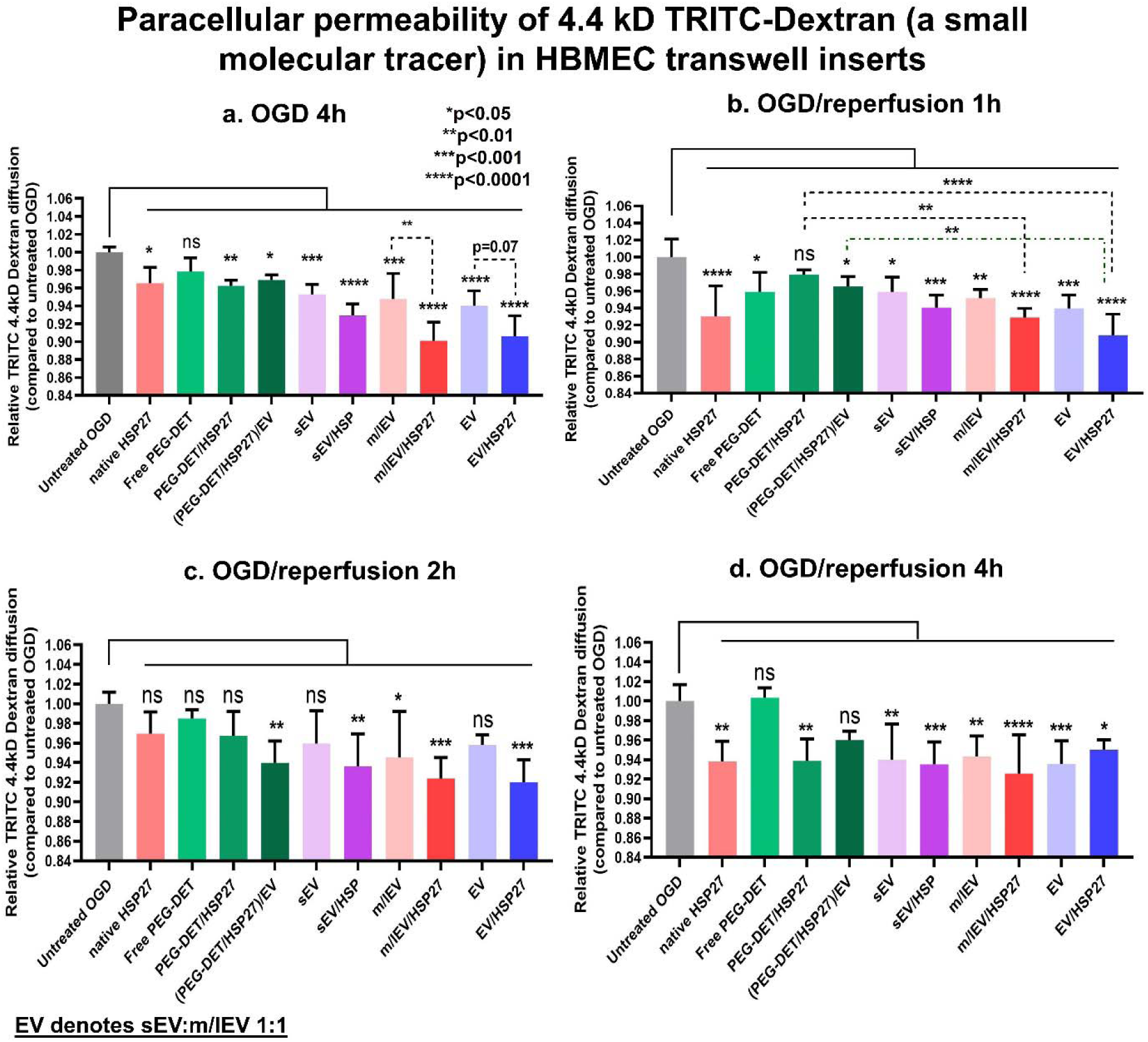
Paracellular permeability of 4.4 kD TRITC-Dextran under OGD, and OGD/reperfusion conditions in pretreated HBMEC transwell culture inserts. HBMECs were seeded in 24-well plates and maintained in a 37°C humidified incubator for a week. The complete growth medium was replaced with 300 µL of growth media containing indicated treatment groups for 72 h. Post-treatment, the treatment media was replaced with 300 µL of OGD medium containing 1 µM 4.4 kD TRITC-Dextran for 24 h. The abluminal chamber was filled with 0.5 mL of complete growth medium. Control, untreated cells were incubated in a complete growth medium in a humidified incubator whereas OGD treatment groups were incubated in an OGD chamber. At 4 h post-OGD **(a)**, a 500 µL volume was collected from the abluminal chamber and a fresh medium was added to the transwell inserts. Post-OGD treatment, HBMECs were washed with PBS and incubated with 300 µL of complete growth medium containing 1µM 4.4 kD TRITC-Dextran and incubated in a humidified incubator for 1-24h. At each time point, a 500 µL volume was collected from the abluminal chamber and fresh medium was added to the transwell inserts. The concentration of 4.4 kD TRITC-Dextran was measured at 1 h (OGD/reperfusion 1h, **b**), 2 h (OGD/reperfusion 2h, **c**), and 4 h (OGD/reperfusion 4h, **d**) using a Synergy HTX multimode plate reader at 485/20 nm excitation and 580/50 nm emission settings. The relative diffusion of TRITC 4.4 kD Dextran at each time point was determined by calculating the ratio of [TRITC-Dextran] in the abluminal compartment of treatment groups to that of untreated OGD control. Data represent mean±SD (n=4). * p<0.05, ** p<0.01, *** p<0.001, **** p<0.0001, ns: non-significant.

During the first hour of ischemia/reperfusion (OGD/reperfusion), HBMECs exposed to native HSP27 and (PEG-DET/HSP27)/EV ternary mixtures showed a significant (p<0.05) reduction in 4.4 kD dextran relative diffusion compared to OGD and free PEG-DET polymer-treated cells (**Fig. 9b**). In addition, naïve sEV, EV, sEV/HSP27, m/lEV/HSP27, and EV/HSP27 pretreated HBMECs showed a highly significant (p<0.001) decrease in the relative diffusion of 4.4 kD dextran compared to control HBMECs (**Fig. 9b**). sEV/HSP27, m/lEV/HSP27, and EV/HSP27 mixtures treatment resulted in a consistent and significant reduction in the paracellular permeability of 4.4 kD dextran for 2, 4, and 24 h of reperfusion compared to the OGD control (**Fig. 9c,d, and S24c**). In contrast, the relative diffusion in other treatment groups at 2, 4, and 24 h of OGD/reperfusion was non-significant compared to the untreated cells (**Fig. 9c,d, and S24c**). It should be noted that the sEV/HSP27, m/lEV/HSP27, and EV/HSP27 mixture-mediated decreases in 4.4 kD diffusion were not only retained for prolonged OGD/reperfusion times compared to (PEG-DET/HSP27)/EV (24 h *vs.* 2 h) but also with a significantly (p<0.05) greater magnitude (**Fig. 9** and **Fig. S24b,c**).

#### 3.11.2. Paracellular permeability of <u>65-85 kD TRITC-Dextran (a large molecule tracer)</u> in pre-treated HBMEC culture inserts

The effect of HSP27 mixed with PEG-DET and EVs on the paracellular permeability of 65-85kD TRITC-Dextran was evaluated as previously described for the small, 4.4 kDa tracer (section 3.10.1). Here, 65-85kD TRITC-Dextran was used as a large molecular mass tracer simulating the diffusion of large molecules such as proteins through the damaged BBB ^9^. Prior to subjecting cells to OGD (pre-OGD) (**Fig. S24d**), cells treated with sEV/HSP27 and m/lEV/HSP27 mixtures showed a significant (p<0.05) reduction in TRITC-Dextran permeability compared to control, untreated cells. In addition, naïve sEV, and EVs showed a considerable decrease in 65-85 kD TRITC-Dextran permeability (**Fig. S24d**). The data suggested that naïve sEV and m/lEV and their HSP27 mixtures may protect the barrier properties of BEC tight junctions compared to untreated cells under normoxic conditions. Interestingly, cells treated with native HSP27, PEG-DET/HSP27, and (PEG-DET/HSP27)/EV mixtures did not show any change in the TRITC-Dextran relative diffusion under normoxic conditions. The observed effects of decreased permeability were selective only for the EV-treated cells.

HBMECs pretreated with native HSP27, PEG-DET/HSP27 and (PEG-DET/HSP27)/EV mixtures showed a significant (p<0.01) reduction in 65-85 kD TRITC-Dextran relative diffusion during 4 h OGD exposure (**Fig. 10a**). sEV/HSP27 and m/lEV/HSP27 mixtures showed a significant (p<0.0001) reduction in 65-85 kD dextran relative diffusion compared to untreated OGD cells. Naïve sEV, m/lEV, and EVs also showed a statistically significant (p<0.05) reduction in dextran permeability under OGD conditions compared to untreated, OGD control (**Fig. 10a**). The data suggested that PEG-DET/HSP27 and (PEG-DET/HSP27)/EV mixtures can limit the diffusion of large molecules post-ischemia. Importantly, naive sEV and m/lEV limited the dextran diffusion before and OGD conditions, whereas a mixture of HSP27 and EVs showed a synergistic effect on decreasing dextran diffusion during ischemia. Notably, there was no difference in the 65-85 kD TRITC-Dextran relative diffusion between control and HBMECs treated with HSP27 mixtures at 24 h of OGD exposure (**Fig. S24e**).

**Fig. 10.**
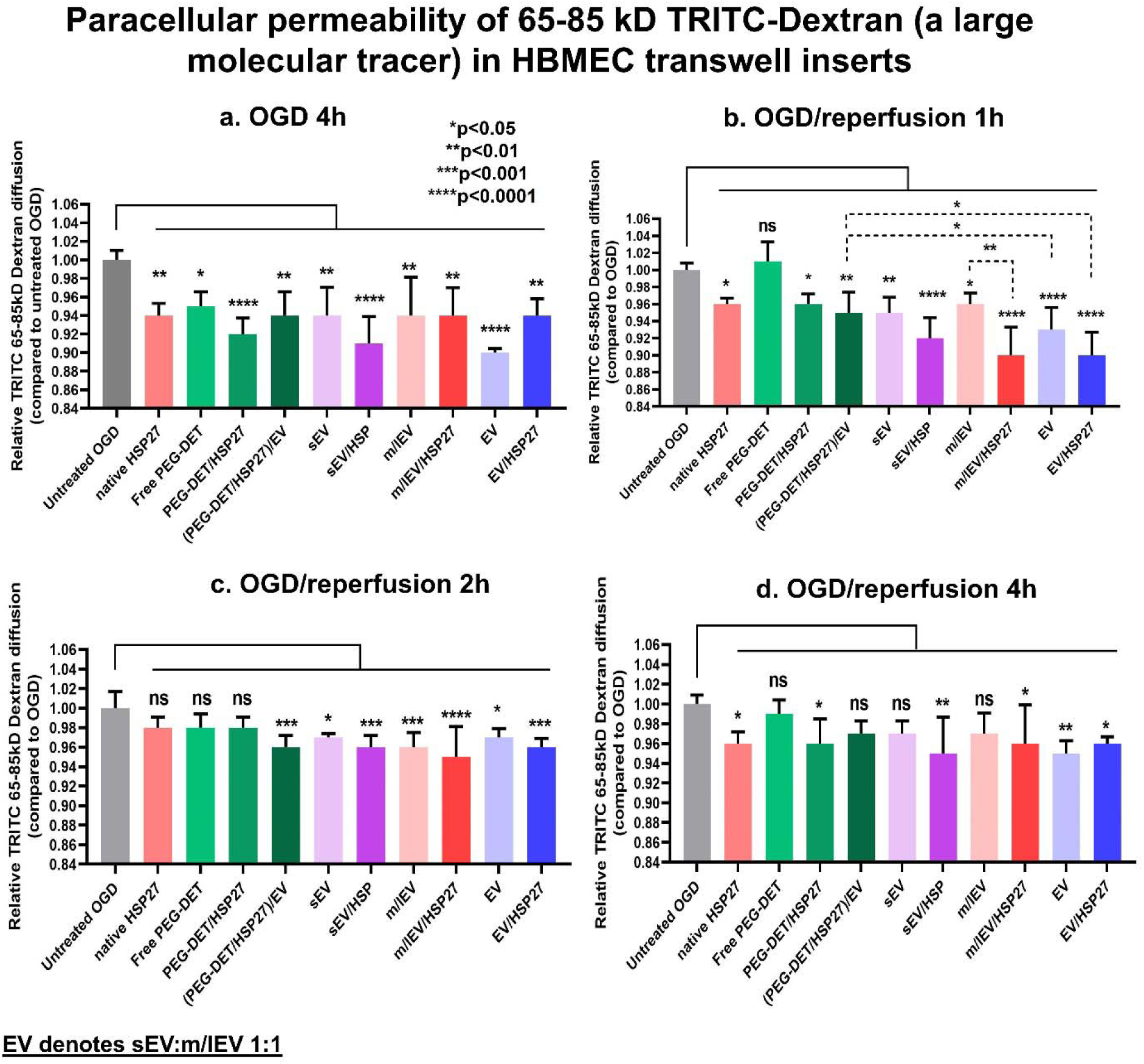
Paracellular permeability of 65-85 kD TRITC-Dextran under OGD, and OGD/reperfusion conditions in pretreated HBMEC transwell culture inserts. HBMECs were seeded in 24-well plates and maintained in a 37°C humidified incubator for a week. The complete growth medium was replaced with 300 µL of growth media containing indicated treatment groups for 72 h. Post-treatment, the treatment media was replaced with 300 µL of OGD medium containing 1 µM 65-85 kD TRITC-Dextran for 24 h. The abluminal chamber was filled with 0.5 mL of complete growth medium. Control, untreated cells were incubated in a complete growth medium in a humidified incubator whereas OGD treatment groups were incubated in an OGD chamber. At 4 h post-OGD **(a)**, a 500 µL volume was collected from the abluminal chamber and a fresh medium was added to the transwell inserts. Post-OGD treatment, HBMECs were washed with PBS and incubated with 300 µL of complete growth medium containing 1µM 65-85 kD TRITC-Dextran and incubated in a humidified incubator for 1-24 h. At each time point, a 500 µL volume was collected from the abluminal chamber and fresh medium was added to the transwell inserts. The concentration of 65-85 kD TRITC-Dextran was measured at 1 h (OGD/reperfusion 1h, **b**), 2 h (OGD/reperfusion 2h, **c**), and 4 h (OGD/reperfusion 4h, **d**) using a Synergy HTX multimode plate reader at 485/20 nm excitation and 580/50 nm emission settings. The relative diffusion of TRITC 65-85 kD Dextran at each time point was determined by calculating the ratio of [TRITC-Dextran] in the abluminal compartment of treatment groups to that of untreated OGD control. Data represent mean±SD (n=4). * p<0.05, ** p<0.01, *** p<0.001, **** p<0.0001, ns: non-significant.

Post-OGD, the OGD medium was replaced with fresh complete growth medium to evaluate the effects of HSP27 on ischemia/reperfusion-mediated diffusion of 65kD TRITC-Dextran. Exposure of cells to PEG-DET/HSP27 and (PEG-DET/HSP27)/EV mixtures showed a significant (p<0.05) reduction in relative diffusion 1 h post-ischemia/reperfusion (**Fig. 10b**). Importantly, sEV/HSP27, m/lEV/HSP27, and EV/HSP27 mixtures showed a significant (p<0.0001) reduction in dextran relative diffusion compared to OGD control after 1 h ischemia/reperfusion. Moreover, naïve sEVs and EVs also showed a significant (p<0.01) reduction in diffusion after reperfusion suggesting that naïve EVs increase BEC tight junction integrity immediately after ischemia/reperfusion. In addition, their mixtures with HSP27 synergistically reduce the large molecule infiltration across the BECs (**Fig. 10b**). Importantly, (PEG-DET/HSP27)/EV mixtures, sEV/HSP27, m/lEV/HSP27, and EV/HSP27 mixtures showed a significant (p<0.001) reduction in relative diffusion compared to OGD control 2 h post-OGD/reperfusion. sEV/HSP27, m/lEV/HSP27, and EV/HSP27 mixtures showed continual significant (p<0.01) reduction in 65-85 kD relative diffusion compared to OGD control for 4 h of reperfusion, whereas changes in the relative diffusion of dextran, were insignificant (p>0.05) in the other treatment groups compared to OGD control (**Fig. 10c**). It can be inferred that in addition to the naïve EV-mediated protection of tight junction integrity pre-OGD/normoxia, during OGD, and OGD/reperfusion, their mixtures with HSP27 can protect BECs during OGD/reperfusion. There were no further differences in dextran relative diffusion amongst the OGD control and treatment groups during 4-24 h of ischemia/reperfusion (**Fig. 10d** and **S24f**).

## 4. Discussion

This study aimed to evaluate the effects of the innate EV mitochondria and EV/HSP27 mixtures on the tight junction integrity and metabolic function of ischemic BECs. This one-two-punch strategy allowed us to harness the innate EV mitochondria to increase cellular bioenergetics and mixtures of EV/exogenous HSP27 protein to reduce paracellular permeability in ischemic BECs. This novel approach to protecting the ischemic BECs can potentially limit damage to the BBB, an integral component of the neurovascular unit. Loss of mitochondrial function increases BBB permeability leading to a secondary injury that exacerbates post-stroke damage ^44^. Protection of the BBB can limit early structural damage and prevent chronic neurological dysfunction post-stroke ^9, 44–48^.

The present study investigated (1) the effects of the innate EV mitochondria on the bioenergetics of recipient HBMECs under normoxic and ischemic conditions, and (2) the effects of EV/HSP27 mixtures on the paracellular permeability of tracer molecules across ischemic BEC monolayers. The results of our studies demonstrated that EV mitochondria, specifically m/lEVs, transferred into and colocalized with the mitochondrial network of the recipient BECs. As a result, EV-treated primary HBMECs demonstrated increased intracellular ATP levels and mitochondrial respiration. Prophylactic treatment of EV/HSP27 mixtures and (PEG-DET/HSP27)/EV ternary mixtures significantly reduced ischemia-induced paracellular permeability of small and large tracer molecules across primary HBMEC monolayers.

BECs form the first layer of the BBB, and they contain about two to five-fold greater mitochondrial content compared to peripheral endothelial cells ^49^. BEC-derived EVs contain BBB receptors such as transferrin and insulin that enable EVs to cross the BBB for the treatment of various neurovascular disorders ^50–52^. A greater mitochondrial load and their natural affinity for BBB targeting motivated us to isolate sEVs and m/lEVs from the hCMEC/D3 BEC cell line. We used a differential ultracentrifugation protocol, the most commonly used EV isolation method ^53^, to isolate m/lEVs and sEVs from the EV-conditioned medium. Post-ultracentrifugation and resuspension, sEVs and m/lEVs showed their characteristic particle diameters (100-250 nm, **Fig. 1a**) that largely aligned with the previous reports ^19, 23, 34, 54, 55^.

We have observed an apparent discrepancy between DLS and TEM sizing of sEVs and m/lEVs. For sEV, TEM analysis showed a smaller particle size than DLS measurements, whereas TEM sizing of m/lEVs was larger compared to DLS. It is important to note that sEVs and m/lEVs are heterogenous EV subpopulations with polydispersity indices ∼ 0.4 (**Fig. 1a**). Therefore, the observed particle size differences of EVs measured by different techniques could be attributed to significant differences in the operating principles and sample handling/processing conditions. DLS estimates hydrodynamic particle diameters in a solvent, and the hydrodynamic diameter includes the core of the particle and the liquid layer surrounding the lipid layer. On the other hand, TEM measures the core size of an individual particle in a dried state under a vacuum. The environmental conditions may shrink the particles due to the osmotic diffusion of solvent from the aqueous core to the surrounding space. Importantly, TEM represents the morphology of only a small portion of the sample, which can further be affected by operator bias.

We also compared the particle size distribution of sEV and m/lEV obtained from DLS and NTA. Although DLS and NTA are commonly used for particle size measurements, both methods technically differ in weighing the particle size distribution (PSD). DLS reports intensity-weighted particle size distribution, whereas NTA reports number-weighted distribution ^56^. The differences between the number-, volume-, and intensity-average distribution extracted from the DLS measurements are shown in **Fig. S25a**. It should be noted that there is a significant (at least p<0.001) difference between sEV and m/lEV particle diameters irrespective of the type of representation: number, volume, or intensity -averages (Fig. S3b, colored lines). As expected and consistent with published reports, m/lEVs showed a larger particle diameter than sEV. In contrast, NTA did not show any statistical difference between sEV and m/lEV particle diameter; instead, it showed a slightly larger sEV particle diameter than m/lEV (**Fig. S25b**).

Anionic phospholipid components such as phosphatidylinositol, phosphatidylserine, and glycosylated lipid derivatives exert a net negative zeta potential on the sEVs and m/lEVs membranes (**Fig. 1a**) ^57, 58^. The broad polydispersity indices demonstrate the natural heterogeneity of both EV sub-populations. Similar to cells and other biomolecules, it is important to preserve EV physicochemical characteristics and biological activities during storage conditions that critically determine the scope of their therapeutic application. In our studies, sEVs and m/lEVs isolated from a conditioned medium retained their particle diameters, dispersity indices, and zeta potential after three consecutive freeze-thaw cycles (samples were frozen at −20°C for 24 h and then were thawed at room temperature for 1 h, **Fig. S1c-h**). Jeyaram *et al.* showed that EVs isolated from biofluids such as blood, milk, urine, and conditioned medium preserved their physical and functional properties stored at −80°C compared to 4° and − 20°C ^59, 60^. Moreover, freeze-thaw cycles at −20° and −80°C did not affect the stability of plasma exosome miRNA ^61^.

Our TEM analysis demonstrated that m/lEVs, but not sEVs, contained mitochondria (**Fig. 2c,d**). Mitochondria-rich m/lEVs structures were consistent with published reports ^33–36^. Prior studies have demonstrated that mitochondria or mitochondrial components such as mtDNA, and mitochondrial proteins were secreted into the extracellular milieu and transferred between cells ^34, 36, 62–67^. For instance, mesenchymal stem cell-derived-m/lEVs transferred mitochondria into the recipient macrophages leading to increased cellular bioenergetics ^34^. Guescini *et al.* demonstrated that exosomes were released from glioblastoma and astrocytes transferred mtDNA from glioblastoma to astrocytes ^68^.

We confirmed the presence of ATP5A protein (a subunit of mitochondrial adenosine triphosphate synthase complex ^23, 69^) in m/lEVs using western blotting (**Fig. 2e**) as we did in our previous studies ^19, 23^. ATP5A plays important role in mitochondrial ATP production by catalyzing the synthesis of ATP from ADP in the mitochondrial matrix during oxidative phosphorylation ^70^. A considerably higher ATP5A band density in m/lEVs compared to sEVs suggested that m/lEVs contain a greater mitochondrial load compared to sEVs. D’Acunzo *et al.* demonstrated the presence of ATP5A in EVs isolated from mice brains using mass spectroscopy-based proteomic analysis ^71^. Sanson *et al.* also reported ATP5A expression as a mitochondrial marker in stromal cell-derived EVs using western blot analysis ^72^. We performed western blotting in sEVs and m/lEVs lysates to determine the presence of TOMM20, a 16 kDa outer mitochondrial membrane protein, as an additional mitochondrial marker (**Fig. 2e**). TOMM20 is a translocase of the outer membrane receptor that regulates the import of specific proteins from the cytosol ^73^. The selective presence of TOMM20 in m/lEVs and cell lysates, but not in sEVs lysates further confirmed that m/lEVs contain mitochondria. Silva *et al*. also demonstrated the presence of TOMM20 in mesenchymal stromal cell-derived, mitochondria containing extracellular vesicles ^74^. sEVs showed the presence of CD9 (**Fig. 2e**), a tetraspanin marker associated with exosomal cargo selection, binding, and uptake of sEV by target cells ^75^. Collectively, our TEM images of sectioned EVs and the presence of TOMM20 in m/lEVs indicated the presence of mitochondria in m/lEVs, whereas sEVs contain mitochondrial proteins but not entire mitochondria.

We isolated sEVs and m/lEVs from the conditioned medium of Mitotracker deep red (MitoT-red) pre-stained hCMEC/D3 cells. MitoT-red is a mitochondrion membrane potential-dependent carbocyanine dye that selectively stains polarized mitochondria, and its fluorescence intensity is reduced during mitochondrial depolarization ^76, 77^. We demonstrated that m/lEVs contain a greater mitochondrial load compared to sEVs and are colocalized with recipient BECs mitochondria (**Fig. 3-5 and Fig. S11-17**). m/lEV-mitochondria were transferred into the recipient BECs within 24 h of incubation. Increasing the m/lEV doses and incubation times significantly increased uptake into the recipient BECs. Importantly, the m/lEV-mitochondria efficiently colocalized with the mitochondria network in the recipient BECs. The co-localization of m/lEVs and recipient BEC mitochondria was confirmed by the presence of overlapping signals of the EV-mitochondria fluorescence signals with the recipient BEC mitochondria signals. We used two orthogonal approaches to stain the mitochondrial network in the recipient BECs: Mitotracker green and the CellLight mitochondria-GFP BacMam technique (**Fig. 5, and Fig. S17**). The carbocyanine Mitotracker green dye stains the functional mitochondria whereas CellLight Mitochondria-GFP BacMam comprising a fusion construct of α-pyruvate dehydrogenase and emGFP packaged in the baculoviral vector stains a structural mitochondrial matrix protein (α-pyruvate dehydrogenase) ^25, 37^. Thus, utilizing two orthogonal types of staining techniques, we demonstrated an efficient colocalization of polarized EV mitochondria with the polarized mitochondria in the recipient BECs (via Mito-T-green staining, **Fig. 5**) and colocalization of functional, polarized EV mitochondria with the structurally intact mitochondria in the recipient BECs (via CellLight Mitochondria-GFP staining, **Fig. S17**). m/lEVs and sEVs showed a dose-dependent increase in colocalization at 72 h; specifically, the m/lEVs demonstrated a significantly greater colocalization coefficient compared to sEVs (**Fig. 5b**). The selective mitochondrial packaging into BEC-derived m/lEVs compared to sEVs was consistent with published reports ^34, 66, 68^. Overall, our data demonstrated the m/lEV-mediated transfer of functional mitochondria and their colocalization with the recipient BEC mitochondrial network.

One of the main functions of mitochondria is to synthesize ATP from ADP during mitochondrial aerobic respiration, and therefore, we measured relative ATP levels in the recipient BECs treated with sEVs or m/lEVs using a luciferase-based ATP assay. Our results demonstrated that naïve sEVs and m/lEVs mediate a dose-dependent significant increase in the relative ATP levels at 48 h-post incubation (**Fig. 6a-c**). Importantly, m/lEVs outperformed sEVs in increasing recipient cell viability and the effects persisted for 72 h-post incubation (**Fig. 6c**). Islam *et al.* reported that mitochondria containing m/lEVs derived from bone marrow stromal cells increased ATP levels of alveolar epithelial cells ^63^. Guo *et al*. demonstrated that the transfer of mitochondria isolated from donor bone marrow-derived mesenchymal cells (BMSC) into the recipient BMSCs increased cellular ATP production, proliferation, and migration, and repaired bone defects *in vitro* and *in vivo* ^78^.

Numerous studies have demonstrated ischemia-induced cerebral endothelial dysfunction/apoptosis and BBB breakdown ^79–82^. In our studies, BECs exposed to OGD exposure in a hypoxic chamber led to about 60% endothelial cell death at 24 h compared to untreated cells (**Fig 6d)**. The observed data is consistent with the published reports ^83, 84^. Our results showed that primary HBMECs treated with naïve sEVs and m/lEVs resulted in a four to five-fold increase in endothelial ATP levels compared to control, untreated cells (**Fig. 6d**). Importantly, EV-mediated increases in ATP levels were dose-dependent and m/lEVs outperformed sEVs in rescuing the ATP levels and consequently the survival of ischemic HBMECs 24 h post-OGD (**Fig. 6d**). Importantly, m/lEVs isolated from rotenone- and oligomycin-exposed BECs showed a loss of RTN-m/lEV and OGM-m/lEV mitochondria functionality to a much greater extent than RTN-sEVs and OGM-sEVs, respectively (**Fig. 6f**). It is likely that the rotenone and oligomycin-mediated inhibitions of mitochondrial complexes I and V in donor hCMEC/D3 BECs affected the functional mitochondrial load in m/lEVs, hence, RTN-m/lEVs and OGM-m/lEVs showed a dramatic and complete loss of m/lEV-mediated increase in ATP levels. Besides, consistent with our prior results, sEVs lack entire mitochondria, and therefore, rotenone-and oligomycin-mediated inhibitions of mitochondrial complex I in donor cells minimally affected sEV functionality.

We investigated the effects of m/lEV and sEV treatment on mitochondrial respiration and glycolytic capacity function in the recipient BECs using a Seahorse setup. The state-of-art Seahorse extracellular flux (XF) analyzer allows real-time analysis of extracellular acidification rate (ECAR), an indicator of glycolysis, and the oxygen consumption rates (OCR), an indicator of mitochondrial respiration in live intact cells ^85, 86^. We demonstrated that m/lEVs resulted in increased OCR compared to sEVs suggesting that m/lEV-mediated significant increase in mitochondrial respiration may likely be due to their innate mitochondrial load—including mitochondria and mitochondrial proteins (**Fig. 6i**). We also noted that m/lEVs showed a significantly (p<0.05) greater increase in recipient BEC glycolysis capacity compared to sEVs (**Fig. 6j**). Importantly, Phinney *et al.* demonstrated that MSC-derived EVs significantly increased maximum OCR in the recipient macrophages compared to controls in MSC-macrophage cocultures ^34^. Overall, through the use of these orthogonal tools (microscopy studies, ATP, and Seahorse assays), we have demonstrated that m/lEVs contain functional mitochondria compared to sEVs.

In a pilot experiment, we determined that intravenously injected mitochondria-containing m/lEVs showed neuroprotection in a mouse model of stroke. In our previous work, superoxide dismutase (SOD1) protein formulated with cationic polymers (nanozymes) injected *i.v.* during the onset of reperfusion in a rat middle cerebral artery occlusion (MCAo) model showed a reduction in infarct volume compared to saline ^15, 87, 88^. It should be noted that here we administered m/lEVs two hours post-ischemia/reperfusion in an effort to simulate delayed administration that occurs in clinical stroke scenarios and still demonstrate a ca. 40% reduction in infarct volume compared to vehicle-injected mice. These results from this pilot study provide proof of concept to advance our studies. It is important to note that this is the <u>first</u> demonstration of *in vivo* protective effects of m/lEVs. A larger number of mice will be utilized to confirm therapeutic efficacy in male and female mice (n=12 mice/group/sex) where, we will perform efficacy studies in mice, we will access infarct volumes at 72 hours, and will also analyze behavioral recovery using neurologic deficit scoring and corner tests in treated mice.

We used EVs and PEG-DET to formulate HSP27 mixtures. The isoelectric point of human recombinant HSP27 is 5.89 ^43^, therefore, it exerts a net negative charge at physiological pH 7.4 (**Fig. 8f**). The diethyltriamine side chain of PEG-DET cationic diblock copolymer has two pKa values associated with its molecular conformation (*gauche vs. anti*). The *gauche* conformation of DET exerts a pKa of 9.9 that induces the formation of stable mixtures with negatively charged polynucleotides at physiological pH 7.4 ^15, 89^. We confirmed the formation of PEG-DET/HSP27 mixtures using native PAGE followed by Coomassie staining (**Fig. 8a,b**) and dynamic light scattering (**Fig. 8e,f**). Despite the negative surface charge of EVs and the native HSP27 protein at pH 7.4, EV/HSP27 mixtures showed about 20% interactions in native PAGE analysis (**Fig. 8c,d**). Haney *et al.* also reported that macrophage-derived exosomes incubated with bovine liver catalase protein did not affect the particle diameter and dispersity indices of the sEVs/catalase mixture ^90^. Notably, the zeta potential of EV/HSP27 mixtures was significantly different compared to native HSP27. The weaker interactions of HSP27 with EVs were particularly advantageous and avoided intrusive modes of HSP27 loading (such as sonication, freeze/thaw cycles, and saponin-mediated loading). Such intrusive modes of loading may damage the EV membrane integrity and inversely impact the functionality of innate EV cargo, specifically, their mitochondria. We further confirmed the m/lEV and HSP27 interactions using an immunoprecipitation pull-down assay (**Fig. S26**). We engineered ternary mixtures of EVs with PEG-DET/HSP27 mixtures at different w/w ratios to increase the HSP27 loading into hCMEC/D3-derived EVs. The positively-charged PEG-DET/HSP27 mixtures (+9 mV, **Fig. 8f**) formed electrostatic interactions with sEV and m/lEV which was confirmed by an intermediate HSP7 band density between PEG-DET/HSP27 and sEV/HSP27 mixtures (**Fig. 8c,d**) and increased resulting particle diameters (**Fig. 8g**). (PEG-DET/HSP27)/EV ternary mixtures showed a near-electroneutral zeta potential (**Fig. 8h**) that may allow longer systemic circulation *in vivo*. The inclusion of EVs in (PEG-DET/HSP27)/EV ternary mixtures may facilitate interactions with the BBB and mediate endothelial targeting ^91^ during *in vivo* delivery while the cationic PEG-DET can enhance the cellular uptake and facilitate the endosomal escape of HSP27 ^15^.

We evaluated the prophylactic (PEG-DET/HSP27)/EVs and EV/HSP27 treatment-induced protection of tight junction integrity by measuring the relative diffusion of hydrophilic tracers. We used fluorescent tracers varying in molecular mass: we used 65-85 kDa TRITC-Dextran, a large molecular weight tracer to simulate the infiltration of proteins and larger blood-borne molecules, and a 4.4 kDa TRITC-Dextran to simulate the diffusion of small molecules during ischemia/reperfusion.

We showed naïve EV, (PEG-DET/HSP27)/EV, and EV/HSP27-induced protection of tight junction integrity against the paracellular flux of small molecules under OGD and OGD/reperfusion conditions (**Fig. 9a-d**). (PEG-DET/HSP27)/EV ternary mixtures strengthened the tight junctions to restrict 4.4 kD dextran entry up until 4 h of OGD exposure and as long as 4 hours of OGD/ reperfusion (**Fig. 9a-d**). Importantly, naïve EVs and their HSP27 mixtures retarded small molecule permeability during OGD conditions and OGD/reperfusion. Notably, the magnitude of EV/HSP27 mixture-mediated protection is considerably greater than naïve EVs demonstrating a synergistic effect of EVs and HSP27 in increasing BEC tight junction integrity (**Fig. 9a-d**). In addition, HBMECs treated with (PEG-DET/HSP27)/EV ternary mixtures efficiently decreased 65-85 kD TRITC-Dextran relative diffusion for 4 h of OGD exposure followed by immediate two hours of reperfusion compared to untreated cells, native HSP27, and free PEG-DET-treated groups (**Fig. 10a-d**). These results indicated that (PEG-DET/HSP27)/EV-mediated efficient transfer of HSP27 into BECs restores the tight junction integrity and may protect the BBB during ischemia/reperfusion injury. Importantly, naïve EVs and EV/HSP27 mixtures-treated HBMECs showed efficient protection of tight junction integrity during OGD and the first hour of ischemia/reperfusion (**Fig. 10b**).

Interestingly, although ternary mixtures of (PEG-DET/HSP27)/EV showed prolonged protection of the tight junction integrity compared to native HSP27 and PEG-DET/HSP27 mixtures, the magnitude of protection is relatively lower than naïve EVs and EV/HSP27 mixtures. It is likely that the weak interactions between EV and HSP27 enable efficient EV and HSP27 uptake and allow exerting the maximum therapeutic potential. In contrast, the stronger electrostatic interactions in (PEG-DET/HSP27)/EV ternary mixtures may impede the release of HSP27 resulting in a slightly lower therapeutic effect compared to EV/HSP27 mixtures. sEV/HSP27, m/lEV/HSP/27, and m/lEV/HSP27 showed superior BEC tight junction protection compared to PEG-DET-based groups. The possible reasons for EV/HSP27 mixture-mediated superior BEC protection could be due to optimal binding and effective intracellular release of HSP27 in the case of EV/HSP27 in comparison to PEG-DET/HSP27. It should be noted that regardless of the small magnitude of decreases in relative permeabilities, EV/HSP27 mixtures show a statistically significant effect compared to the controls. It remains to be investigated if alternate engineering approaches, such as HSP27 protein loading into EVs using sonication ^92^, can further increase the magnitude of the observed effects. Our future works will optimize the process of protein loading into EVs.

## 5. Conclusion

This one-two-punch approach using EVs increased the BEC mitochondrial function due to the innate EV mitochondrial load, and EV/HSP27 protected tight junction integrity in ischemic BECs. Naïve m/lEVs and sEVs increased ATP levels (albeit m/lEV showed a greater magnitude of ATP increases), mitochondrial respiration, and glycolytic capacities in the recipient BECs. For the first time, *i.v.* injected m/lEVs showed potential for neuroprotection in a mouse model of ischemic stroke. (PEG-DET/HSP27)/EV and EV/HSP27 mixtures restored tight junction integrity in primary human BECs by limiting the paracellular permeability of small and large molar mass tracer molecules during ischemia/reperfusion injury. The outcomes of the present study indicate that this approach has the potential to protect the damaged BBB *in vivo* that in turn can ameliorate the long-term neurological damage and dysfunction in rodent models of ischemic stroke.

## Abbreviations

BBB: Blood-brain barrier
BECs: Brain endothelial cells
Calcein AM: Calcein acetoxymethyl
ECAR: Extracellular acidification rate
EVs: Extracellular vesicles
EXOs: Exosomes
FT cycle: Freeze/Thaw cycle
GAPDH: Glyceraldehyde 3-phosphate dehydrogenase
HSP27: Heat shock protein 27
hCMEC/D3: human cerebral microvascular endothelial cell line
HBMEC: primary human brain microvascular endothelial cells
MVs: Microvesicles
m/lEV: medium-to-large EVs
MitoT-red-EV: Mitotracker deep red-labeled extracellular vesicles
OCR: Oxygen consumption rate
OGD: Oxygen-glucose deprivation
OGD/RP: Oxygen-glucose deprivation/reperfusion
PEI: Polyethylenimine
PEG-DET: poly (ethylene glycol)-b-poly (diethyl triamine)
ROS: Reactive oxygen species
sEV: small EVs
TRITC: Tetramethyl rhodamine iso-thiocyanate

## Acknowledgments

This work was supported via start-up funds for the Manickam laboratory from Duquesne University (DU) and a 2021 Faculty Development Fund (Office of Research, DU) to the PI. We would like to acknowledge the Neurodegenerative Undergraduate Research Experience for funding DXD, MF and AS through a grant from the National Institute of Neurological Disorders and Stroke (R25NS100118). The authors are thankful to Drs. Lauren O’Donnell and Manisha Chandwani and Ms. Yashika Kamte (DU) for flow cytometry support.

