## Supplementary information for "Mitochondria-containing Extracellular Vesicles (EV) Reduce Mouse Brain Infarct Sizes and EV/HSP27 Protect Ischemic Brain Endothelial Cultures"

**†Corresponding author:**

Devika S Manickam, Ph.D.

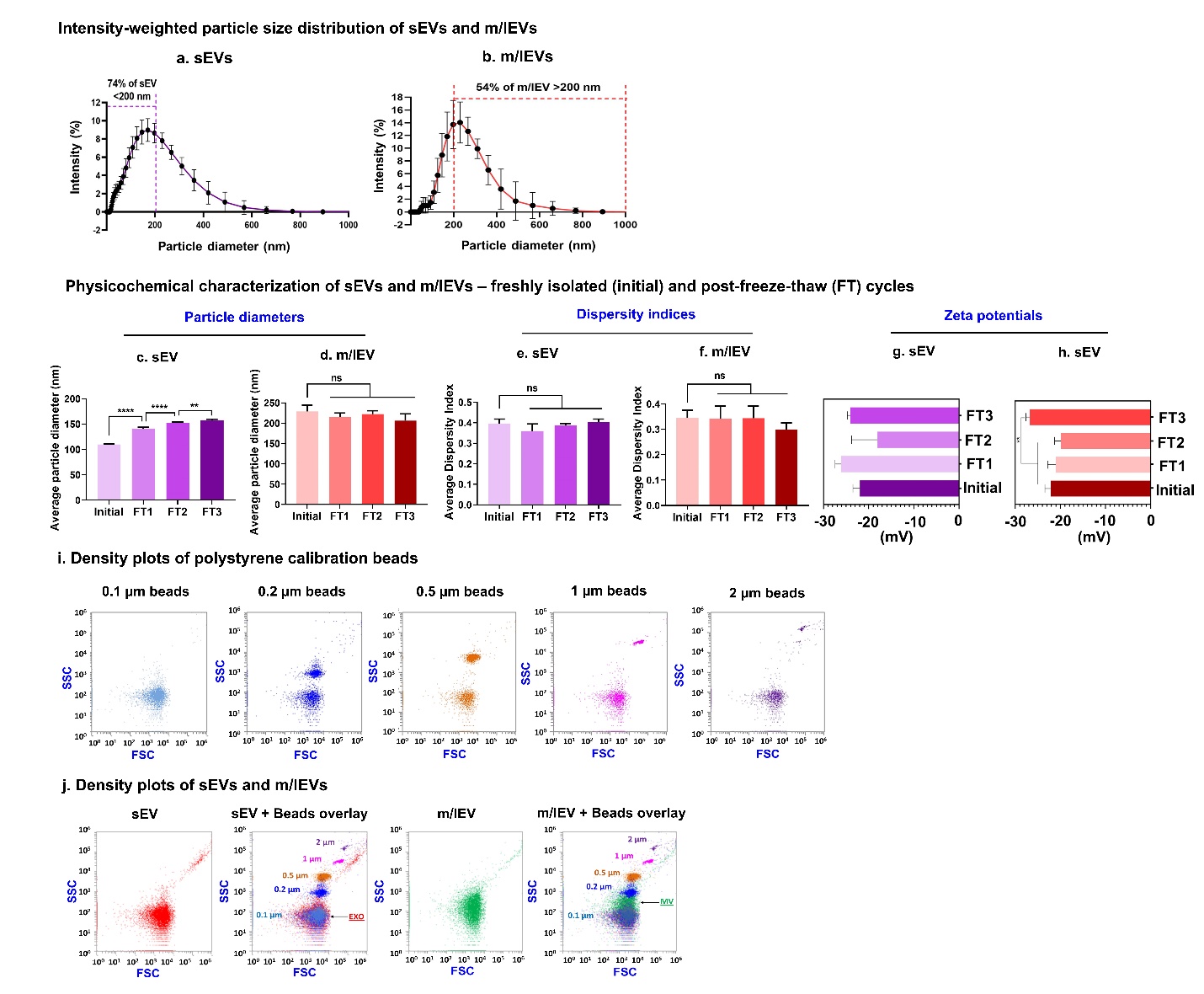

**Fig. S1. Physicochemical characteristics and stability of EVs during storage conditions.** (**a-b**) Intensity-weighted particle size distribution of freshly isolated sEVs and m/lEVs using dynamic light scattering. The percentages at the cut-off diameter of 200 nm were estimated from the Malvern Pro DLS raw data showing the distribution of particle diameters as a function of % scattered light intensity. Average particle diameters (**c,d**), and dispersity indices (**e,f**), and zeta potentials (**g,h**) of hCMEC/D3 BECs-derived EVs after three freeze-thaw (FT) cycles were determined using dynamic light scattering on a Malvern Zetasizer Pro-Red. Freshly-isolated samples (Initial) were used as controls. Samples were diluted to 0.5 mg protein/mL in 1x PBS for particle diameter and 10 mM HEPES buffer pH 7.4 for zeta potential measurements. The samples were stored at -20°C for 24 h and thawed at room temperature for 30 min prior to analysis. Data are presented as mean±SD of n=3 measurements. * p<0.05. The particle counts of polystyrene calibration beads (**i**), sEVs, and m/lEVs (**j**) were captured in forward scatter (FSC), side scatter (SSC) plots using Attune NxT flow cytometer. The histograms of calcein-positive events of intact sEVs and m/lEVs post-three FT cycles were detected using a small particle side scatter 488/10-nm filter in an Attune flow cytometer. Unstained EVs were used to gate the histograms for estimating percentage calcein-positive counts.

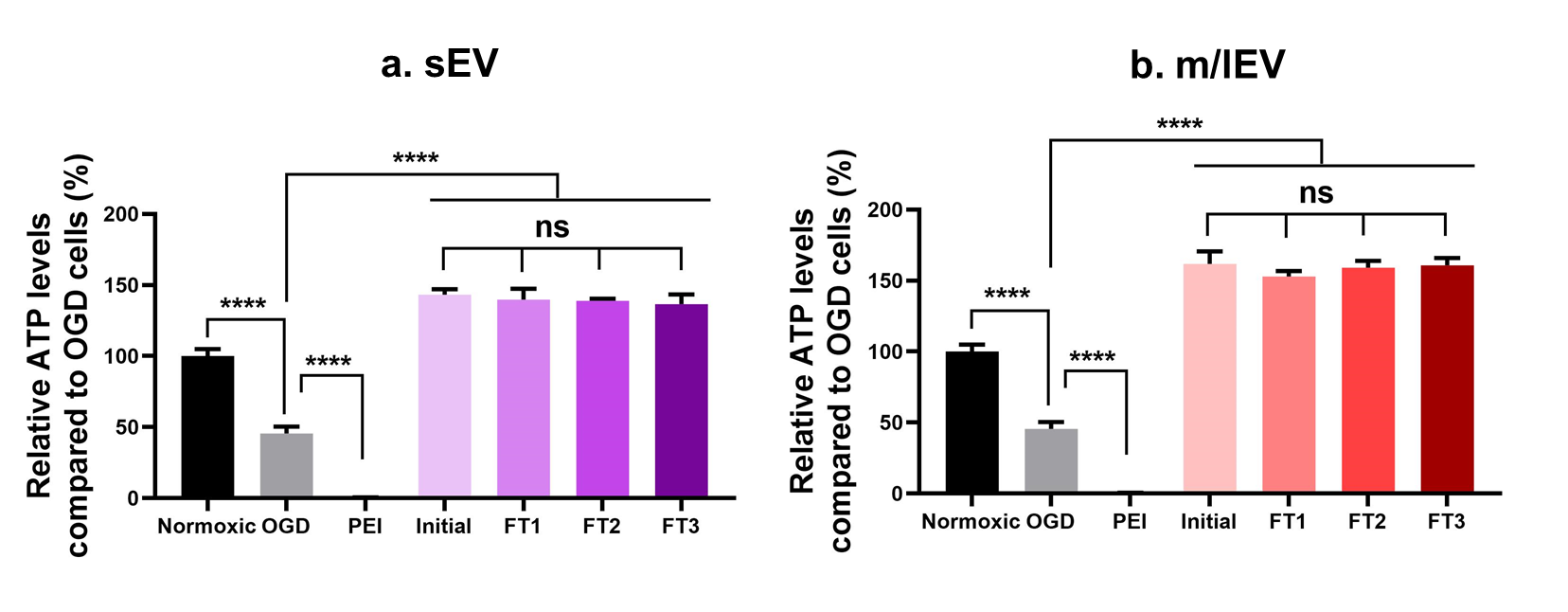

**Fig. S2. EVs retained their functionality during the storage conditions.** HBMECs were cultured in the 96-well plates until 80% confluency in complete growth medium in a 37 °C humidified incubator. Confluent monolayers were treated with sEV (**a**) and m/lEV (**b**) at 50 µg EV proteins in OGD medium for 24 h. Cells treated with OGD medium were used as OGD control, whereas cells treated with complete growth medium were used as the normoxic control. Post-treatment, cells were incubated with a 1:1 mixture of fresh growth medium and Cell titer Glo reagent. The relative luminescence units (RLU) were measured using a SYNERGY HTX multimode plate reader at 1s integration time. Relative ATP levels were calculated by normalizing the RLU of treatment groups to the RLU of OGD control. Data represent mean±SD (n=3).

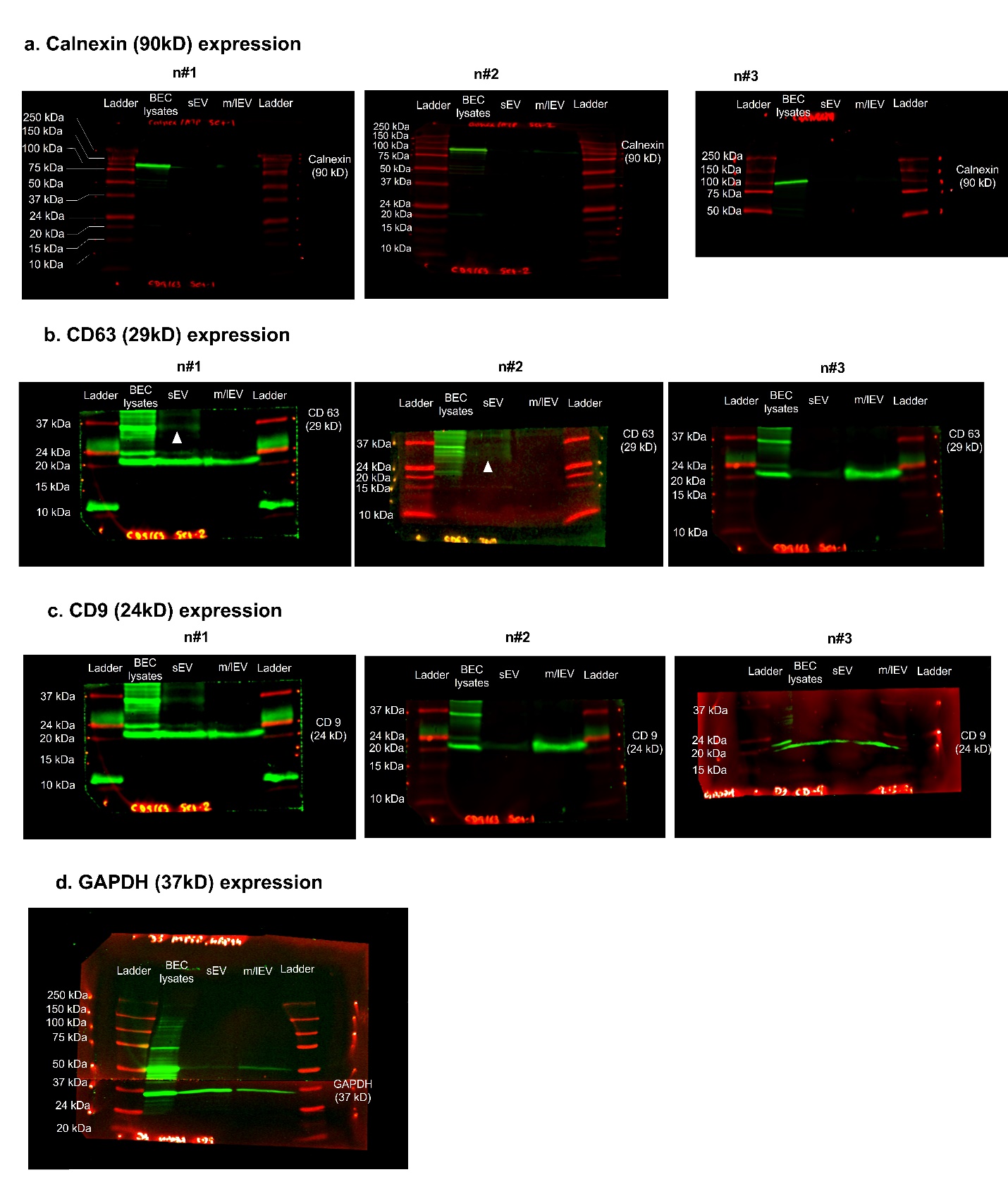

**Fig. S3. Uncropped western blots of calnexin, CD63, CD9, and GAPDH expression in sEVs and m/lEVs shown in Fig. 1d.**

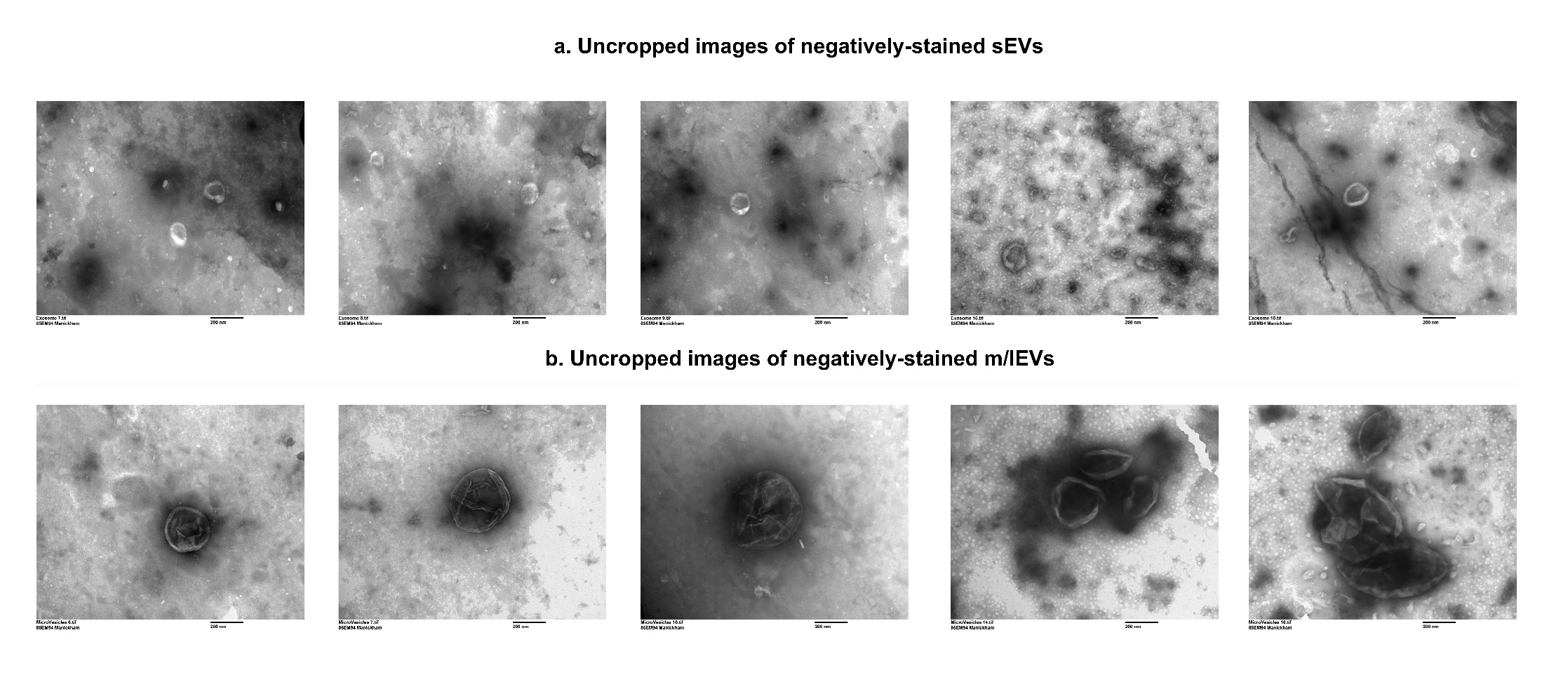

**Fig. S4. Uncropped TEM images of negatively stained sEVs (a) and m/lEVs (b) shown in Fig. 2a, b.**

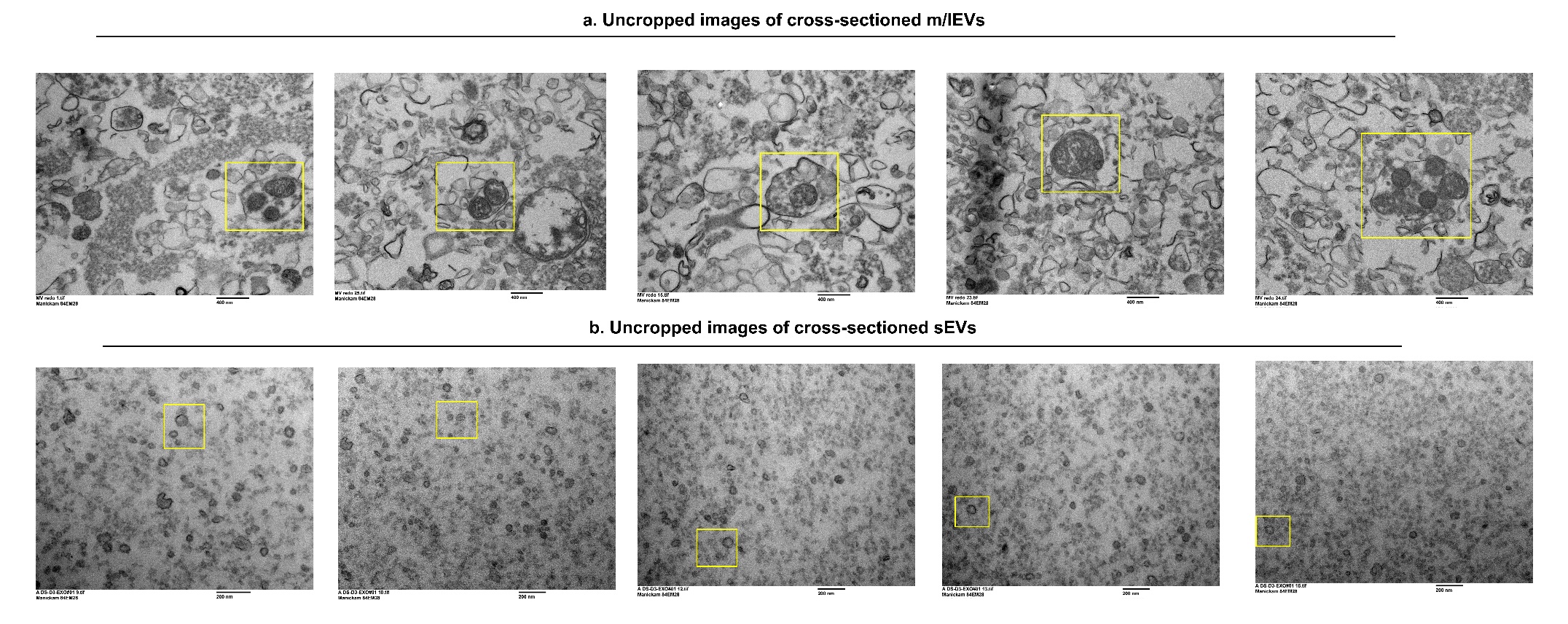

**Fig. S5. Uncropped TEM images of cross-sectioned m/lEVs (a) and sEVs (b) shown in Fig. 2c, d.**

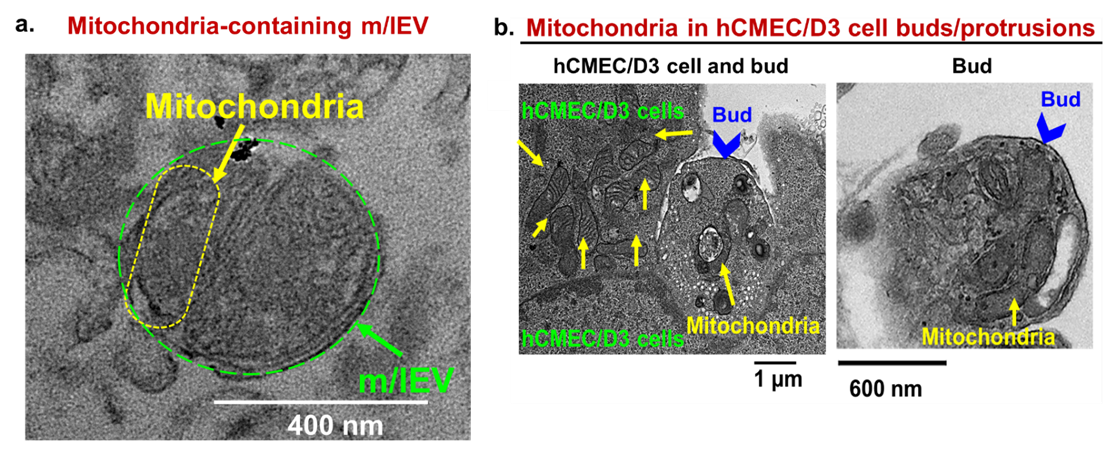

**Fig. S6. Transmission electron microscopy images of hCMEC/D3 cell buds/protrusions.** hCMEC/D3 monolayers were cultured in a 6-well plate. The cells were washed with 1x PBS and fixed with 2.5% glutaraldehyde in PBS overnight. The cell suspension was pelleted, and ultrathin 70 nm sections were imaged using a JEOL JEM 1400 Plus transmission electron microscope. The blue arrowheads indicate hCMEC/D3 cell buds containing mitochondria (yellow arrowheads).

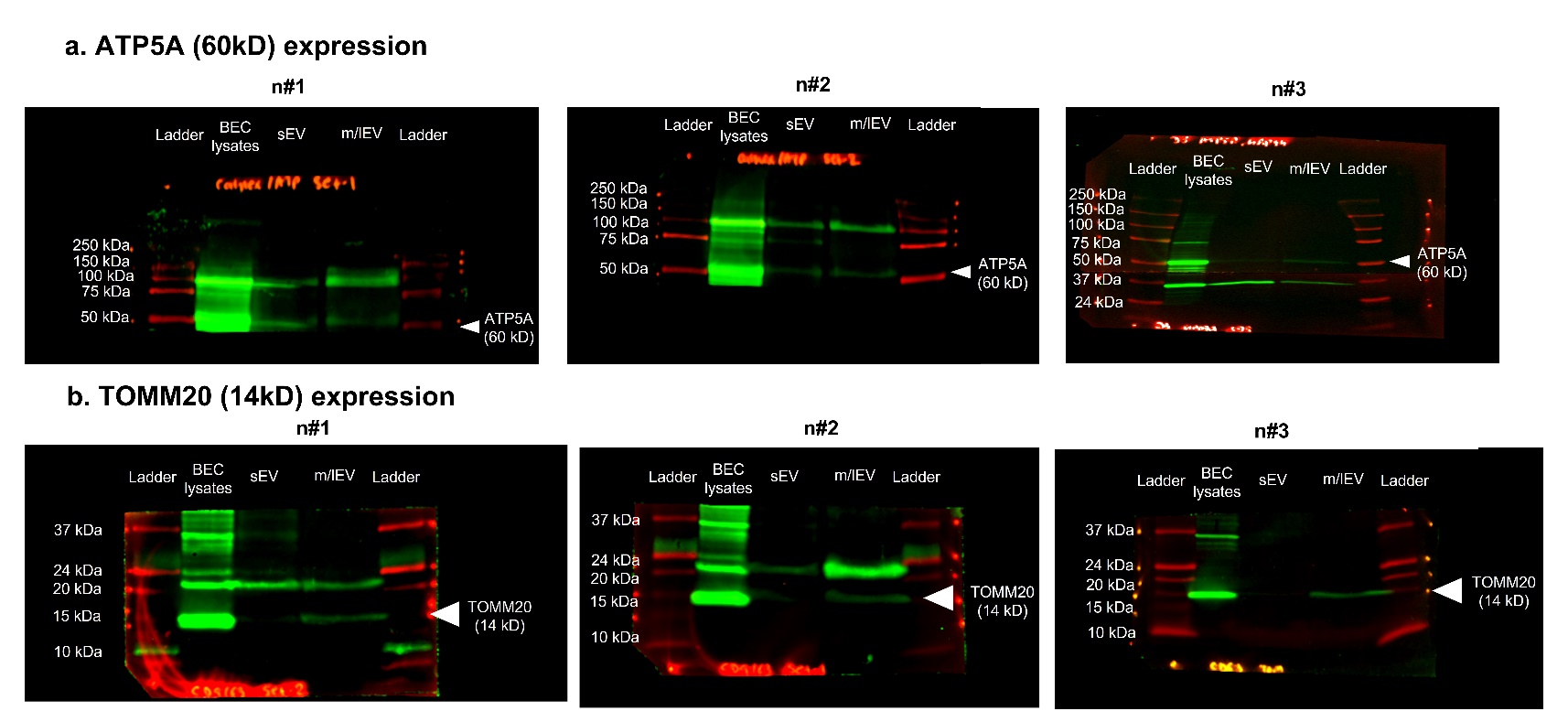

**Fig. S7. Uncropped western blots of ATP5A and TOMM20 expression in sEV and m/lEVs shown in Fig. 2e.**

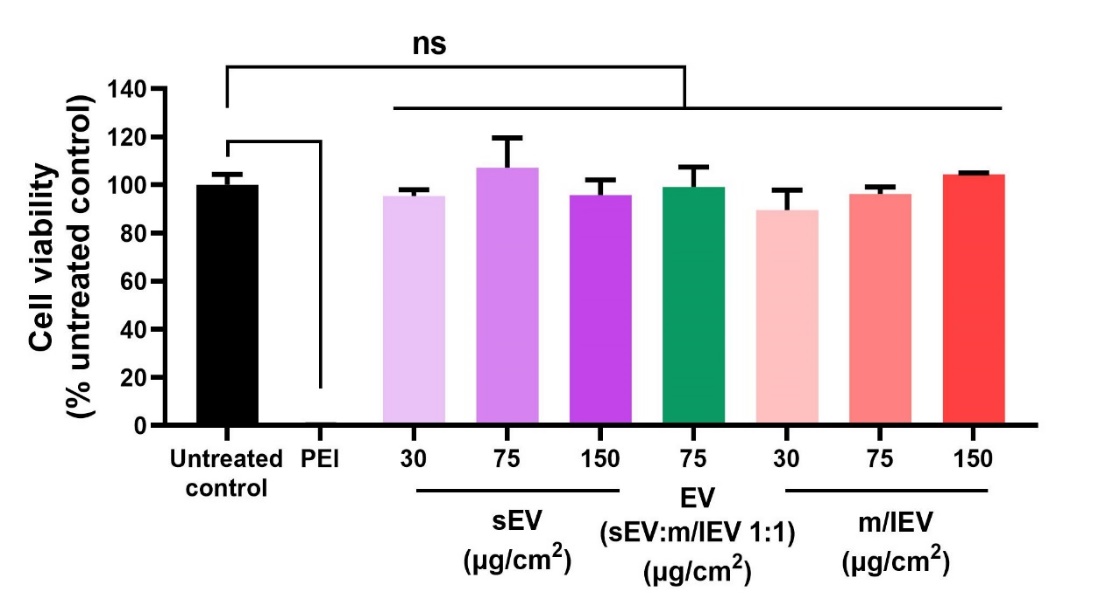

**Fig. S8. Cytocompatibility of MitoTracker Deep red-labelled EVs with human brain endothelial cells.** hCMEC/D3 cells were cultured in the 96-well plates at 16,500 cells/well. The cells were cultured for 4 days in complete growth medium in a 37 °C humidified incubator. Post-confluency, the cells were treated with Mitotracker deep red (MitoT-red) stained sEV and MitoT-red-m/lEVs at 30, 75, and 150 µg EV protein/cm^2^ in complete growth medium for 72 h in a humidified incubator. Seventy-two h post-incubation, the treatment medium was replaced with a 1:1 mixture of growth medium and Cell titer Glo reagent. The relative luminescence units (RLU) of the samples were measured using a SYNERGY HTX multimode plate reader at 1s integration time. The RLU of treatment groups was normalized to the RLU of untreated cells to determine the relative cell viability. Data presents mean±SD (n=6). ****p<0.0001, ns: non-significant.

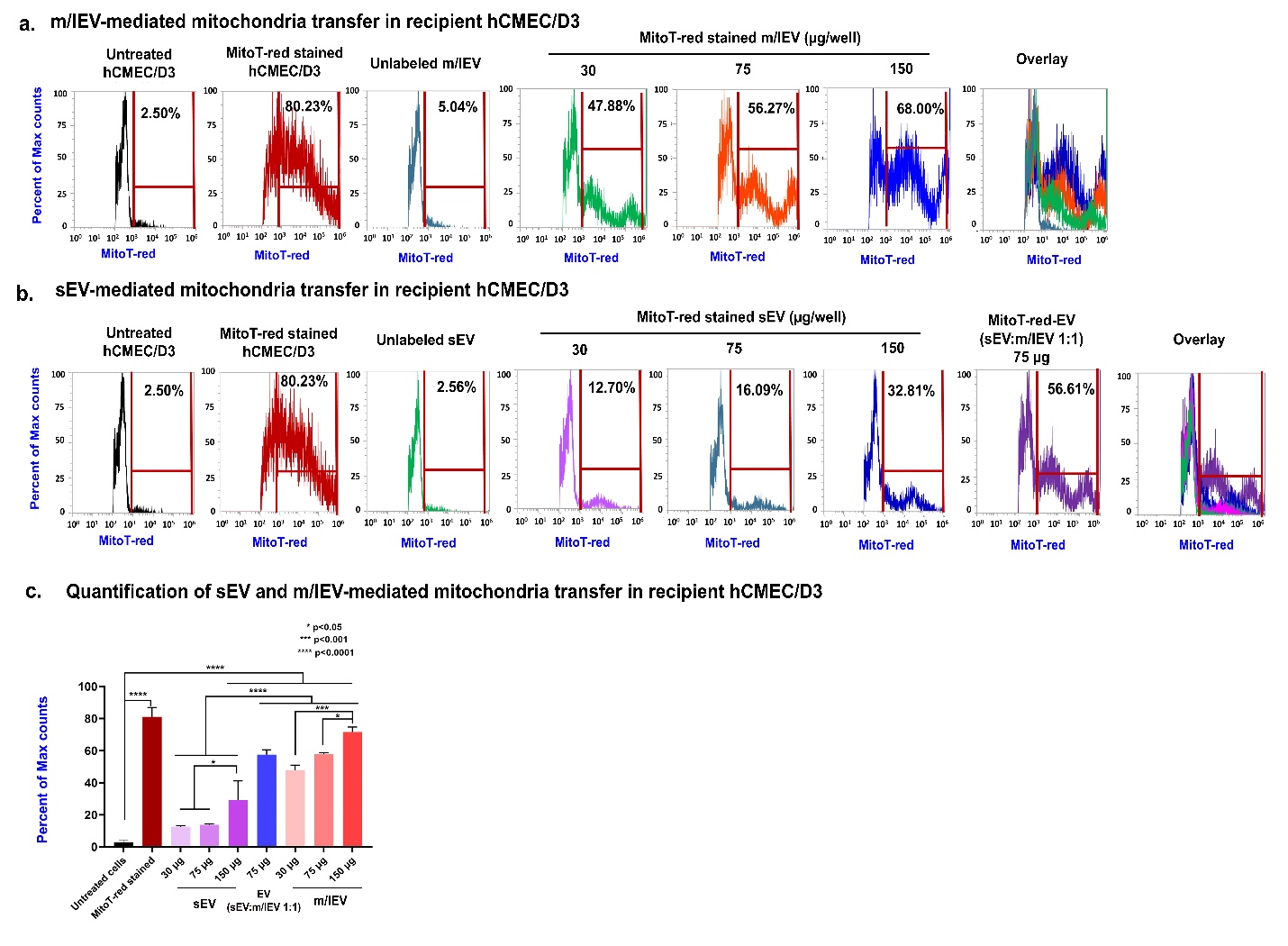

**Fig S9.** **EV mitochondria are transferred into recipient hCMEC/D3 cells at varying doses.** hCMEC/D3 cells were cultured in 48-well plates for 48 h in a humidified incubator. Cells were then incubated with the indicated amounts of MitoTracker red-labeled samples: (MitoT-red)-sEV, MitoT-red-EV (at a 1:1 sEV: m/lEV ratio, collectively referred to as EVs), MitoT-red-MV diluted in complete growth medium for 72 h. The cells were washed, collected, and run through Attune NxT flow cytometer post-incubation. The histograms of MitoT-red events of hCMEC/D3 cells treated at indicated doses of MitoT-red-m/lEV (**a**) and sEV (**b**) for 72 h were detected using a side scatter 674/10-nm filter in an Attune flow cytometer. Unstained EVs were used to gate the histograms for estimating percentage MitoT-red-positive counts. (**c**) Quantification of sEV and m/lEV-mediated mitochondria transfer in recipient hCMEC/D3 cells at 72 h post-exposure.

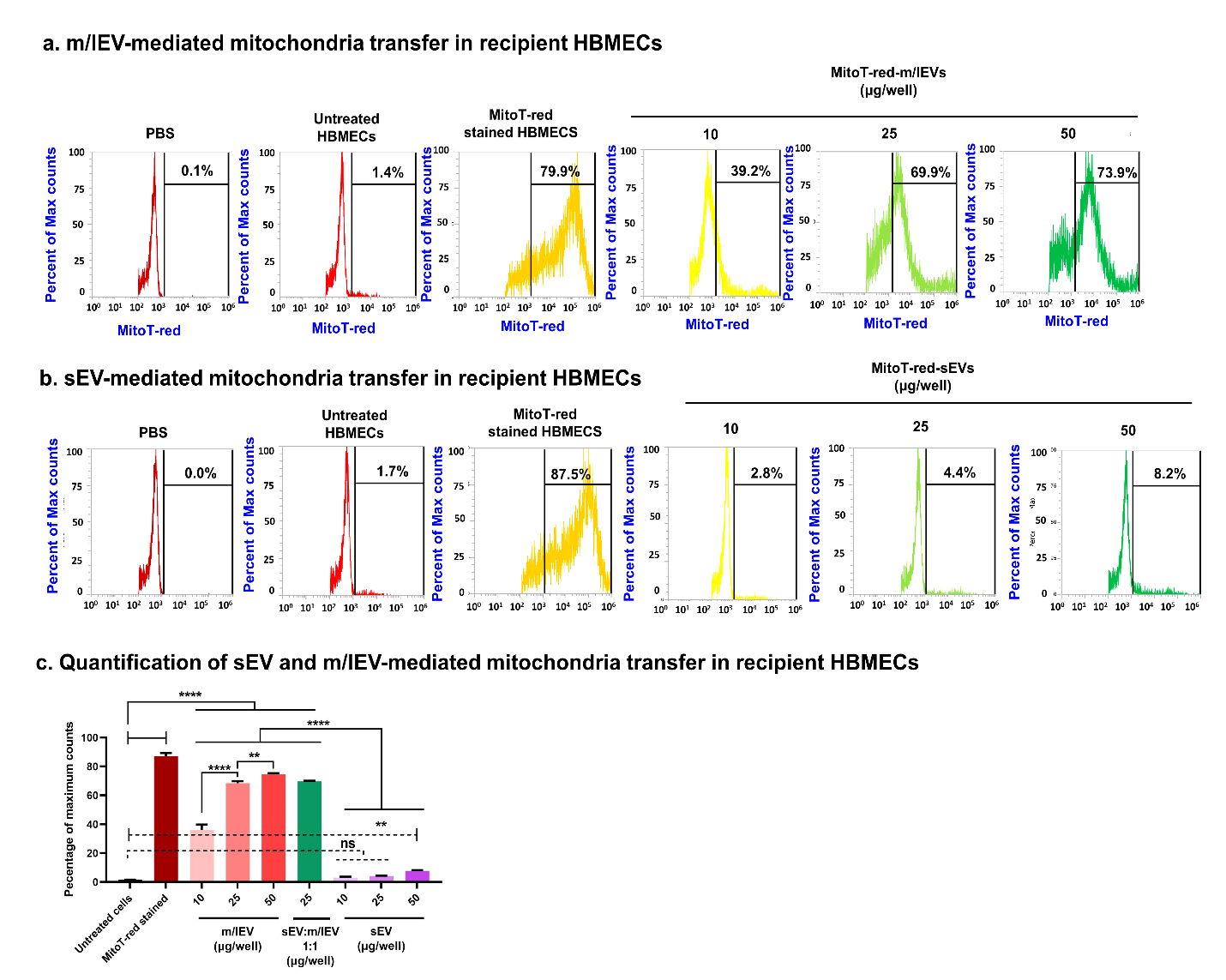

**Fig S10.** **EV mitochondria are transferred into recipient HBMECs at low doses.** HBMECs were cultured in 48-well plates for 48 h in a humidified incubator. Cells were then incubated with the indicated amounts of MitoTracker red-labeled samples: (MitoT-red)-sEV, MitoT-red-EV (at a 1:1 sEV: m/lEV ratio, collectively referred to as EVs), MitoT-red-MV diluted in complete growth medium for 72 h. Post-incubation, the cells were washed, collected, and run through an Attune NxT flow cytometer. The histograms of hCMEC/D3 cells treated at indicated doses of MitoT-red-m/lEV (**a**) and sEV (**b**) for 72 h were collected using a side scatter 674/10-nm filter in an Attune flow cytometer. Untreated HBMECs and unstained EVs were used as controls to gate the background signals in histograms. MitoT-red-stained HBMECs were used as a positive control to gate the histograms for MitoT-red-positive counts. Subsequently, this gate was applied to quantify the percentage of MitoT-red HBMECs treated with MitoT-red-EVs. (**c**) Quantification of sEV and m/lEV-mediated mitochondria transfer in recipient HBMECs at 72 h post-exposure.

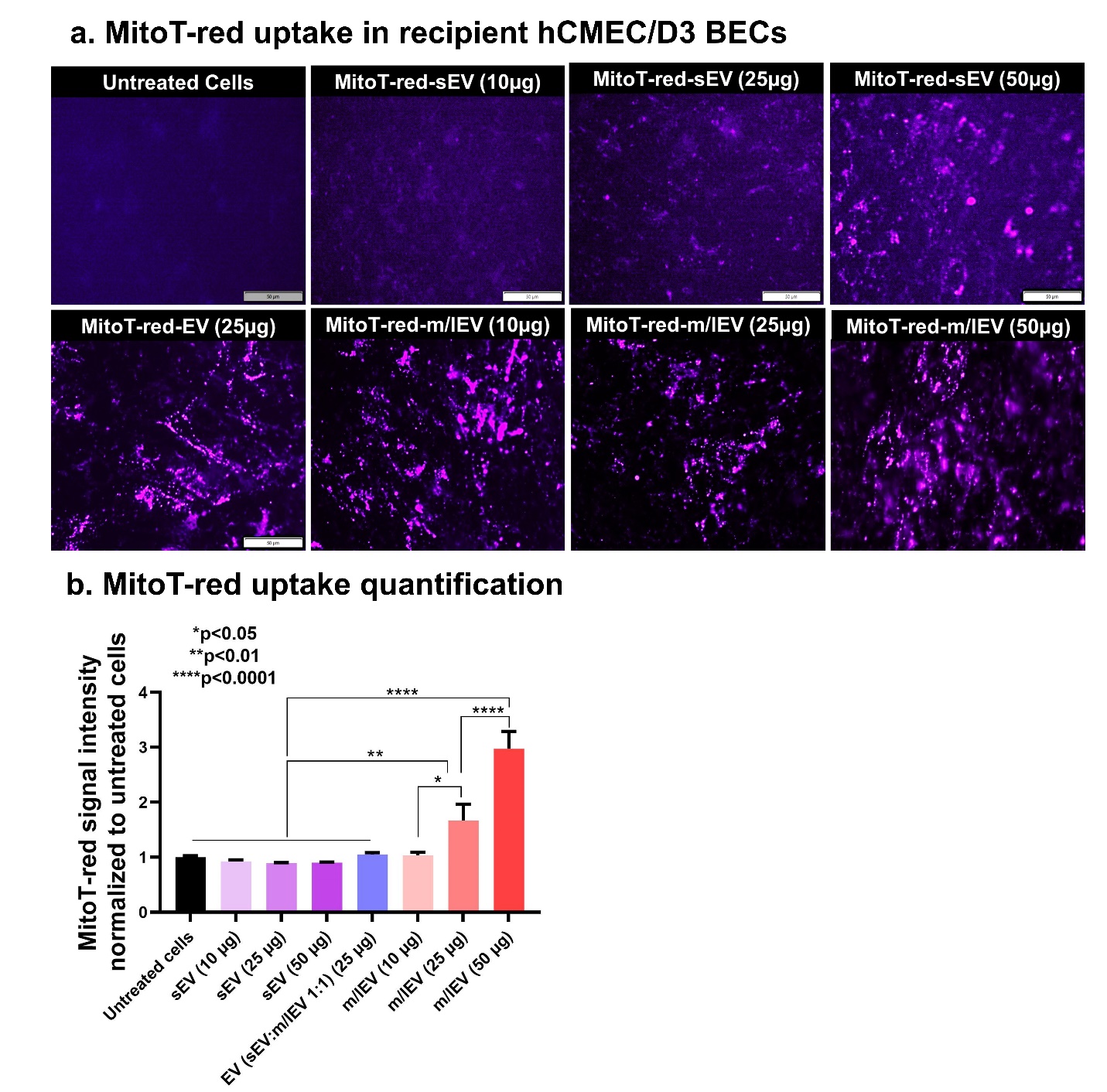

**Fig. S11. Transfer of EV mitochondria into the recipient hCMEC/D3 cells. (a)** hCMEC/D3 were cultured in 96-well plates until 80% confluency in a 37 °C humidified incubator. Post-confluency, cells were incubated with the indicated samples diluted in complete growth medium containing for 72 h. Post-incubation, the treatment mixtures were replaced with phenol-red-free growth medium. Intracellular MitoT-red signals were observed using the Cy5 channel (purple puncta) at 20x magnification using an Olympus IX 73 epifluorescent inverted microscope equipped with CellSens Dimension software. Scale bar: 20 µm. (**b**) The total sum of grayscale intensity was obtained from the acquired images, and the values of treatment groups were normalized to those of untreated cells. Data represent mean±SD (n=3 images per treatment group). *p<0.05, **p<0.01, ***p<0.001, ****p<0.0001

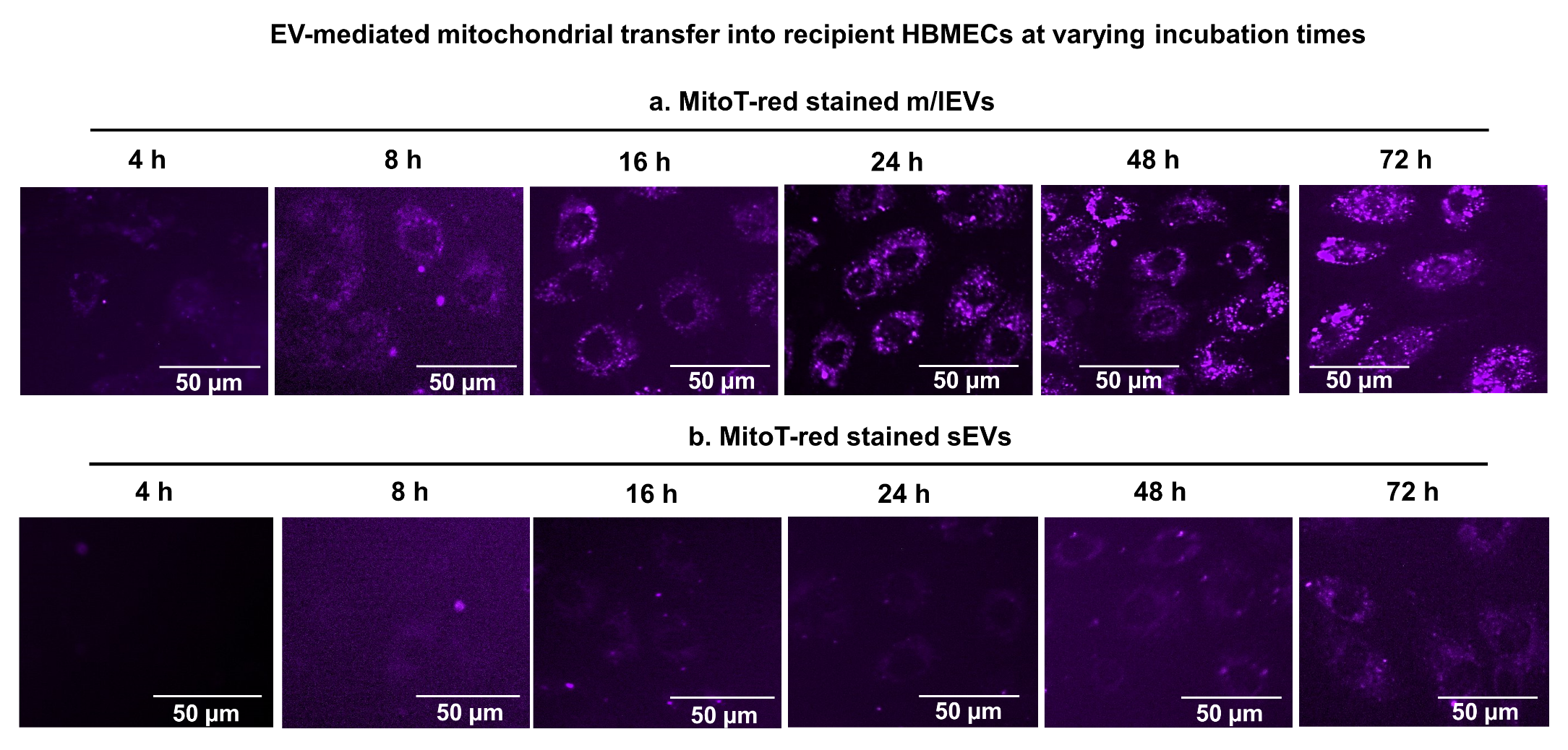

**Fig. S12. Transfer of EV mitochondria into the recipient HBMEC at varying incubation times.** Confluent HBMECs were incubated with 30 µg EV protein of (**a**) MitoT-red-m/lEVs and (**b**) MitoT-red-sEVs diluted in complete growth medium for 4 to 72 h. Post-incubation, the cells were washed and incubated with phenol-red-free growth medium. Intracellular MitoT-red-sEV/m/lEV signals were observed under an Olympus IX 73 epifluorescent inverted microscope using a Cy5 channel (purple puncta). Scale bar: 50 µm.

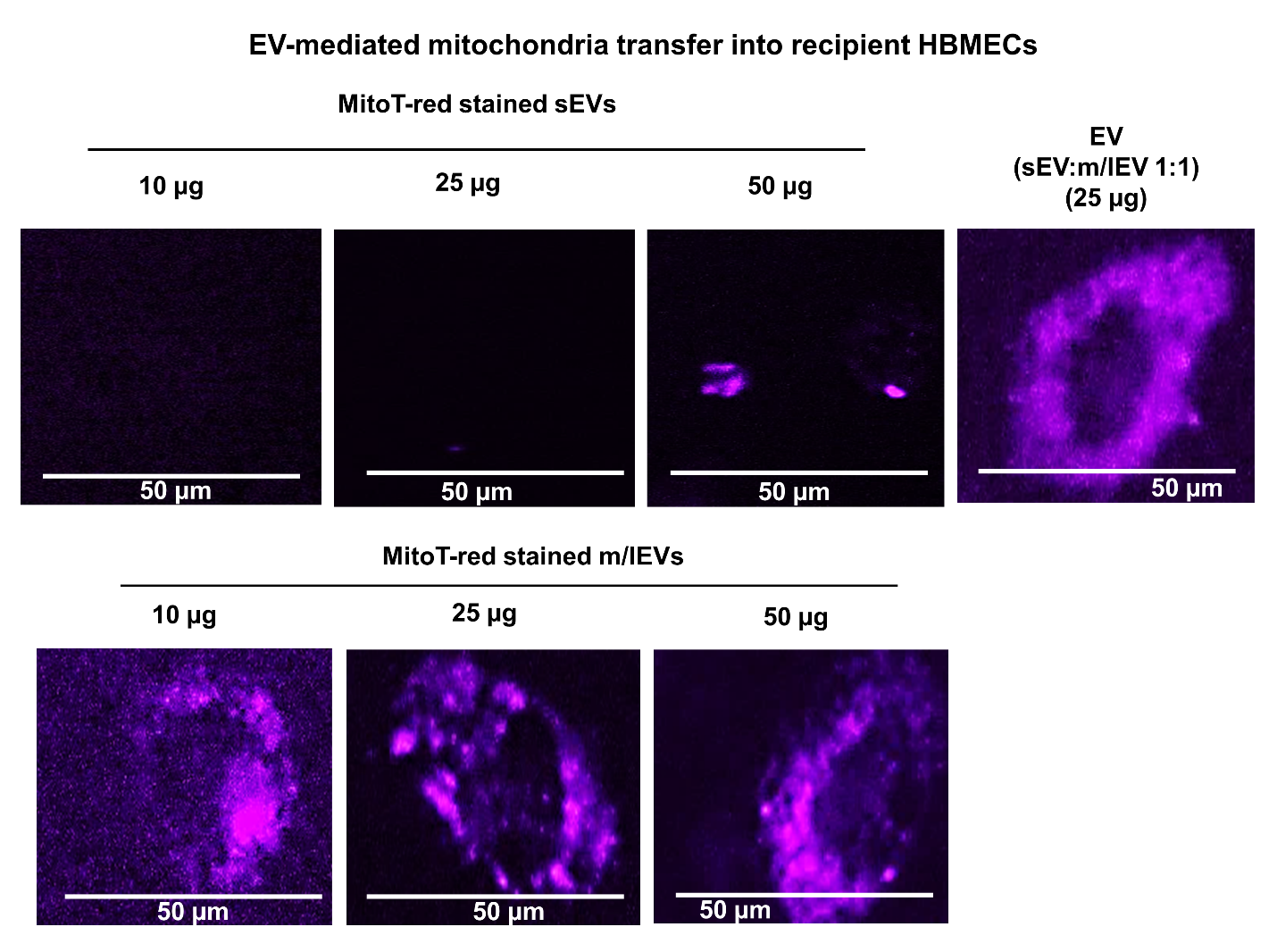

**Fig. S13. Transfer of EV mitochondria into the recipient HBMEC at varying EV doses.** Confluent HBMECs were incubated with 10, 25, and 50 µg EV protein of MitoT-red-sEVs and MitoT-red-m/lEVs diluted in complete growth medium for 24 h. Post-incubation, the cells were washed and incubated with a phenol-red-free growth medium. Intracellular MitoT-red-sEV/m/lEV signals were observed under an Olympus IX 73 epifluorescent inverted microscope using a Cy5 channel (purple puncta). Scale bar: 50 µm. These are magnified images of the main text **Fig. 4**.

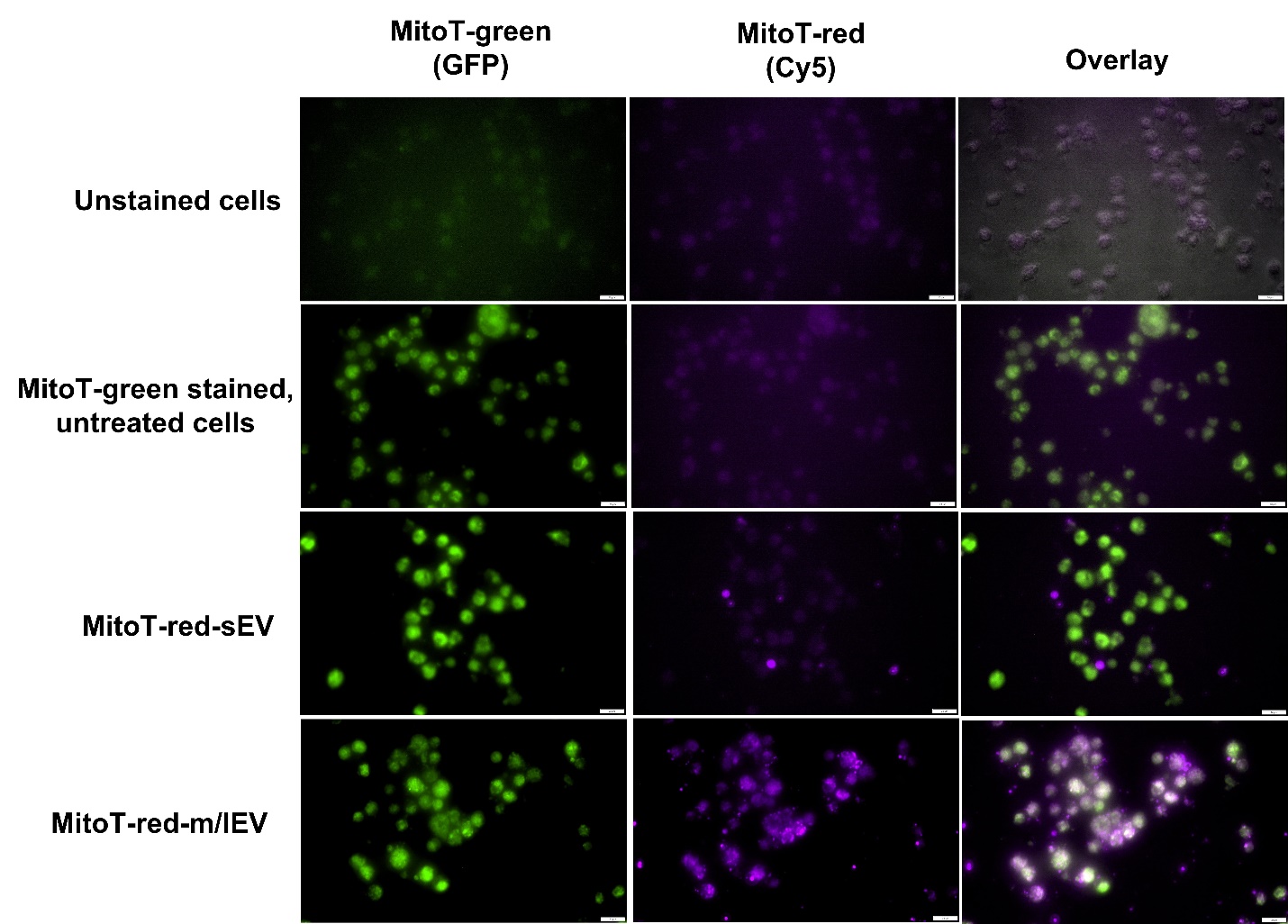

**Fig. S14. Colocalization of EV mitochondrial load with the recipient hCMEC/D3 cell mitochondria.** hCMEC/D3 cells were cultured in 96-well plates until 80% confluency in a 37 °C humidified incubator. Post-confluency, hCMEC/D3 mitochondria were stained with Mitotracker Green for 30 min. Post-staining, the cells were washed and treated with MitoT-red-sEV and MitoT-red-m/lEV at 50 µg/well for 72 h. Untreated cells and cells treated with only MitoTracker Green were used as controls. Post-incubation, the treatment mixture was replaced with phenol-red-free growth medium. Mitotracker Green fluorescence was detected using the GFP channel, whereas the MitoT-red-EV mitochondria were detected using the Cy5 channel in an Olympus IX 73 epifluorescent inverted microscope. The colocalization was observed as yellow signals in the overlay images. Scale bar: 50 µm.

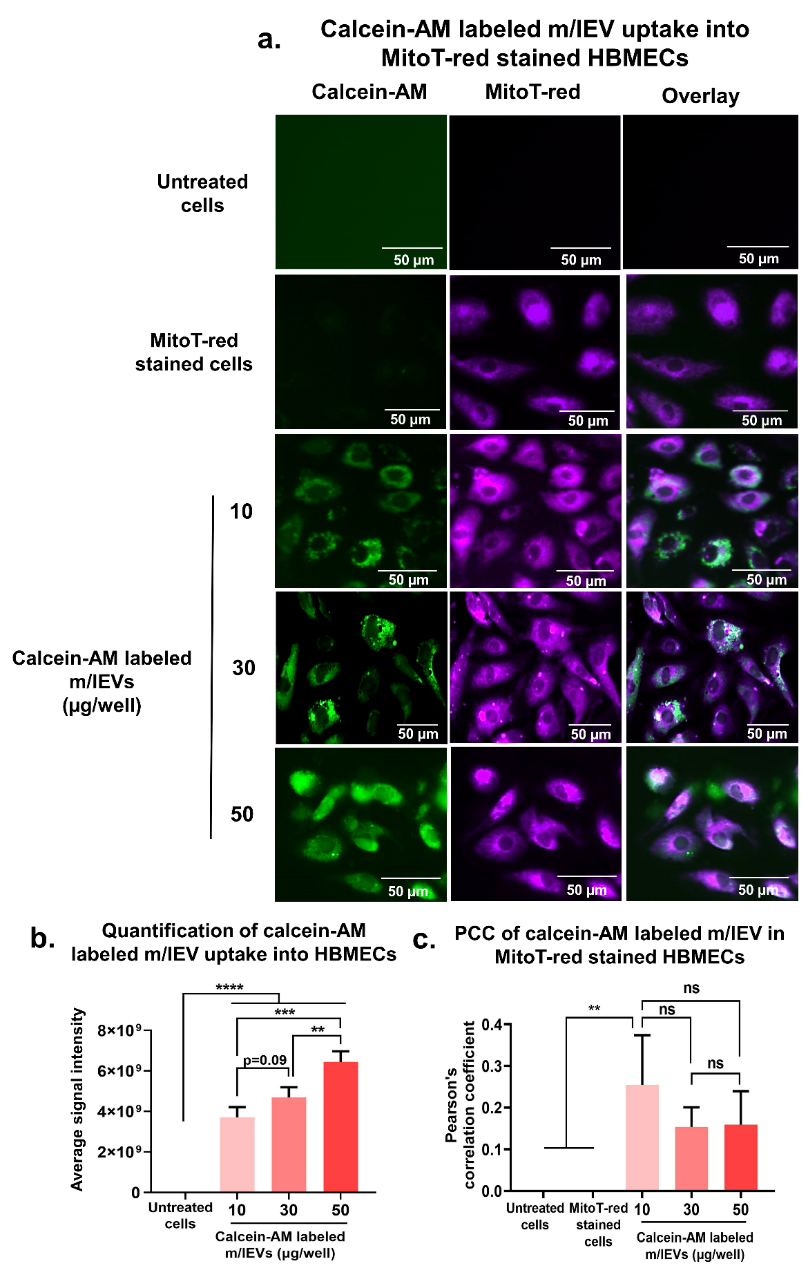

**Fig. S15**. **Calcein-AM labeled m/lEV transfer into MitoT-red-stained HBMECs**. a) Primary HBMECs were treated with calcein-AM labeled m/lEVs at 10- 50 µg EV protein for 24 h. Post-treatment, the cells were washed with 1x PBS and stained with 100 nM MitoT-red in complete culture medium for 30 min. Post-MitoT-red staining, the cells were washed with 1x PBS and treated with phenol-red free culture medium. Untreated cells and cells stained with MitoT-red alone were used as controls. Calcein-AM labeled m/lEV uptake was observed under the GFP channel, whereas MitoT-red-stained HBMECs mitochondria were observed under the Cy5 channel using an Olympus epifluorescence microscope. b) Calcein-AM-m/lEV associated signal intensities in the GFP channel were acquired using cellSens software. c) The Pearson's correlation coefficient (PCC) between the GFP (calcein-AM labeled m/lEVs) and the Cy5 channel (HBMEC mitochondria) was obtained using cellSens software. Data are presented as mean±SD (n=3 images per treatment group). *p<0.05, **p<0.01, ***p<0.001,*p<0.0001, ns: non-significant.

**Discussion for MitoT-red specificity for m/lEVs mitochondria colocalization in the recipient mitochondria**

We performed additional microscopic studies to determine that the observed Pearson's correlation coefficient (PCC) in **Fig. 5b** for colocalization of MitoT-red m/lEVs with recipient MitoT-green stained mitochondria is specific to MitoT-red signals. We used calcein-AM stained m/lEVs as a general EV label (1) and incubated HBMECs stained with Mitotracker deep red (MitoT-red) to evaluate the PCC between calcein-AM and MitoT-red signals. Incubation of EVs with calcein-AM leads to permeation of calcein-AM into EVs. Following that, calcein-AM is hydrolyzed by EV intraluminal esterases and converted into a membrane impermeant green fluorescent calcein (1, 4). HBMECs were incubated with calcein-AM-labeled m/lEVs at 10, 30, and 50 µg EV protein/well for 24 h. The uptake of calcein-AM-labeled m/lEVs in HBMECs was observed under the GFP channel (green fluorescence) using an Olympus epifluorescent microscope. HBMECs mitochondria were stained with MitoT-red and observed under the Cy5 channel (purple fluorescence). PCC for GFP (calcein-AM labeled m/lEVs) and Cy5 channel (HBMEC mitochondria) was obtained using cellSens software.

HBMECs treated with calcein-AM labeled m/lEVs at a 10 µg EV protein showed green fluorescence in HBMECs, suggesting m/lEVs uptake in HBMECs (**Fig. S15a**). The fluorescence intensity increased at doses from 10 to 50 µg EV protein (**Fig. S15a**). Quantitative analysis showed that m/lEV-mediated signal intensity increased significantly (p<0.001) with amounts from 10 to 50 µg EV protein (**Fig. S15b**). On the other hand, MitoT-red signals (purple signals) under the Cy5 channel indicated the presence of mitochondria in HBMECs. We acquired PCC between both channels to determine the correlation between calcein-AM and MitoT-red signals in HBMECs. The PCC of calcein-AM m/lEVs at a 10 µg EV dose was significantly (p<0.01) higher than controls (untreated cells and only MitoT-red treated cells, **Fig. S15c**). However, there was no statistical difference in the PCC among different m/lEVs doses (**Fig. S15c**). Collectively, the dose-dependent increase in calcein-AM labeled m/lEV uptake, but not the correlation between calcein-AM labeled m/lEVs and MitoT-red-stained HBMEC mitochondria suggests that calcein-AM labeled m/lEVs were not colocalized with HBMEC mitochondria. However, MitoT-red-stained m/lEVs showed a dose-dependent significant (p<0.0001) increase in PCC with MitoT-green stained HBMEC mitochondria (**Fig. S16**). Therefore, the observed dose-dependent significant increase in the PCC of MitoT-red-stained EVs and MitoT-green-stained HBMEC mitochondria is likely specific to MitoT-red signals (**Fig. S16**).

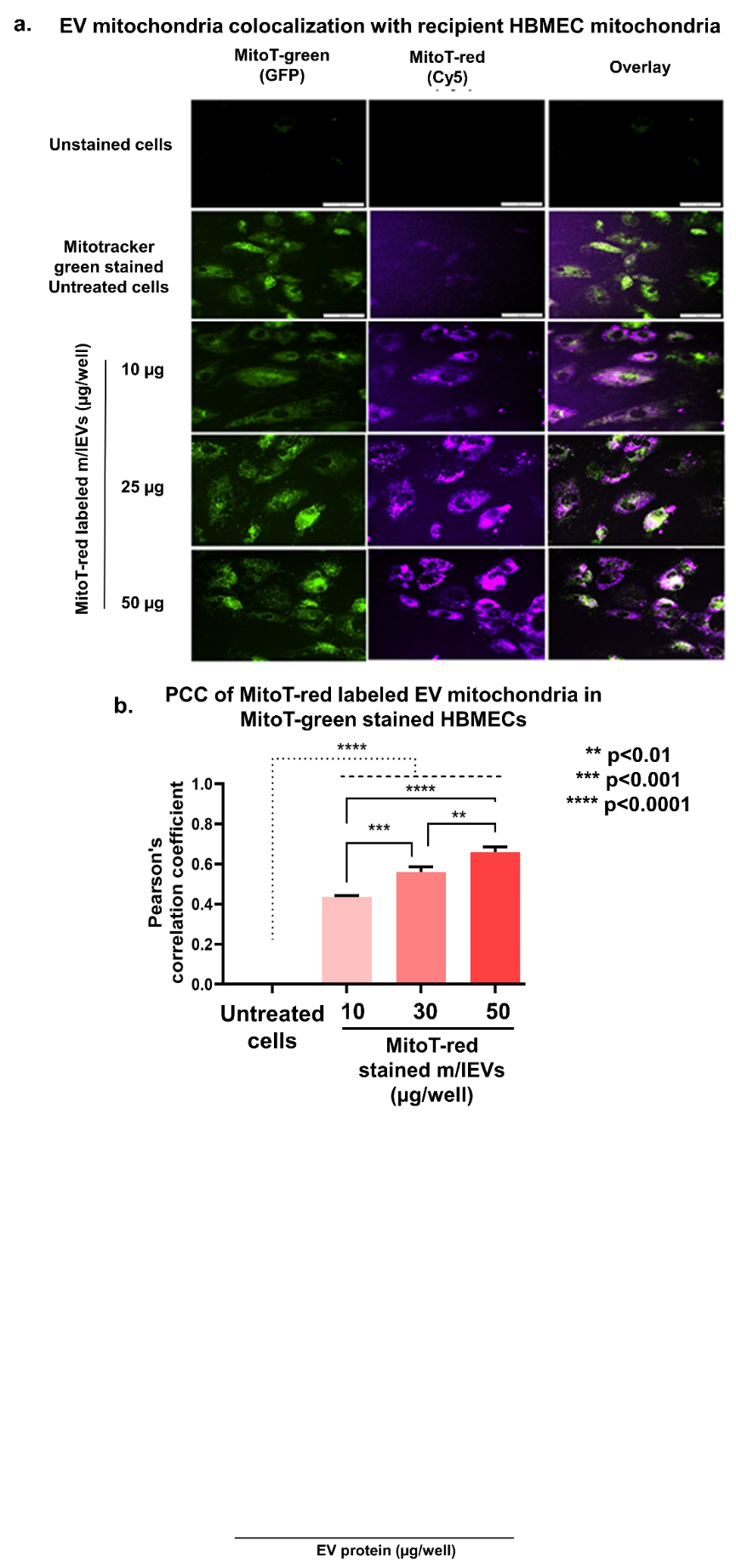

**Fig. S16.** **Colocalization of EV mitochondria with the recipient HBMEC mitochondria. (a)** HBMECs were cultured in 96-well plates until 80% confluency in a humidified incubator. HBMECs were stained with Mitotracker Green for 30 min. Post-staining, the cells were washed and treated with the indicated doses of MitoT-red-m/lEV for 24 h. Untreated cells and cells stained with MitoTracker Green alone were used as controls. Post-incubation, the treatment mixture was replaced with phenol-red-free growth medium. The Mitotracker green fluorescence in recipient HBMEC was detected under the GFP channel, whereas the purple fluorescence associated with EV mitochondria was captured under the Cy5 channel in an Olympus IX 73 epifluorescent inverted microscope. Colocalization of the mitochondrial signals was confirmed by the presence of yellow signals in the overlay images. Scale bar: 50 µm. (**b**) Pearson's correlation coefficient was obtained from the overlay images of Cy5 and GFP channels at constant signal intensities for both channels using cellSens software. Data are presented as mean±SD (n=3 images per treatment group). *p<0.05, **p<0.01, ***p<0.001, ****p<0.0001

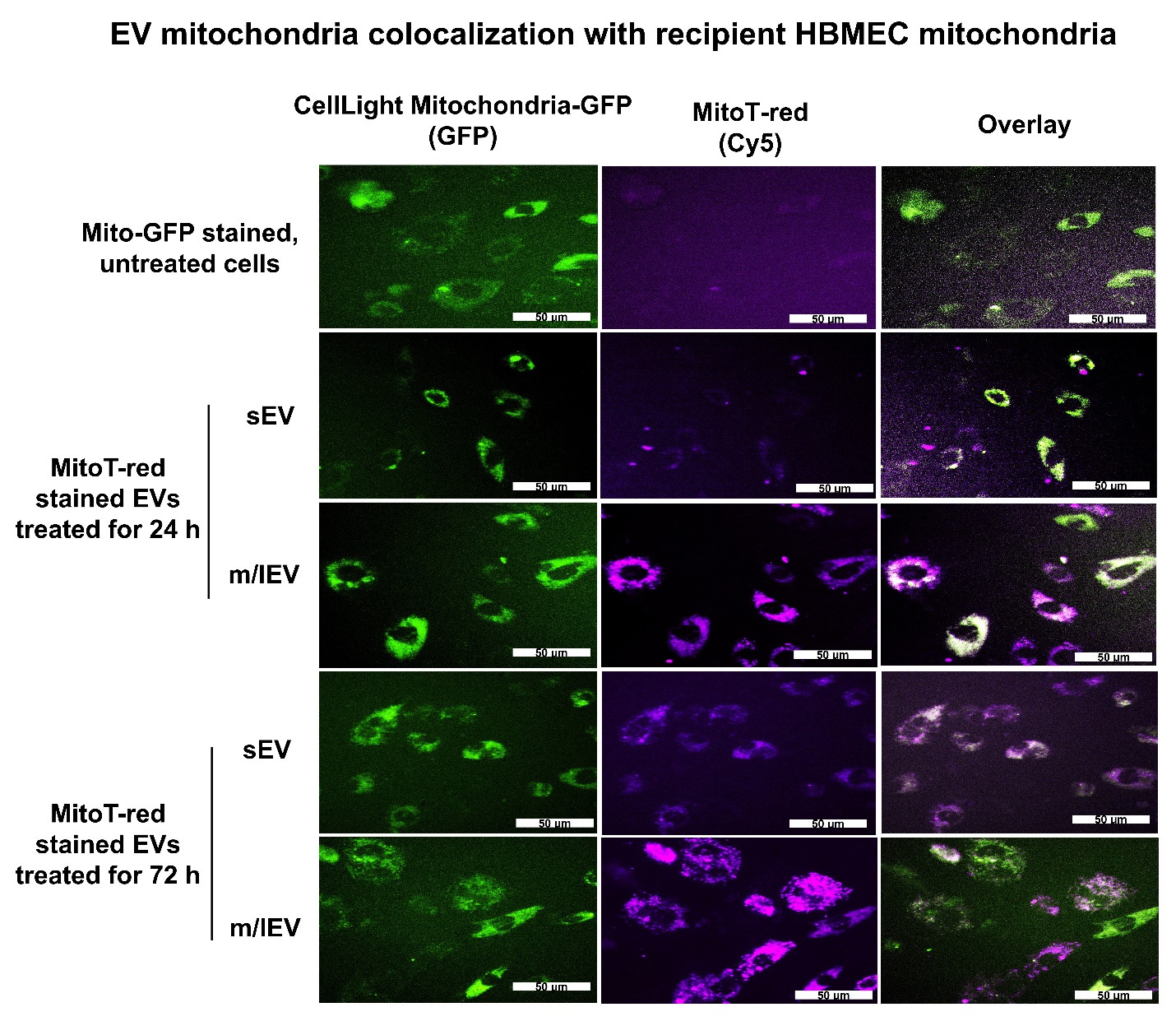

**Fig. S17. Colocalization of EV mitochondria with the recipient HBMEC mitochondria using CellLight Mitochondria-GFP BacMam technique.** Confluent HBMECs were transduced with CellLight Mitochondria-GFP at 2µL/10,000 cells for 16-18 h. The transduction mixture was then removed, cells were washed, and treated with complete growth medium containing MitoT-red-sEV and m/lEV at 50 µg/well for 24 and 72 h. Untreated cells and cells treated only with CellLight Mitochondria-GFP were used as controls. The green fluorescence associated with CellLight Mitochondria-GFP in recipient HBMECs was acquired using the GFP channel, whereas the purple fluorescence associated with polarized mitochondria from MitoT-red sEV and m/lEV was captured using Cy5 channel using an Olympus IX 73 epifluorescent inverted microscope. Scale bar: 50 µm.

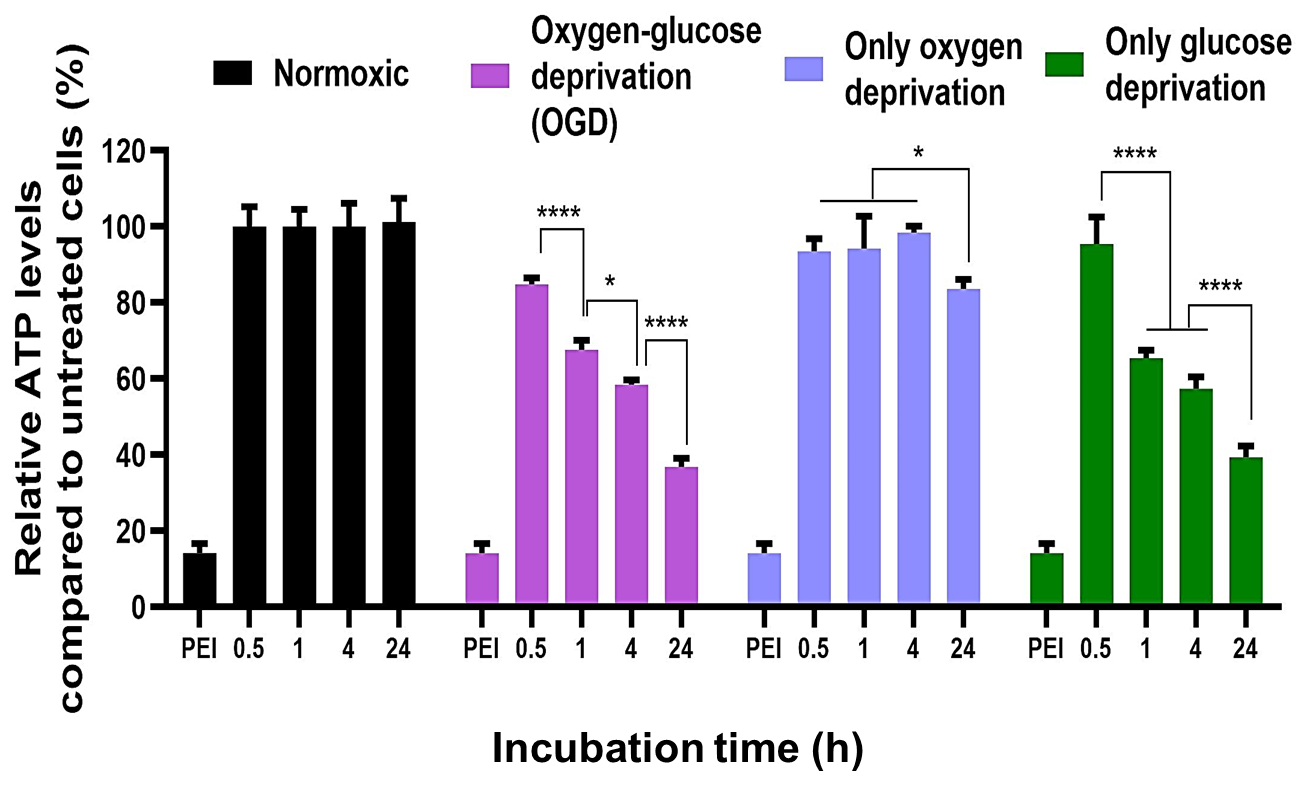

**Fig. S18. *In vitro* oxygen glucose-deprivation model.** HBMECs were cultured in the 96-well plates until 80% confluency in a complete growth medium in a 37 °C humidified incubator. Confluent normoxic HBMEC monolayers were treated with OGD medium and incubated in a hypoxic Billups-Rothenberg chamber at 37°C for 24 h. The following groups were also included: Normoxic HBMECs cultured in complete growth medium in a humidified incubator, oxygen-deprived cells cultured in complete growth medium in a hypoxic chamber, and glucose-deprived cells cultured in an OGD medium in a normoxic chamber. The cell viability of normoxic HBMECs was used as a control to calculate the cell viability in all treatment conditions. Data represent mean±SD (n=3). * p<0.05, ** p<0.01, *** p<0.001, **** p<0.0001.

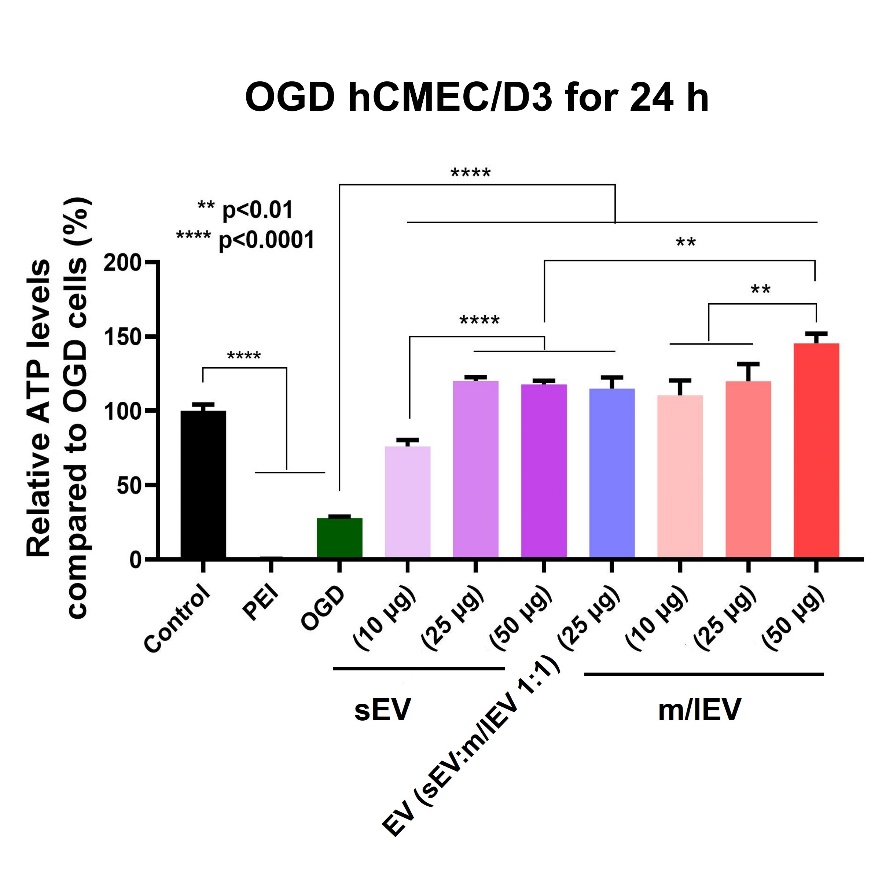

**Fig. S19. EV-mediated increase in hCMEC/D3 ATP levels during hypoxic conditions.** hCMEC/D3 cells were cultured in the 96-well plates until 80% confluency in complete growth medium in a 37 °C humidified incubator. Normoxic confluent monolayers were treated with the indicated doses of sEV and m/lEV in OGD medium, and the cell viability was measured 24 h post-treatment. Untreated cells were used as a control. Data represent mean±SD (n=3). * p<0.05, ** p<0.01, *** p<0.001, **** p<0.0001.

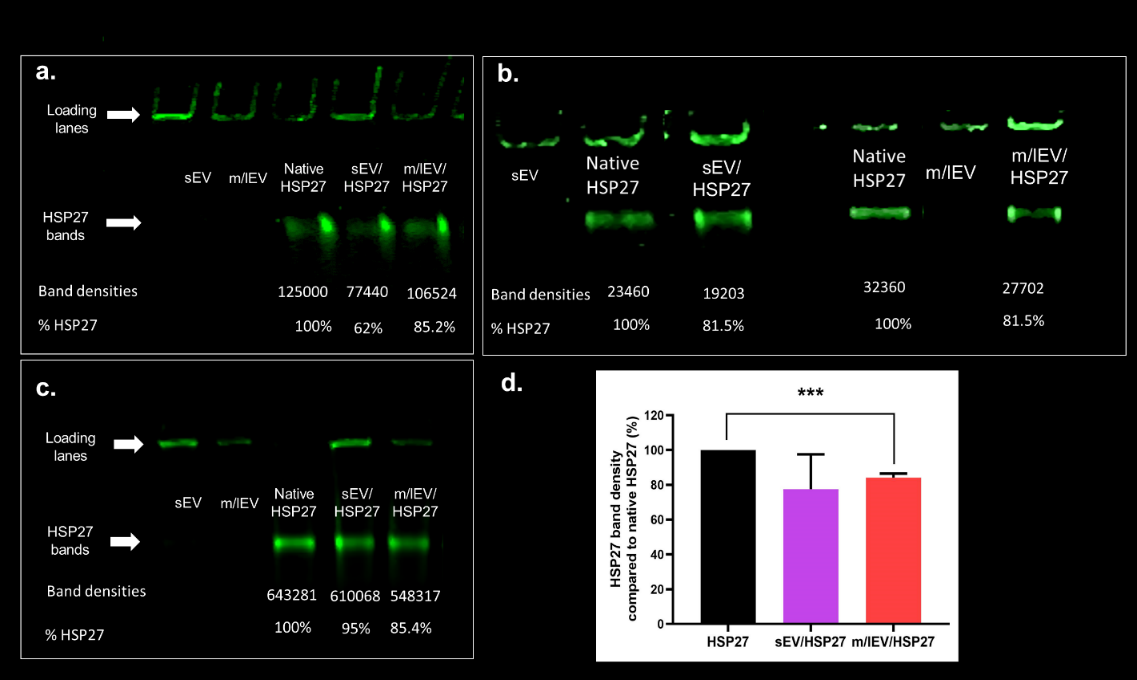

**Fig. S20. Formation of EV/HSP27 mixtures.** (**a-c)** Native HSP27 and mixtures of sEV/HSP27 and m/lEV/HSP27 at 10:1 weight/weight (w/w) ratios were loaded in an SDS-free 4-10% polyacrylamide gel at one µg HSP27 per lane. Free sEV and m/lEV equivalent to the amounts in 10:1 w/w mixtures were used as controls. The indicated samples were loaded in the gel at one µg HSP27/lane. Each gel was run at 100 V for two h and stained using Biosafe Coomassie G250. The gel was then scanned at 800 nm using an Odyssey imager at intensity setting 5. (**d**) Densitometry analysis was performed by measuring band densities of HSP27 in the different experimental groups in comparison to the band density of native HSP27 in the respective gel using Image Studio 5.0 software ***p<0.001

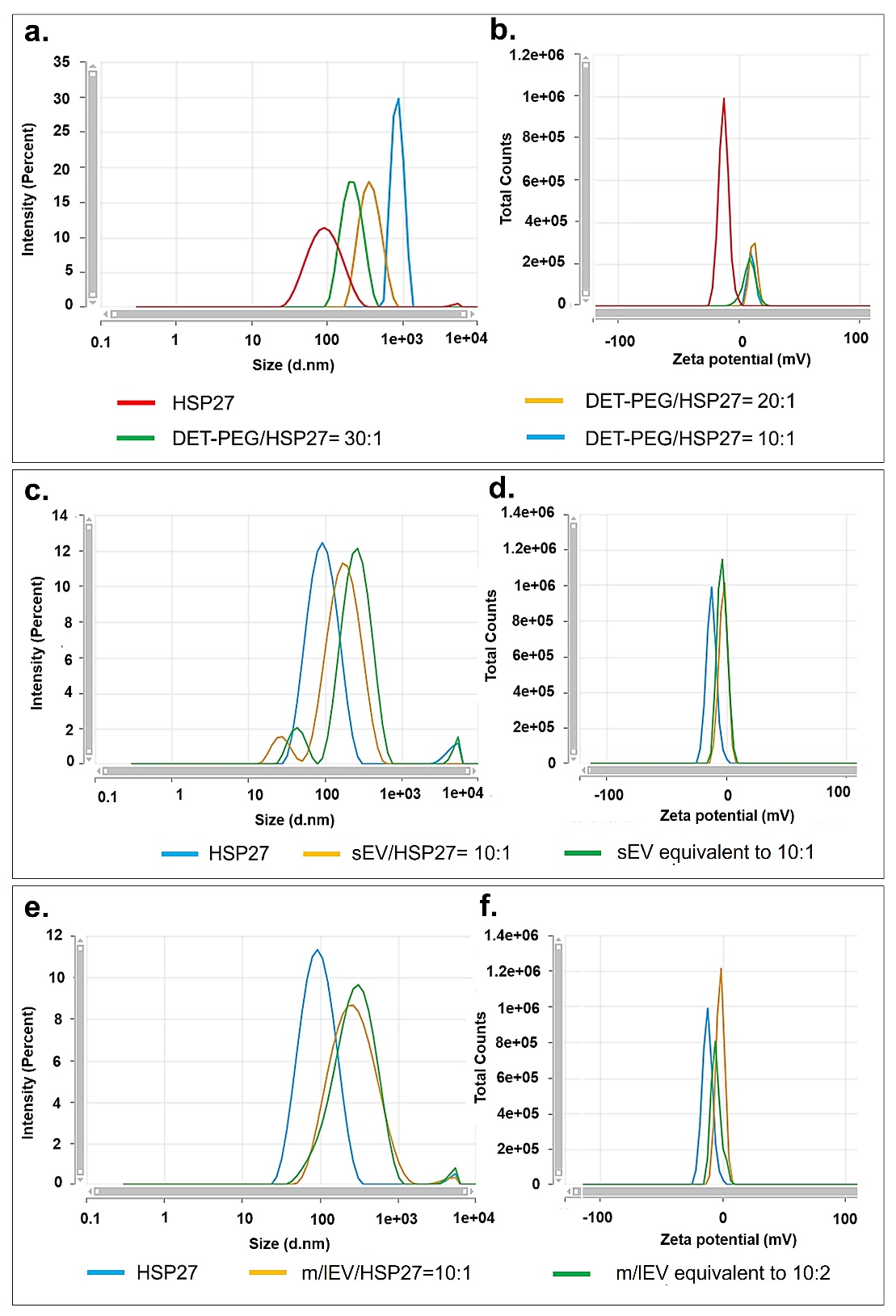

**Fig. S21. Intensity distribution and total photon count vs. zeta potential plots of HSP27 mixtures.** Native HSP27, and PEG-DET/HSP27 at 10:1, 20:1, and 30:1 w/w ratios (**a,b**) and sEV/HSP27 (**c**), m/lEV/HSP27 (**d**) at 10:1 w/w ratios were diluted in 10 mM HEPES buffer pH 7.4 for particle size measurements, and the sample was further diluted in 800 µL of 10 mM HEPES buffer pH 7.4 for zeta potential measurements using a Malvern Zetasizer Pro.

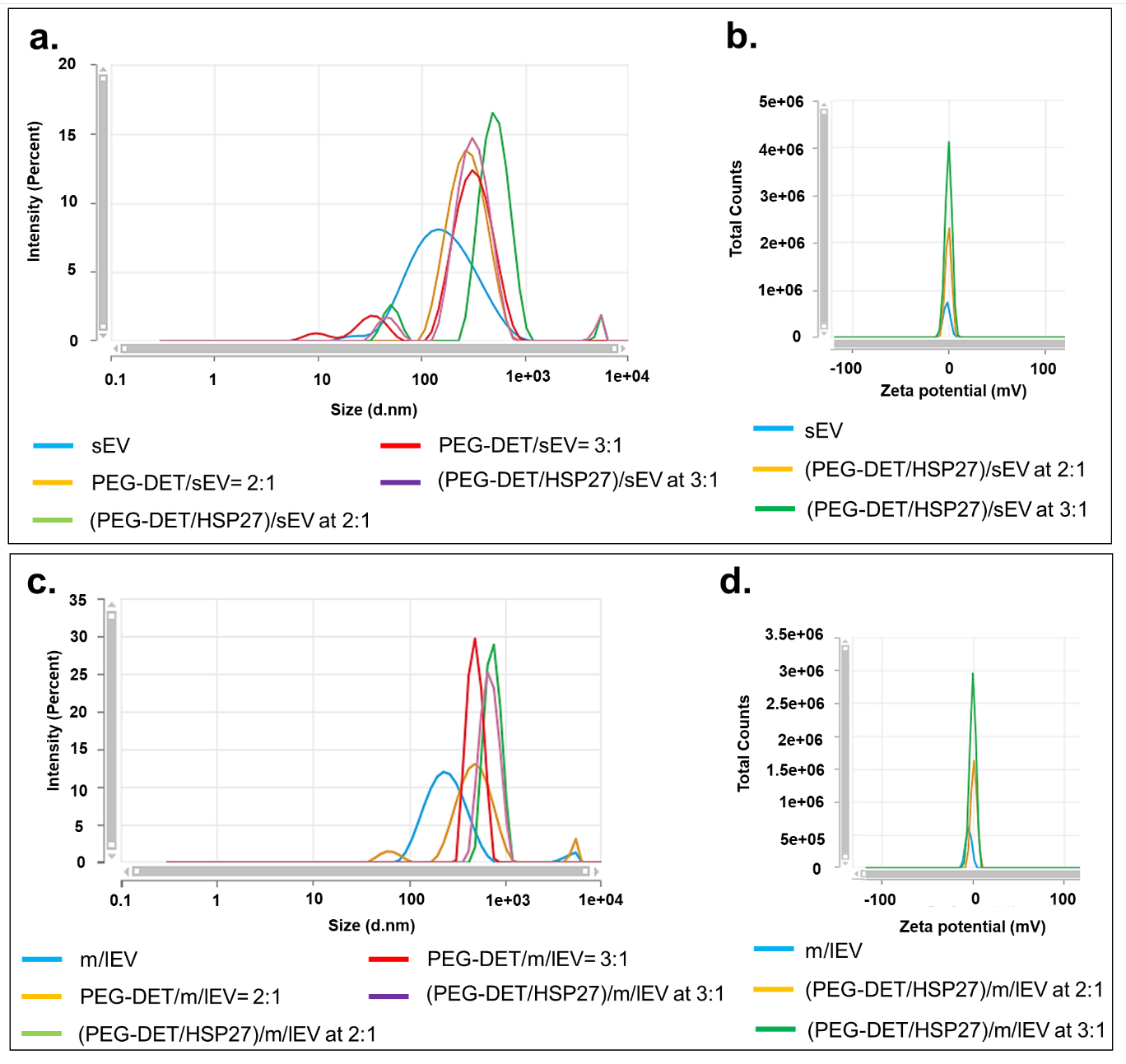

**Fig. S22. Intensity distribution and total photon count vs. zeta potential plots of (PEG-DET/HSP27)/EV ternary mixtures.** PEG-DET/HSP27 mixtures were prepared at 20:1 and 30:1 w/w ratios followed by mixing with 10 µg of sEV (**a,b**) and m/lEV (**c,d**). PEG-DET/HSP27, EV/HSP27, and (PEG-DET/HSP27)/EV binary and ternary mixtures were diluted in 10 mM HEPES buffer pH 7.4 for particle size measurements. The samples were further diluted into 800 µL of 10 mM HEPES buffer pH 7.4 for zeta potential measurements using a Malvern Zetasizer Pro.

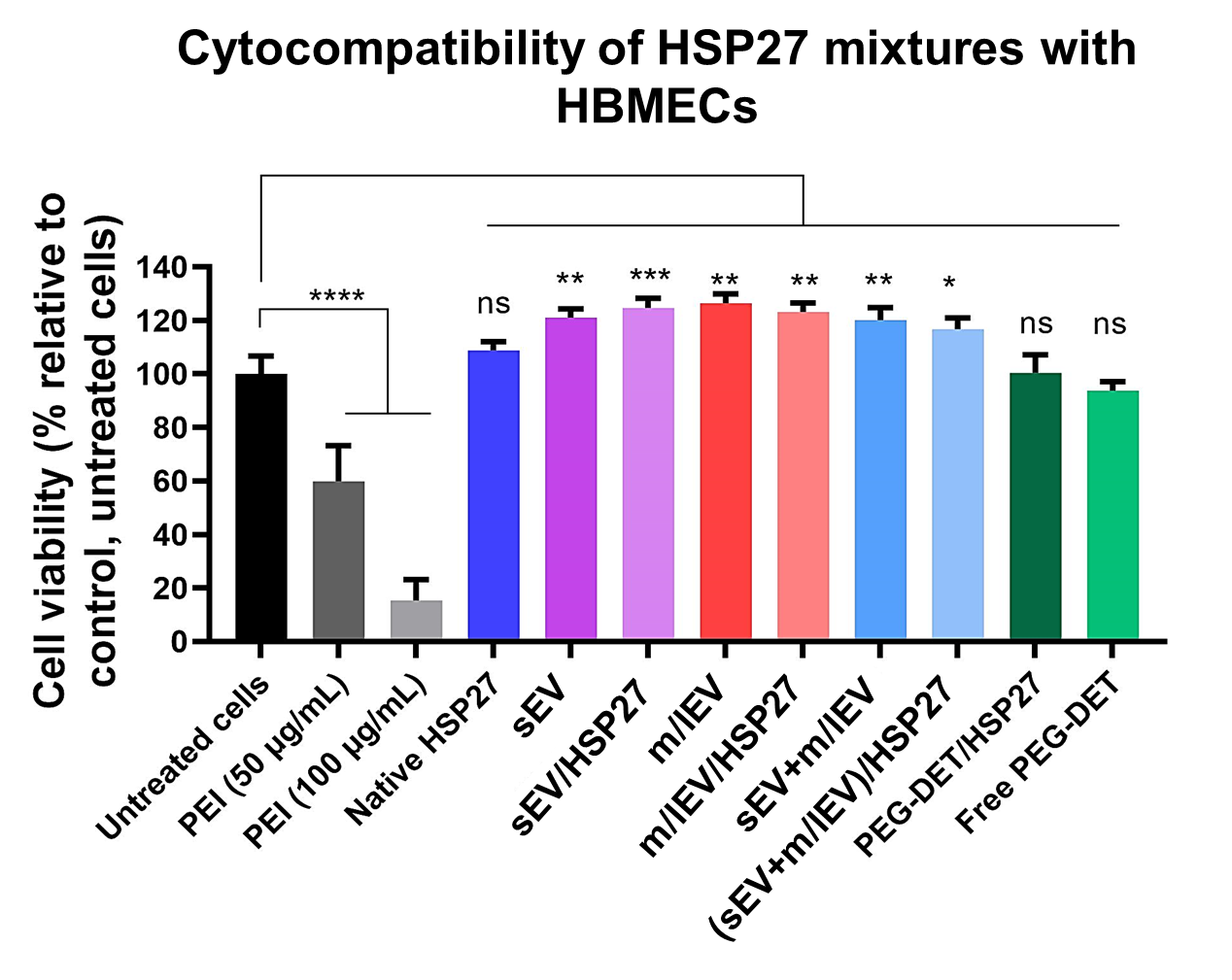

**Fig. S23. Cytocompatibility of HSP27 mixtures with HBMECs.** Normoxic confluent monolayers were treated with the indicated samples at a dose of 2 µg of HSP27 per well. Cells were treated for 72 h before measuring cell viability, and values were normalized to untreated cells. Data represent mean±SD (n=3). * p<0.05, ** p<0.01, *** p<0.001, **** p<0.0001.

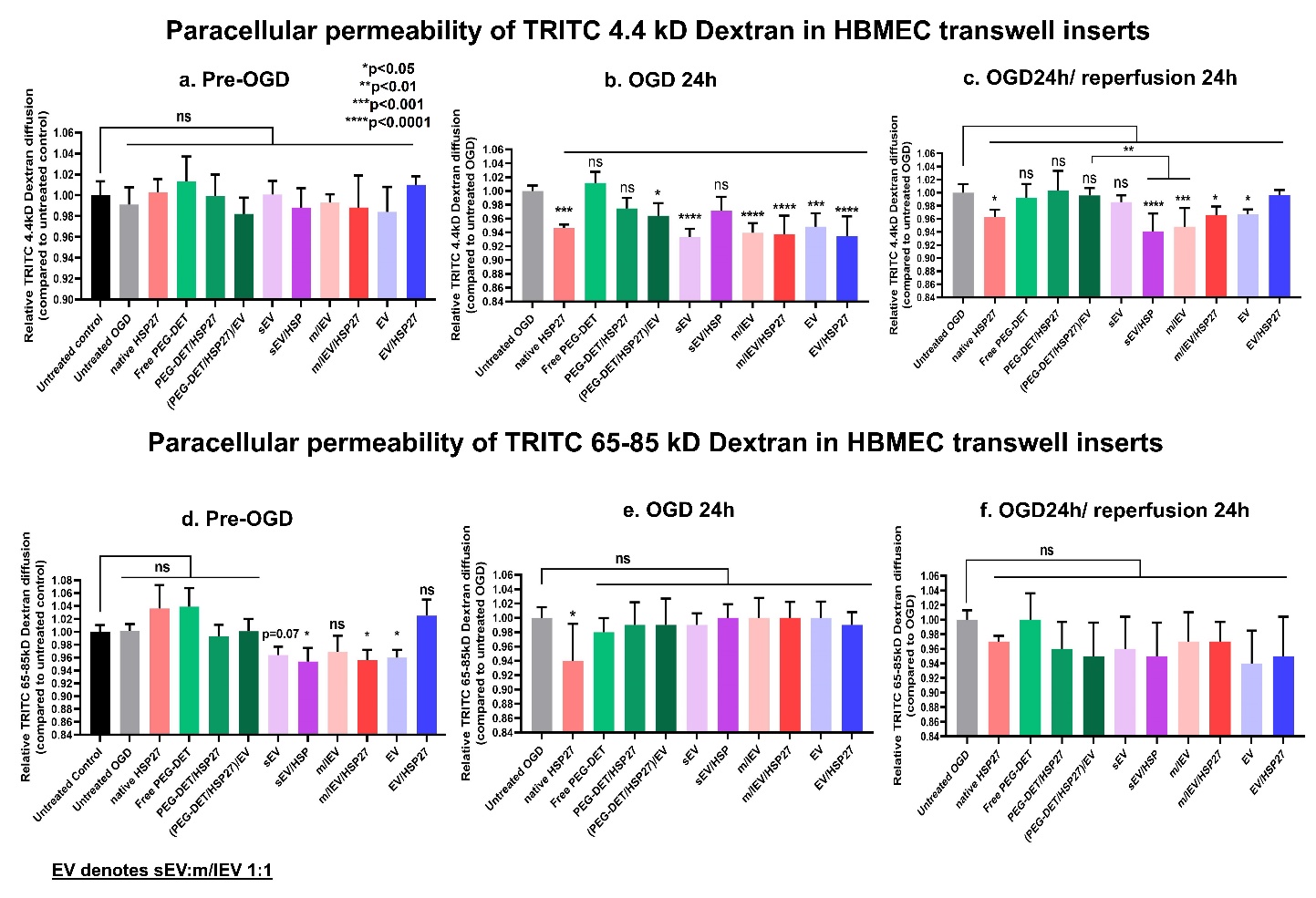

**Fig. S24: Paracellular permeability of 4.4 kD and 65-85 kD TRITC-Dextran pre-OGD, under 24 h of OGD exposure, and OGD 24 h/24 h of reperfusion conditions in pretreated HBMEC transwell culture inserts.** HBMECs seeded in 24-well transwell inserts were cultured in a 37°C humidified incubator for a week. The complete growth medium was replaced with 300 µL of growth medium containing the indicated treatment groups for 72 h. Post-incubation, the medium was replaced with a complete growth medium containing 1µM 4.4 kD (**a**) or 65-85 kD TRITC-Dextran (**d**) for 1 h (Pre-OGD). Post-treatment, the diffusion of 4.4 kD dextran (**b**) and 65-85 kD dextran (**e**) were measured in OGD conditions. Post-OGD treatment, HBMECs were washed with PBS and incubated with 300 µL of complete growth medium containing 1µM 4.4 kD (**c**) or 65-85 kD (**f**) TRITC-Dextran and then incubated in a humidified incubator for 24 h. The diffusion of 4.4 kD dextran and 65-85 kD dextran were measured in OGD/reperfusion conditions at 24 h. The concentrations of 4.4 and 65-85 kD TRITC-Dextran were measured using a Synergy HTX multimode plate reader at 485/20 excitation and 580/50 nm emission settings. The relative diffusion of TRITC Dextran at each time point was determined by calculating the ratio of [TRITC-Dextran] in the abluminal compartment of treatment groups to that of untreated OGD control. Data represent mean±SD (n=4). * p<0.05, ** p<0.01, *** p<0.001, **** p<0.0001, ns: non-significant.

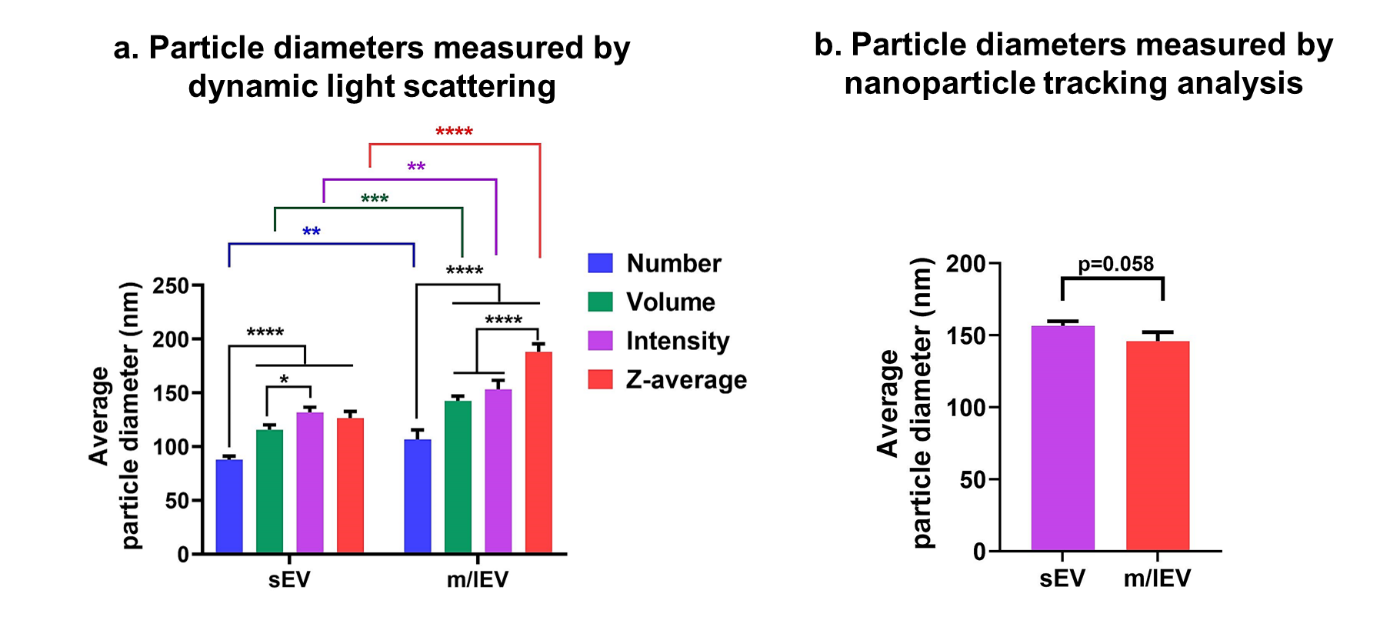

**Fig. S25. Average particle diameter of sEV and m/lEV measured using DLS and NTA.** (**a**) the average particle diameter of sEV and m/lEV at 0.1 mg/mL in PBS was measured based on a number, volume, intensity, and z-average using DLS (Malvern Zetasizer Pro). (**b**) sEV and m/lEV at 0.5 mg/mL in PBS were diluted 100x for NTA.

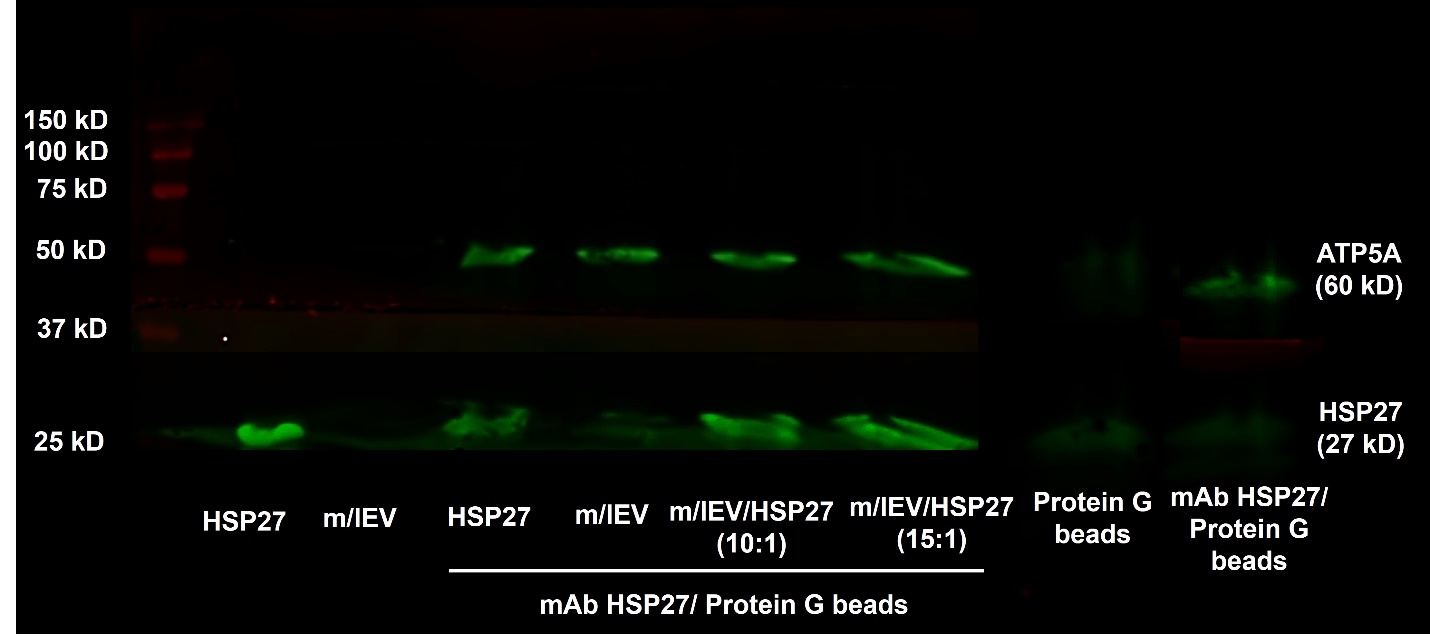

**Fig. S26. Pull-down assay of m/lEV/HSP27 mixtures via protein G magnetic beads using immunoprecipitation method.** m/lEV/HSP27 mixtures at 10:1 and 15:1 weight: weight ratios were incubated with HSP27 antibody for 2 h at room temperature, then incubated with protein G magnetic beads for 1 h at room temperature. The magnetic beads were collected and incubated with Laemmli SDS sample buffer for 10 min at 95°C. Next, the beads were separated, and the supernatant was run through 4-10% SDS-PAGE at 120 V for 90 min. Native HSP27 and m/lEVs not incubated with HSP27 antibodies and protein G beads were used as controls. The proteins were transferred to a 0.45 µm nitrocellulose membrane. The membrane was incubated with a blocking solution for 1 h at room temperature. The membrane was incubated with mouse ATP5A and HSP27 primary antibodies overnight at 4°C. Following that, the membrane was incubated with secondary antibodies for 1 h at room temperature. The blot was scanned under Odyssey imager at 800 nm channel and intensity settings 5.

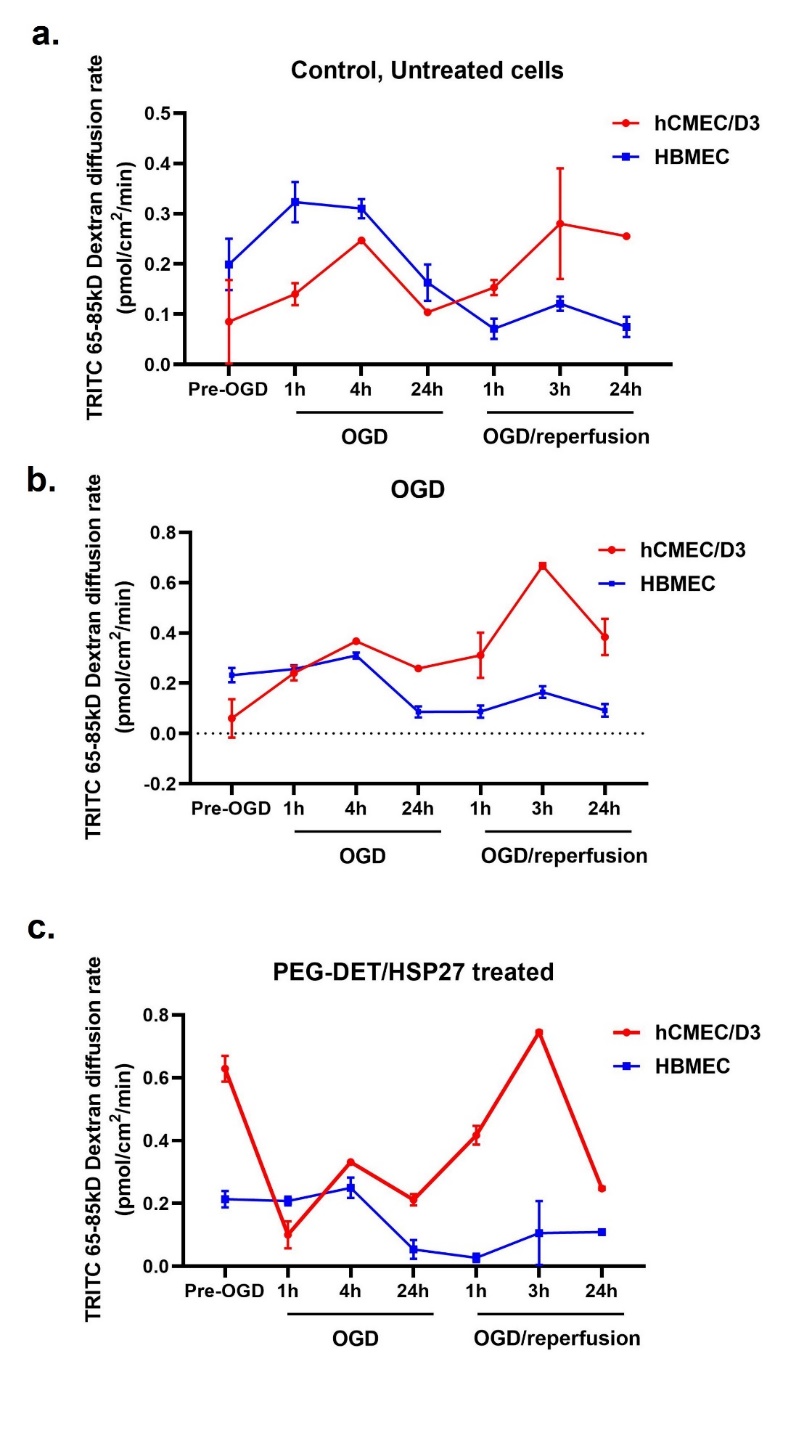

**Fig. S27. Comparison of 65-86kD dextran diffusion rates across transwell monolayers of the hCMEC/D3 endothelial cell line vs. primary HBMECs.** The diffusion rate of 65-85 kD dextran was measured in normoxic cells (**a**), OGD- cells (**b**), and PEG-DET/HSP27-treated cells (**c**) during pre-OGD, OGD for 24 h, and OGD/reperfusion for 24 h conditions. For Pre-OGD conditions, diffusion rate, control, OGD control, and PEG-DET/HSP27 treated cells were incubated with a complete growth medium containing 1µM 65-85 kD TRITC-Dextran for 1 h. For the OGD group, the medium was replaced with an OGD medium containing 1 µM 65-85 kD TRITC-Dextran, and the diffusion rate was measured at 1, 4, and 24 h in the abluminal chamber. Post-OGD, the medium was removed, cells were washed, and treated with a complete growth medium containing 1 µM 65-85 kD TRITC-Dextran. During the OGD/reperfusion phase, the dextran diffusion rate was measured at 1, 3, and 24 h in the abluminal chamber.

**Discussion on leakiness of hCMEC/D3 BEC vs. HBMECs monolayers**

The diffusion rate of the larger 65-85 kD TRITC-Dextran in untreated hCMEC/D3 cells and HBMECs were compared at pre-OGD conditions, during OGD, and OGD/reperfusion conditions (**Fig. S27**). The baseline diffusion rate was found to be different for both cell models. The diffusion rate gradually increased and was relatively higher for the hCMEC/D3 cell line compared to the HBMEC monolayer during ischemia/reperfusion suggesting the inherent leakiness of the hCMEC/D3 cell line compared to primary HBMECs. Therefore, we studied the effects of naïve EV and HSP27 mixtures on the diffusion rate of 4.4 kD and 65-85 kD TRITC-dextran during OGD and OGD/reperfusion conditions in primary HBMEC monolayers. Naïve EVs, PEG-DET/HSP27, and EV/HSP27 mixtures were cytocompatible with HBMECs for 72 h of exposure under normoxic conditions (**Fig. S23**).
